# A patient derived xenograft repository capturing clinical and molecular heterogeneity of large B-cell lymphoma

**DOI:** 10.64898/2026.01.19.700406

**Authors:** Haopeng Yang, Kotaro Arita, Kevin Bowman, Dai Chihara, Jared Henderson, Griffin Rost, Estela Rojas, Sydney Parsons, Priya Lakra, Aneela Abedin, Sattva S. Neelapu, Paolo Strati, Loretta J. Nastoupil, Luis Fayad, Swaminathan P. Iyer, Alma Rodriguez, Frederick B. Hagemeister, Luis Malpica, Hun Ju Lee, Laura Hilton, David W. Scott, Richard Eric Davis, Christopher R. Flowers, Jason R. Westin, Giorgio Inghirami, Francisco Vega, Michael R. Green

**Affiliations:** Department of Lymphoma & Myeloma, University of Texas MD Anderson Cancer Center, Houston, TX, USA; University of Texas MD Ander Cancer Center UT Health Houston Graduate School of Biological Sciences, Houston, TX, USA; Centre for Lymphoid Cancer, BC Cancer, Vancouver, BC, Canada; Department of Pathology and Laboratory Medicine, New York Presbyterian Hospital, Weill Cornell Medicine, New York, NY 10065, USA; Department of Hematopathology, University of Texas MD Anderson Cancer Center, Houston, TX, USA; Department of Hematopoietic Biology and Malignancy, University of Texas MD Anderson Cancer Center, Houston, TX, USA

## Abstract

Large B-cell lymphomas (LBCLs) are a clinically and molecularly diverse group of malignancies with a rapidly evolving therapeutic landscape that has introduced new areas of clinical need such as post-CD19 chimeric antigen receptor T (CART19) progression. Patient derived xenograft (PDX) models are an important tool for mechanistic studies and preclinical evaluation of new therapies and can be generated from a variety of clinical contexts that capture tumor-intrinsic resistance mechanisms. We therefore undertook a comprehensive effort to generate PDX models that encompass the molecular landscape of LBCLs and include important clinical scenarios for new drug development. Here we describe the first 48 models within this publicly available repository, capturing the transcriptional and genetic subsets of LBCL. These models also include 23 generated from post-CART progression biopsies which reproduce patterns of progression driven by *CD19* mutation or expression loss, as well as tumor cell-intrinsic CART19 resistance that we validate *in vivo*.

**STATEMENT OF SIGNIFICANCE:** Yang *et al.* describe X-LYMPH (<u>X</u>enografts of <u>Lymph</u>oma), a publicly available and molecularly annotated PDX repository that captures the heterogeneity of large B-cell lymphoma. X-LYMPH includes models of chimeric antigen receptor T cell resistance, providing a shared foundation for mechanistic research and therapeutic development for lymphomas.

## INTRODUCTION

Large B-cell lymphomas (LBCLs) are an aggressive but clinically and molecularly heterogeneous group of malignancies. Although the majority of cases can be cured with first line chemo-immunotherapy regimens(1), relapsed or refractory cases (rrLBCL) are difficult to manage, highlighting a need for model systems to study resistance mechanisms and test new therapeutics. To date, cell lines have been the predominant model system for the study of LBCL. However, the most employed cell lines were developed in the pre-rituximab era, largely from effusions or leukemic-phase disease, and thus do not fully capture disease biology and/or resistance mechanisms to modern therapies. Although resistance to small molecules can be induced by chronic low-dose exposure to drug(2), there is no clear path towards modeling resistance to cell therapy *in vitro*. Mechanistic and preclinical studies within the context of cell therapy resistance therefore currently rely on the development of models from biopsies taken following cell therapy failure for which resistance was developed *in vivo*. However, such model systems have not yet been described or made available for academic research.

Patient-derived xenograft (PDX) models can be generated by implanting patient tumor biopsy material into immunodeficient mice, with a higher success rate for growth than *in vitro* culture. Some efforts have been made towards generating and characterizing LBCL PDX models(3,4), demonstrating fidelity to the originating patient biopsy(4–6). However, these pre-date the CD19 chimeric antigen receptor T cell (CART19) era, the number remains too small to capture the molecular diversity of LBCL, and not all are openly available to investigators outside of the originating center. We therefore undertook an initiative to generate a large repository of LBCL PDX models, including from relapses occurring following CART19, to provide an academic resource to advance pre-clinical and mechanistic studies of LBCL. Here we describe our experience in this initiative, detail the first cohort of 48 models and show evidence of their utility as models of CART19 resistance.

## RESULTS

### Generation of PDX models of high-risk LBCL

We generated PDX models from core needle biopsies of LBCL patients by renal capsule implantation in NOD.Cg-*Prkdc^scid^ Il2rg^tm1Wjl^/SzJ* (NSG) mice and monitoring the mice for up to 6 months. Tumors that grew (F1) were passaged fresh into additional NSG mice to generate F2 tumors that were extensively characterized, and seeds cryopreserved for future use (**Figure 1A**). Using pathology reports from synchronous cores that were clinically evaluated for diagnostic purposes, we determined that 133 of the implanted biopsies were LBCL in origin, including 98 diffuse large B-cell lymphoma (DLBCL), 19 high grade B-cell lymphoma (HGBL), 6 primary mediastinal large B-cell lymphoma (PMBL) and 10 other LBCL subtypes. HGBLs had the highest success rate for PDX generation (53%), with other histologies having similar take rates (30-34%), resulting in 48 new LBCL PDXs from 45 patients including 3 pairs of PDX biopsies from the same patient (**Figure 1B**). Follicular lymphoma (FL) grade 1-3A biopsies (n=35) generated no successful PDX models. Over a period of ∼6 months, 55 biopsies with a suspicion of lymphoma were implanted the same day of the biopsy procedure without cryopreservation. All other biopsies were viably cryopreserved and thawed for implantation. We evaluated the effect of implant timing on PDX success rate and noted a slight decline with cryopreservation, and additional decline if the biopsy was cryopreserved for greater than 30 days (**Figure 1C**). However, fresh implants were poorly selective for LBCL (**Figure 1D**), whereas viable cryopreservation allowed the consultation of hematopathology reports from synchronous cores to select biopsies for implantation. PDXs tended to be from biopsies taken at later lines of therapy (median=3, range=0-10) (**Figure 1E**). Tissue microarrays were constructed from F2 tumors of each model to facilitate IHC staining. Comparison to available clinical work-up of synchronous cores showed strong concordance for the majority of proteins when comparing dichotomous (positive/negative) classifications (**Figure 1F**). However, heterogeneous patterns of expression were also observed across proteins and samples in tumors and their corresponding PDX (**Figure 1G**). Histology for the 48 new models are shown in **Figure 1H**, summaries of each model provided in supplementary materials, and clinical features in **Table S1**. LBCL PDX models can therefore be generated with high efficiency via renal capsule implantation into NSG mice, but it is recommended to use prior viable cryopreservation (<30 days) to target implantation towards subtypes of interest to optimize resource utilization.

**Figure 1:**
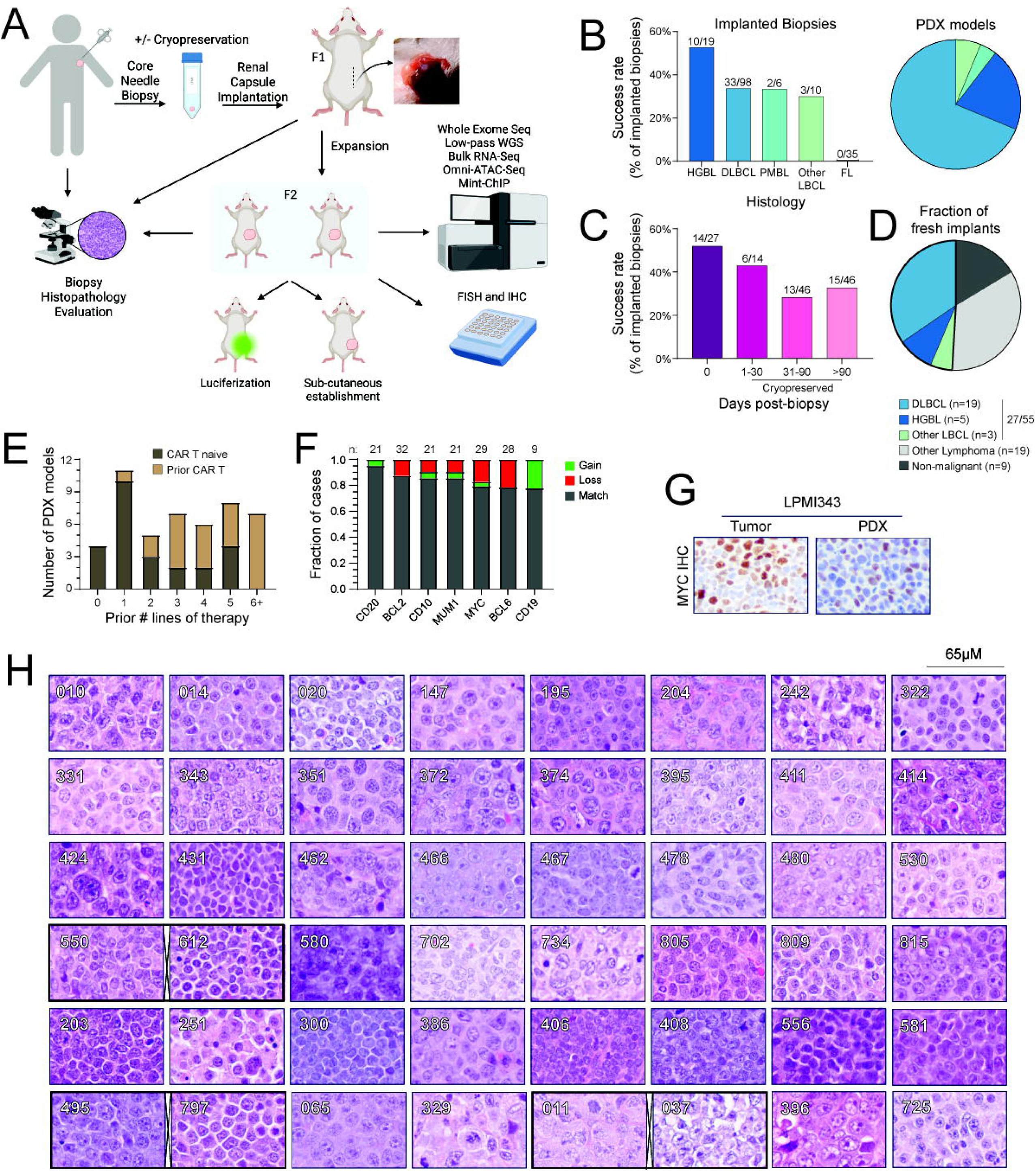
Generation of LBCL PDX models. **A**) A schematic of the workflow used for generating and characterizing PDX models. **B-C**) Success rates of implanted biopsies according to LBCL histology (B) and timing of implantation (C). **D**) Pie chart of the histologies of biopsies implanted without prior cryopreservation. **E**) Number of lines of prior therapy for successful PDX models. **F**) Agreement between IHC of PDX models and clinical workup of synchronous cores. Gain or loss indicates gain or loss of expression in the PDX model relative to the synchronous biopsy. **G**) Example of heterogenous MYC expression in a PDX model and synchronous biopsy. **H**) H&E staining of 48 newly generated PDX models. Dark boxes indicate pairs of models from the same patient.

### PDX models span the molecular landscape of LBCL

Molecular characteristics of PDXs were evaluated by whole exome sequencing (WES), with germline control when available (25/48), low-pass whole genome sequencing (lpWGS) and transcriptional subtyping with the DLBCL90 assay(7) (**Figure 2A; Table S2**). For reference, we obtained and profiled LBCL PDX models from PRoXe and WCM for a total of 80 models. Tumor material was not available for the implanted biopsies because the entire core was engrafted in mice. However, DLBCL is known to be genetically stable(8) thus we compared mutation and LymphGen subtype frequencies to an independent cohort of rrLBCL tumors (n=80)(9). DLBCL90 analysis of the 48 newly generated PDXs identified 4 with PMBL molecular subtype including 2 PMBLs and 2 grey zone lymphomas, 36 activated B-cell-like (ABC) subtype, 30 germinal center B-cell-like (GCB) subtype and 10 unclassified. Of the 30 GCB subtype, 15 were positive for DZsig (**Figure 2A**). WES analysis identified recurrent mutations in genes with well-established roles in DLBCL(10–12) and GISTIC analysis identified significant gains and losses that have been previously described in DLBCL(11,13) (**Figure 2A**; **Figure S1**). However, high genetic complexity led to ambiguous classifications of many models by LymphGen (**Figures 2A-B**), a subtyping system that was generated using newly diagnosed DLBCL. Multiple LymphGen-informing mutations were enriched in PDXs compared to the comparator cohort (**Figure 2C**), including EZB subtype genes such as *CREBBP* and *KMT2D*, though the frequencies of EZB classified samples was similar between cohorts (**Figure 2B**). BN2-informing genes, *NOTCH2* and *SPEN*, were also mutated at higher frequencies in PDXs resulting in a higher frequency of BN2 classified PDXs compared to the comparator cohort (**Figures 2B-C**). However, *NOTCH2* mutations were not restricted to the canonical pattern of N-terminal truncating events that are seen in newly diagnosed DLBCLs of BN2 subtype (**Figure 2D**)(14), highlighting the importance of interpreting data according to mutation type rather than presence or absence. Mutations in *TP53* were present in 42% of PDXs and complex DNA copy number profiles were common (**Figure 2E**), though the A53 LymphGen subtype was called in a lower frequency of PDXs compared to the comparator cohort. Mutations in *MYD88* and *CD79B*, and MCD classified tumors were also modestly under-represented in PDXs, but this effort generated two new models with combined *MYD88*-L265P and *CD79B*-Y197 hotspot mutations that are characteristic of the MCD subtype (**Figure S2**). LBCL PDX models therefore encompass the molecular spectrum of rrLBCL but have biases in molecular features that may require future targeted efforts towards PDX generation to ensure adequate diversity.

**Figure 2:**
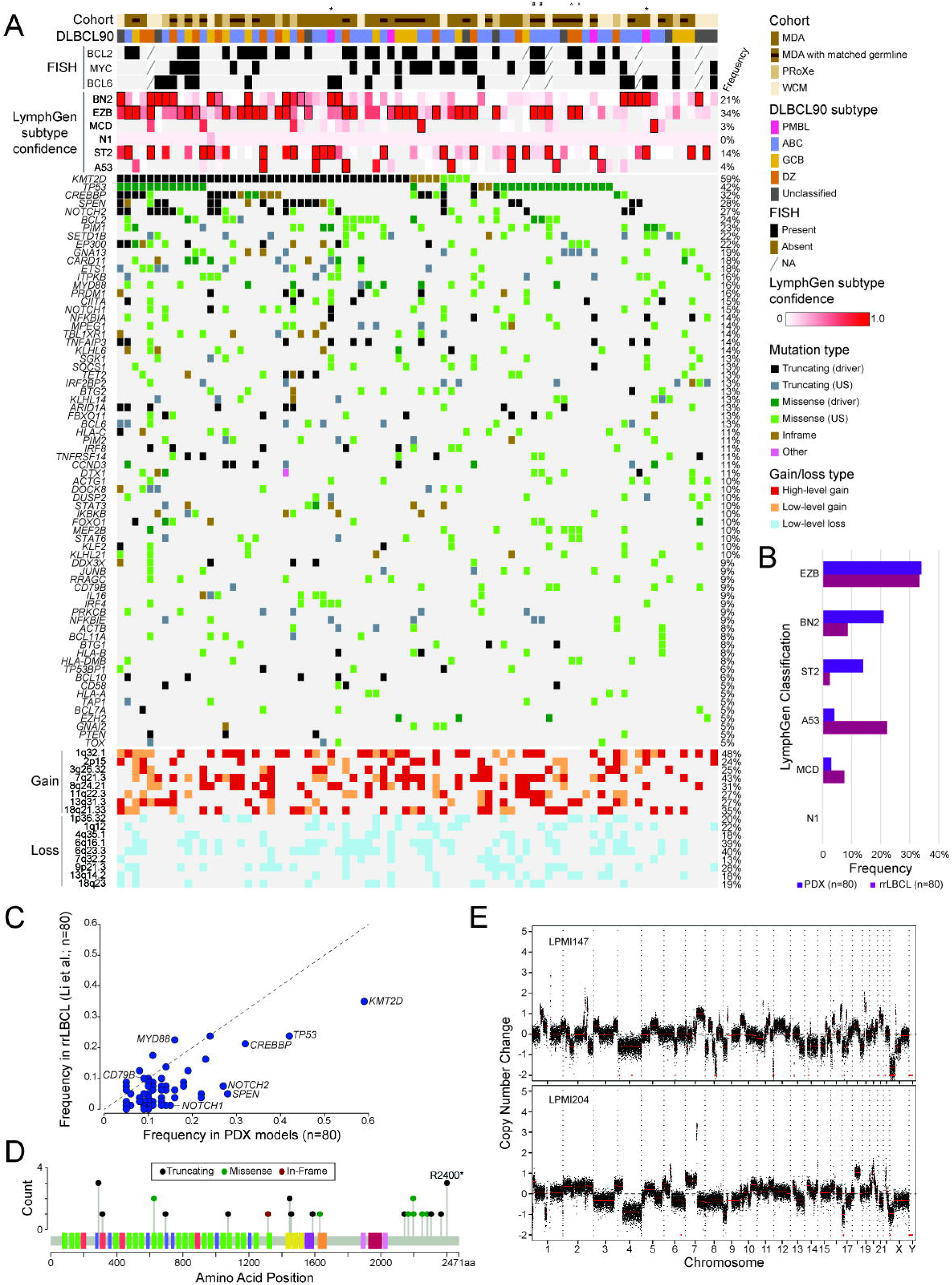
Molecular characteristics of PDX models. **A**) Oncoplot of DLBCL90 transcriptional subtype, FISH, LymphGen classification scores, mutations of individual genes and DNA copy number gains and losses across 80 LBCL PDX models. **B**) Relative frequencies of LymphGen classes in PDX models and independent cohort of rrLBCLs. Samples are counted multiple times if ambiguously assigned to multiple classes. **C**) Relative frequencies of mutations in individual genes in PDX models and an independent cohort of rrLBCLs. **D**) Lollipop plot of *NOTCH2* mutations from PDX models. **E**) Example DNA copy number profiles from lpWGS.

### Models of CART19 resistance

Among the new PDXs, 23 were generated from biopsies obtained after CART19 failure, including 19 with CART19 as the immediate prior line of therapy (**Figure 3A**; **Table S1**). In addition, 11 PDXs from 10 patients were generated from biopsies obtained prior to receiving CART19, of which 4 patients progressed following CART19 and 6 patients were in complete response at last follow-up (**Figure 3A; Table S1**). These models included 3 pairs of PDXs from the same patient. LPMI011 was generated from a biopsy taken following a partial response to CART19 and LPMI037 from the same patient after a second CART19 administration with progressive disease as best response. LPMI495 and LPMI797 were generated from biopsies 13 months apart with 5 intervening lines of therapy including CART19. LPMI550 and LPMI612 generated from biopsies taken 3 months apart with 2 intervening lines of therapy, with the patient subsequently achieving a complete response to CART19 and remaining in remission at last follow-up.

**Figure 3:**
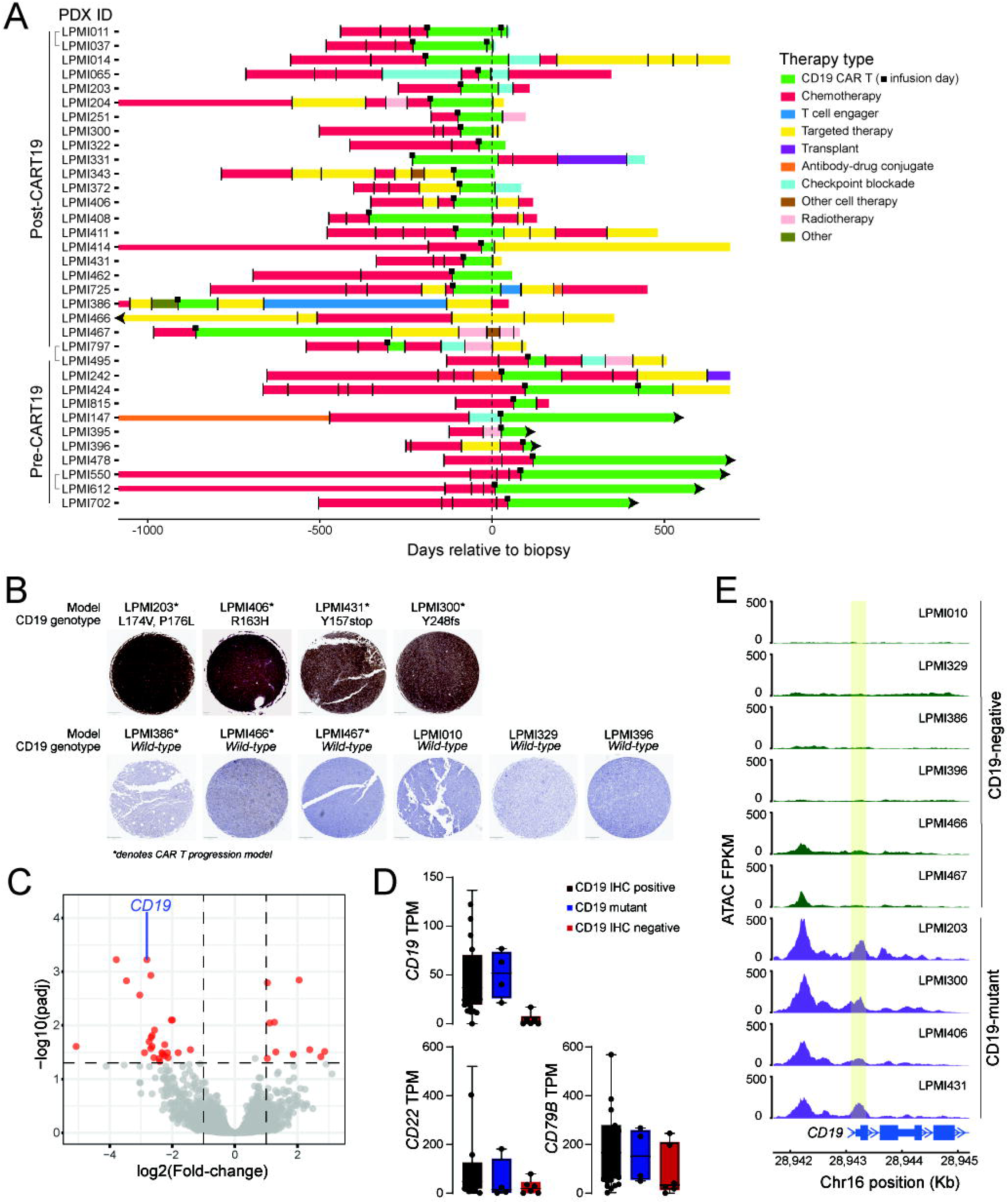
CART19 progression PDX models and CD19 mutation/loss. **A**) Swimmer plot showing lines of therapy, centered on the day of the biopsy that generated the PDX model. **B**) Examples of CD19 IHC in mutant (above) and negative (below) cases. **C**) Volcano plot of differentially expressed genes between CD19 IHC-negative and IHC-positive PDX models showing selective down-regulation of CD19. **D**) Box plots showing CD19, CD22 and CD79B expression according to CD19 mutation and expression status. **E**) Example tracks of chromatin accessibility over the CD19 gene locus showing cases from panel B.

CD19 antigen escape has been described as a mechanism of resistance to CART19(15). WES analysis identified 4 PDXs carrying *CD19* mutations, all generated from post-CART19 progression biopsies. Two possessed monoallelic truncating mutations that reduced CD19 protein expression and two had point mutations within the loop recognized by FMC63, including mutation of arginine 163 that has been previously shown to abrogate CAR binding(16) (**Figure 3B**). IHC also revealed loss of CD19 expression in 6 PDXs, of which 3 were CART19 naïve and 3 from post-CART19 progression (**Figure 3B**). Differential analysis of RNA-sequencing of CD19-negative (n=6) compared to CD19-positive PDX models (n=74) showed significant loss of *CD19* transcript without other broad accompanying signatures or loss of known *CD19* regulators such as *PAX5*(17) (**Figure 3C; Table S3**), suggestive of selective silencing of *CD19* expression. This was supported by ATAC sequencing data that showed a loss of chromatin accessibility at the *CD19* locus in CD19-negative cases but not in CD19-positive cases, indicative of epigenetic silencing of CD19 expression (**Figure 3D**). Notably, comparison of CD19 IHC from patient biopsies with the corresponding PDX model identified 2 instances in which CD19 was negative in the patient biopsy and positive in the PDX model, supporting the notion that CD19 expression is epigenetically regulated and may be subject to restoration. Furthermore, we did not observe loss of CD19 expression in PDX models when the patient biopsy was positive suggesting that this is not a PDX-specific phenomenon. Importantly, *CD19* mutant and protein-negative PDX models maintained variable levels of expression of other CAR targets such as *CD22* and *CD79B* (**Figure 3E**), establishing these as potential systems in which to test the efficacy of alternative CAR designs.

### Tumor cell-intrinsic resistance to CART19

The majority of CART19 progressions retain CD19 expression, highlighting tumor-intrinsic (18) or lymphoma microenvironment (LME)(9) factors in CART19 resistance. PDXs in immunodeficient mice do not maintain their LME and are therefore not a tractable system for evaluating LME-driven CART19 resistance. However, we aimed to investigate whether post-CART19 progression PDXs possessed tumor cell-intrinsic resistance. We generated T cell receptor knock-out (TCR-KO) CART19 cells from 3 healthy donors to avoid the confounding effect of alloreactivity (**Figure 4A**) and confirmed their comparable activity to TCR-intact CART19 cells *in vitro* (**Figure 4B**). Tumors from 7 PDX models, 1 CART19-naïve and 6 post-CART19 progression, were treated with 2×10^6^ CAR-positive TCR-KO CART19 cells (4 mice per donor, 12 per PDX model), half of the mice were euthanized 14days after CART19 infusion and the tumors evaluated for T cell infiltration by flow cytometry and IHC, and the remaining mice monitored by weekly MRI for 4 weeks or until moribund (**Figure 4C**). The CART19-naïve model (LPMI020) showed an increase in size at 1 week post treatment, but with a notable influx of T cells, and responded thereafter with a reduction in tumor size at week 2 and clearance by week 4 (**Figure 4D-E**). The remaining 6 models derived from CART19 progression did not clear the tumor following CART19 treatment but showed different patterns of T cell infiltration and response. Three models showed T cell infiltration within the body of the tumor and a modest reduction in tumor size (LPMI331) or slowing of tumor growth (LPMI462, LPMI204). One model (LPMI408) displayed extensive T cell infiltration in the periphery of the tumor, but with a failure to infiltrate the core, and had rapid progression. The remaining two models (LPMI406, LPMI431) showed very few detectable T cells within the tumor and had rapid progression. These LBCL PDXs therefore capture important clinical scenarios such as post-CART19 progression and retain CART19 resistance *in vivo*, providing experimental systems in which to test therapeutics in this area of high clinical need.

**Figure 4:**
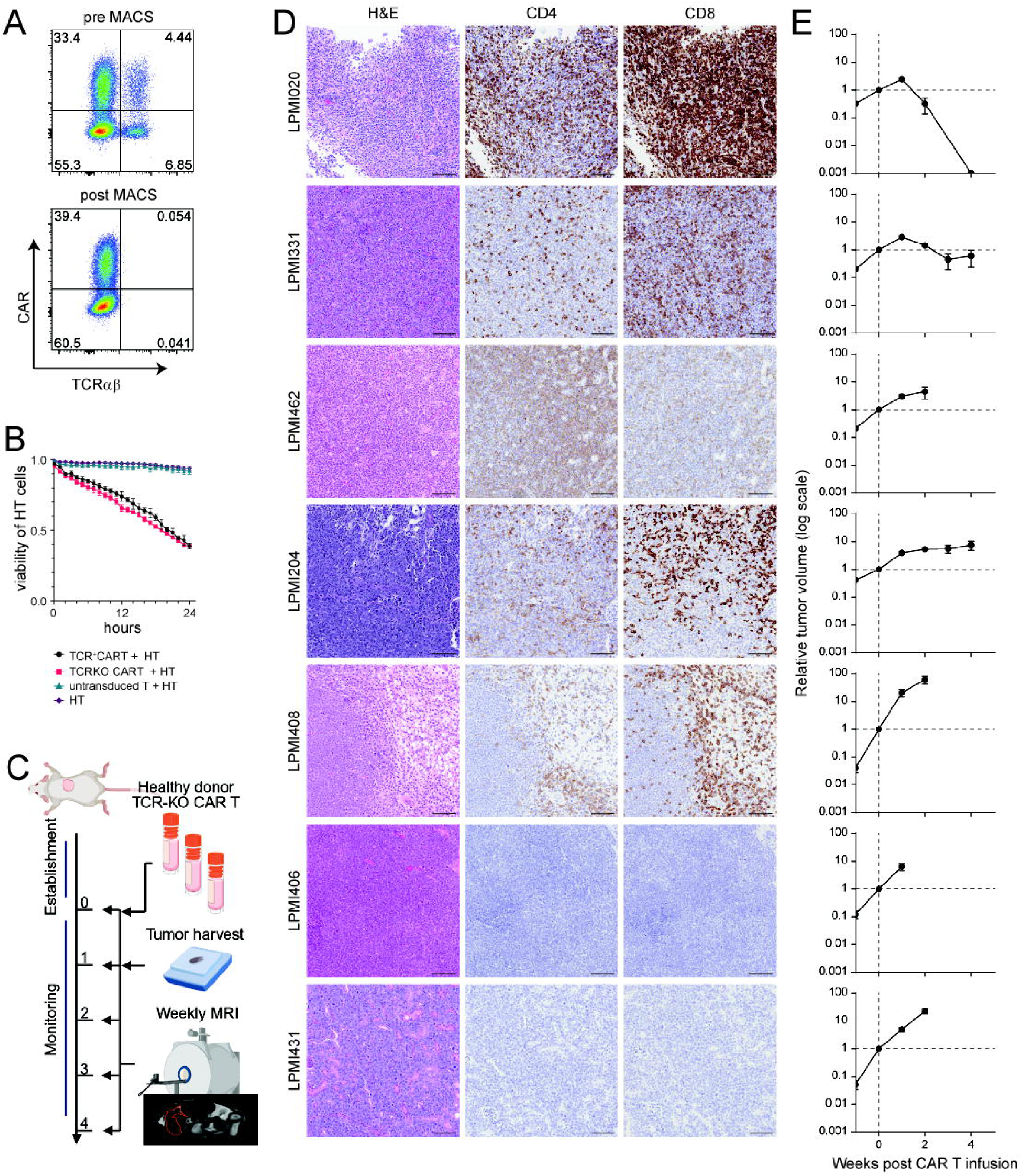
Tumor cell intrinsic CART19 resistance. **A**) Flow cytometry plots showing CAR and TCR positivity following CRISPR knock-out and CAR infection, pre- and post-purification by magnetic beads. **B**) Incucyte analysis of cytotoxicity against the HT cell line comparing TCR intact and TCR-KO CART19 cells from healthy donors. **C**) Schematic for the approach taken to compare CART19 sensitivity of PDX models. **D**) H&E staining and IHC for human CD4 and CD8 in exemplar tumors for each PDX model harvested at day 14 following CART19 treatment. **E**) Tumor volume measured by MRI, relative to the volume on the day of CART19 administration (day 0).

## DISCUSSION

The last decade has seen the approval of multiple new therapies for LBCL, which has introduced new areas of clinical need. Prior efforts to improve patient outcomes in frontline LBCLs have been centered on the ABC transcriptional subtype. However, the approval of polatuzumab vedotin in frontline chemo-immunotherapy combinations for DLBCL improves the outcome of patients with the ABC subtype(19), highlighting “dark zone” or “molecular high grade” subtypes of LBCL as an emerging clinical need in the frontline(20,21). The recent description of genetic subtypes of DLBCL(10–12) has also driven interest in tailoring therapy to genetically-defined subsets of patients(22). However, there remains a paucity of mechanistically rational therapeutic options for molecular subsets of LBCLs. The introduction of CART19 has transformed therapy for rrLBCL(23,24) but patients with post-CART19 progression have dismal outcomes(25). Furthermore, resistance mechanisms to CART19 are largely non-genetic(9,17) and difficult to replicate using cell line models, highlighting the complexity of the underlying biology. New drug development for emerging areas of clinical need will therefore require the proactive development of model systems that capture the molecular complexity of LBCL together with tumor-intrinsic resistance mechanisms that are acquired within patients under the selective pressure of therapy. Here we show that this can be accomplished through prospective generation of PDX models from high-risk patients.

Molecular characterization of our LBCL PDX series shows that it encompasses molecular subsets of disease such as cell of origin and dark-zone transcriptional subtypes and LymphGen genetic subtypes. We note that PDX models have over-representation of mutations in genes such as *TP53*, *KMT2D*, *CREBBP*, *NOTCH2* and *SPEN*, among others. Biases in genetics and other molecular characteristics are to be expected when generating PDX models but must be accounted for when subsequently selecting models for mechanistic or preclinical studies. Furthermore, we show that under-represented subsets such as the MCD genetic subtype can still be generated, suggesting that targeted efforts towards PDX generation could be utilized to fill in gaps of under-represented characteristics.

We show that tumor-intrinsic characteristics such as CART19-resistance is retained in PDX models. Furthermore, we noted different patterns of progression following CART19 treatment *in vivo*, with variable degrees of T cell infiltration and response, highlighting these PDXs as important experimental systems to investigate underlying mechanisms of CART19 resistance and/or to test therapies for this clinically high-need subset of patients. However, PDXs have multiple important limitations that need to be considered. First, their use is resource and time intensive, with *in vivo* experiments commonly taking up to 6 months each. This can be mitigated by the generation of luciferized models that can be implanted subcutaneously, alleviating the need for complex surgeries and imaging, but vascularization and other anatomical differences between sites must be careful considered. Second, our PDX models are developed in immunodeficient mice that lack the immune contexture that is central to LBCL biology. Future advancements in humanization of immune deficient mice might facilitate the reproduction of immune-intact microenvironments, but these will be hampered by the need for HLA matching and rigorous validation of cell-cell communication patterns compared to primary tumors. Further work is therefore required for development of *in vivo* systems that will be applicable for mechanistic immunology and the development of novel immune therapies.

In conclusion, X-LYMPH provides a publicly available repository of molecularly-annotated LBCL PDX models that include post-CART19 progression. These models provide a unique resource for exploration of disease biology and rigorous preclinical testing of new therapies for high-risk patients.

## METHODS

### Resource Availability

All models are available to not-for-profit academic institutions following a materials transfer agreement. Each model is summarized in the supplementary information. To request models, please

Raw sequencing data will be made available through the European Genome-Phenome Archive (EGA) prior to publication of this manuscript. Processed files have been uploaded to the gene expression omnibus (GEO) under accession number GSE314566.

### Sample Collection

All patients in this study provided informed consent and work was carried out in alignment with the Helsinki declaration. All work was approved by the Internal Review Board of MD Anderson (MDA). Core needle biopsies were collected during routine diagnostic procedures by the MD Anderson Lymphoma Tissue Bank and either viably cryopreserved or used fresh. A complete standard operating procedure for biopsy cryopreservation is available in the supplementary materials.

### Renal Capsule Implantation

A complete standard operating procedure for renal capsule implantation is available in the supplementary materials. Core needle biopsies or previously banked PDX seeds (1.5mm^2^) were implanted into the sub-renal capsule of NOD.Cg-*Prkdc^scid^ Il2rg^tm1Wjl^/SzJ* (NSG) mice. For comparison, we additionally acquired and propagated available models from the Public Repository of Xenografts(3) (PRoXe, n=11), Weill Cornell Medical College (WCM, n=18) and 3 Richters transformation models previously established at MDA. Of the WCM PDX models, 11 were previously established at WCM and 7 were newly established at MDA from WCM specimens.

### Validation of PDX Models

Tumors from F1 and F2 generations were formalin-fixed and paraffin-embedded then evaluated by hematoxylin and eosin (H&E) staining by an expert hematopathologist (F. Vega) to confirm LBCL histomorphology. All models were evaluated for EBV status by EBER in situ hybridization (ISH) and EBV-positive models (n=1) discarded. F2 specimens from all models were assembled onto tissue microarrays (TMAs) and stained for PAX5 to confirm B cell identity as well as other clinically relevant immunostains (CD10, CD19, CD20, BCL2, BCL6, MUM1, MYC). TMAs were also evaluated for rearrangements of the *MYC*, *BCL2* and *BCL6* loci by fluorescent in situ hybridization (FISH). Results are summarized within the supplementary materials. Where available, IHC and FISH results were compared to those obtained from clinical workup of synchronous biopsies.

### Molecular analysis

Whole exome (WES) and low pass whole genome (lpWGS) sequencing was performed on F2 tumors and matched germline DNA when available (MDA models only, n=25) to characterize mutations and DNA copy number alterations, respectively, and for LymphGen classification(10), as previously described(9). The DLBCL90 Nanostring assay was conducted to determine PMBL signature, cell of origin (COO) and dark zone signature (DZsig) status, as previously described(7).

### Generation of TCR-KO CART19 cells

PBMCs from three healthy donors were isolated by Ficoll density centrifugation, cryopreserved, and expanded in CTS OpTmizer–based T cell medium supplemented with IL-2. T cells were activated using CD3/CD28/CD2 stimulation for 3 days, followed by electroporation with Cas9–sgRNA ribonucleoprotein complexes targeting *TRAC*. Immediately after editing, cells were transduced with CD19-28ζ CAR lentivirus by spinoculation. Edited cells were cultured in IL-2–supplemented medium, and TCR-positive cells were depleted by magnetic selection. TCR-KO CART19 cells were expanded, cryopreserved, and assessed by flow cytometry for CAR expression and TCR loss. Functional activity was evaluated in co-culture assays with target cells using Incucyte SX5 (Sartorius). Detailed methods are provided in the Supplementary Materials.

### *In vivo* assessment of CART19 response

All experiments evaluating CART19 responses *in vivo* were conducted using the secondary implantation model. Cryopreserved PDX tumor fragments were implanted in NSG mice to generate seed tumors. After adequate tumor growth, fresh tumor fragments were implanted into the experimental mice. Weekly MRIs on Bruker systems were performed at the MDA small animal imaging facility. Once the median tumor volume reached 50 mm^3^, 2×10^6^ CART19 cells, thawed and rested overnight in CAR T medium with rhIL2, were administered via the tail vein.

### Histological analysis of T cell infiltration

All tumor samples were sectioned along with the long axis of kidney, and one part of them was fixed in neutral formalin. Thereafter, H&E staining and immunohistochemistry for CD4 (clone RBT-CD4, Bio SB) and CD8 (polyclonal, NBP2-29475, Novus Biologicals) were conducted by MDA DVMS core facility. Scanned images were evaluated in Aperio ImageScope (Leica Biosystems).

## ACKNOWLEDGEMENTS

This work was supported by Ed and Beatriz Schweitzer, the MD Anderson B-cell Lymphoma Moonshot Program and NCI P01CA272296 (PI, Cerchietti and Flowers). The MD Anderson Lymphoma Tissue Bank is supported by KW Cares. MRG is a scholar of Blood Cancer United (Formerly Leukemia and Lymphoma Society). HY is a fellow of Blood Cancer United (Formerly Leukemia and Lymphoma Society).

