## Supplementary Methods and Figures for "A patient derived xenograft repository capturing clinical and molecular heterogeneity of large B-cell lymphoma"

<u>Table of Contents</u>


Supplementary Methods

Supplementary Figures

- Figure S1: GISTIC analysis of DNA copy number from 80 cases of LBCL PDX.
- Figure S2: Lollipop plot of *MYD88* and *CD79B* mutations from PDX models.

Supplementary Tables

- Table S1: Clinical features of 48 cases LBCL PDX from MD Anderson.
- Table S2: Molecular characteristics of 48 cases LBCL PDX from MD Anderson.
- Table S3: Differentially expressed genes between CD19-positive and CD19-negative PDX.

Other Supplementary Materials

- PDX model descriptive reports.
- Standard Operating Procedure (SOP) for biopsy cryopreservation.
- Standard Operating Procedure (SOP) for renal capsule implantation.

#### SUPPLEMENTARY METHODS

##### Tissue micro array (TMA) generation

Freshly harvested PDX tumors were fixed in 10% formalin for 24-48 hours, followed by an additional 24-hour incubation in 70% ethanol prior to embedding in paraffin. Formalin-fixed paraffin-embedded (FFPE) tumor blocks were used to generate tissue microarrays. Representative tumor regions were identified and marked on hematoxylin and eosin (H&E) stained slides by a pathologist. Using a tissue microarrayer, cylindrical cores of 1.0 mm diameter were extracted from donor blocks and arrayed into recipient paraffin blocks, with normal human tonsil tissue included as controls. Each case was represented by three cores to capture intratumoral heterogeneity. TMA blocks were sectioned at 4 μm thickness for subsequent immunohistochemistry (IHC), in situ hybridization (ISH), and fluorescence in situ hybridization (FISH) analyses.

##### Immunohistochemistry (IHC)

Immunohistochemistry (IHC) was performed using the Leica BOND™ RX autostainer and BOND™ Polymer Refine Detection kit (Leica Biosciences, DS9800). Antibody conditions are listed as follows: BCL2 (Bcl-2/100/D5, Leica Biosystems, RTU), BCL6 (LN22, Leica Biosystems, RTU), CD10 (56C6, Leica Biosystems, RTU), CD19 (BT51E, Leica Biosystems, RTU), CD20 (L26, Leica Biosystems, RTU), CD30 (BER-H2, Cell Marque, RTU), CD38 (SPC32, Leica Biosystems, 1:75), CD79a (JCB117, Dako, 1:50), c-MYC (Y69, Roche-Ventana, RTU), HLA-DR (EPR3692, Abcam, 1:4000), Ki-67 (MIB-1, Biocare Medical, RTU), MUM-1 (MUM1p, Dako, 1:35), and PAX-5 (BSAP, Dako, RTU). Four-micron-thick formalin-fixed, paraffin-embedded (FFPE) tissue microarrays (TMA; 3 cores of 1 mm diameter per case) and whole tissue slides (Ki-67 only) were used. All incubation steps were performed at room temperature with wash cycles between each step unless otherwise noted. Slides underwent baking and dewaxing, followed by heat-induced epitope retrieval (HIER) using Epitope Retrieval solutions 1 (ER1) and 2 (ER2) for 20 minutes at 100°C. Endogenous peroxidase activity was blocked with hydrogen peroxide for 5 minutes, followed by incubation with the primary antibody for 60 minutes. Slides were then treated with post-primary and polymer reagents for 15 minutes each. Antibody-antigen complexes were visualized using DAB Refine for 10 minutes and counterstained with hematoxylin for 5 minutes. Slides were dehydrated through graded ethanol solutions, cleared in xylene, and coverslipped with Cytoseal™ XYL. All slides were evaluated by standard light microscopy. HLA-DR positivity was defined as >30% of neoplastic cells staining positive. Cell-of-origin (COO) classification was determined using the Hans algorithm (CD10, BCL6, MUM1) with a positivity cutoff of >30%. Double-expressor status was defined as >40% of neoplastic cells positive for c-MYC and >50% positive for BCL2.

**In situ hybridization (ISH)**

In situ hybridization for Epstein-Barr virus-encoded RNA (EBER) was performed on whole tissue sections using a fluorescein-labeled peptide nucleic acid probe (Ventana) with the Ventana peptide nucleic acid ISH Detection Kit for FFPE tissues.

**Fluorescence in situ hybridization (FISH)**

FISH analysis for MYC, BCL2 and/or IGH/BCL2, and BCL6 was performed on TMA FFPE tissue sections using the MYC BAP, BCL6 BAP, BCL2 BAP and/or IGH/BCL2 dual-fusion probes (Abbott Laboratories, Des Plaines, IL, USA) following the manufacturers' instructions. For each probe set, a total of 200 interphase nuclei were analyzed. Abnormal signal patterns were defined using laboratory-established cutoffs for FFPE tissues: 1R1G1F (8.1%), 1G1F (4.6%), 1R1F (5.1%), 1G2F (5.0%), and 1R2F (4.2%).

**Low-pass whole genome sequencing (lpWGS)**

PDX tumors were homogenized, and genomic DNA was extracted using the AllPrep DNA/RNA Kit (Qiagen, 80204) according to the manufacturer's instructions. DNA quantity was measured using the Qubit High Sensitivity dsDNA Kit (Life Technologies, Q32854), and DNA integrity was assessed on a TapeStation 4200 (Agilent, G2991BA). A total of 500 ng of genomic DNA was used for library preparation with the KAPA HyperPlus Kit (Roche, KK8512) using enzymatic fragmentation and 6-8 cycles of PCR amplification. Libraries were purified with AMPure XP beads (Beckman Coulter), validated for size distribution on a TapeStation 4200, quantified by Qubit, multiplexed, and sequenced on an Illumina NovaSeq 6000 platform to achieve a target genome-wide coverage of 1X, using a 100-10-10-100 bp read configuration.

**Whole exome sequencing (WES)**

Whole-exome sequencing was performed using the same genomic DNA libraries generated for low-pass whole-genome sequencing to maintain consistency across assays. For exome capture, equal amounts of each library were multiplexed at a 6-8 sample ratio per capture and processed according to the manufacturer's protocols (IDT or NimbleGen). Captured libraries were subsequently amplified, quantified, and sequenced to achieve a target coverage of 50X for tumor samples, enabling reliable detection of coding mutations, small insertions/deletions, and other functional variants. For the corresponding PBMC samples, the same workflow was used except sequencing depth was adjusted to a target of 30X coverage.

**RNA-sequencing (RNA-seq)**

PDX tumors were homogenized, and total RNA was extracted using the AllPrep DNA/RNA Kit (Qiagen, 80204) and quantified with the Qubit High Sensitivity RNA Kit (Life Technologies, Q32852). One microgram of RNA was used to generate sequencing libraries with the KAPA HyperPrep RNA Kit with RiboErase (Roche, KK8560), using 8 cycles of PCR amplification according to the manufacturer's instructions. Libraries were assessed for fragment size distribution on a TapeStation 4200 (Agilent, G2991BA), quantified with the Qubit High Sensitivity dsDNA Kit (Life Technologies, Q32854), multiplexed, and sequenced on an Illumina NovaSeq 6000 using a 100-10-10-100 bp read configuration.

**ATAC-sequencing (ATAC-seq)**

Viably cryopreserved PDX tumor cells were thawed and processed using the Dead Cell Removal Kit to eliminate non-viable cells. Fifty thousand live tumor cells were washed in ATAC-seq resuspension buffer, resuspended in 50 µL of ATAC-seq resuspension buffer containing 0.1% NP-40, 0.1% Tween-20, and 0.01% digitonin, and incubated on ice for 3 min. Following lysis, nuclei were washed with cold ATAC-seq resuspension buffer containing 0.1% Tween-20 (without NP-40 or digitonin), then resuspended in 50 µL of transposition mixture and incubated at 37 °C for 30 min in a thermomixer at 1000 rpm. Fragmented DNA was purified using the Zymo DNA Clean & Concentrator-5 kit (Zymo, D4024). ATAC-seq libraries were prepared with the KAPA HyperPrep Kit (Roche, KK8502) using 10 cycles of PCR amplification. Libraries were assessed on a TapeStation 4200 (Agilent, G2991BA), quantified with the Qubit High Sensitivity dsDNA Kit (Life Technologies, Q32854), multiplexed, and sequenced on an Illumina NovaSeq 6000 using a 100-10-10-100 bp read configuration.

**lpWGS analysis**

Raw sequencing reads were trimmed using Trimmomatic, and both pre- and post-trimming quality metrics were assessed with FastQC. Trimmed reads were aligned to the hg38 human reference genome using the Burrows-Wheeler Aligner (BWA). Following deduplication, BAM files were processed with XenoFilter to remove contaminating mouse reads and subsequently realigned around insertions/deletions with base quality score recalibration (BQSR) using GATK. The processed BAM files were used to generate copy number segmentation data with CopyWriteR using 100-kb bins, and tumor fractions were estimated with IchorCNA. Copy number alterations identified by IchorCNA were further analyzed using the LymphGen classifier and GISTIC2 to assess recurrent genomic alterations. For visualization, data were represented as log2 copy number changes and segmented using CopyWriteR, and peaks of significant DNA copy number gains and losses were identified using GISTIC2.0.

**WES analysis**

WES data processed using an in-house computational pipeline. Raw sequencing reads were trimmed with Trimmomatic, and both pre- and post-trimming quality metrics were evaluated using FastQC. Trimmed reads were aligned to the hg38 human reference genome using the Burrows-Wheeler Aligner (BWA). Following deduplication, BAM files were processed with XenoFilter to remove contaminating mouse reads and then realigned around insertions and deletions with GATK Base Quality Score Recalibration (BQSR). Coverage metrics were calculated using Picard GATK-CollectHsMetrics, and variants were called using GATK-Mutect2 and functionally annotated with GATK-Funcotator, and high-confidence variants were selected using GATK-SelectVariants. When matched PBMC samples were available, variants were called with GATK-Mutect2 using the PBMC as a germline control.

**RNA-seq analysis**

RNA-seq data were processed using an in-house computational pipeline. Raw reads were trimmed with Trimmomatic, and both pre- and post-trimming quality metrics were assessed using FastQC. Trimmed reads were aligned to the human reference genome (hg38) using STAR in 2-pass mode. To remove contaminating mouse reads, trimmed reads were filtered using XenoFilteR. Only samples with at least 20 million uniquely mapped reads were retained for downstream analyses. Gene-level quantification was performed with HTSeq using the following parameters: -m union -f bam -i gene_id -s reverse -t exon --additional-attr=gene_name. The resulting DESeqDataSet was filtered to retain genes with a minimum of 10 total reads across all samples. Transcripts per million reads (TPM) values were calculated for expression normalization. For gene set enrichment analyses, normalized counts were evaluated using curated B-cell lymphoma gene sets and the SignatureDB database (https://lymphochip.nih.gov/signaturedb/).

#### ATAC-seq analysis

Technical replicates for each ATAC-seq sample were merged prior to quality control assessment with FastQC. Adapter sequences were removed from paired-end FASTQ files through Trimmomatic trimming, followed by alignment to the hg19 human reference genome using the Burrows-Wheeler Aligner (BWA). Post-alignment processing included coordinate-based sorting and filtering to exclude reads mapping to mitochondrial DNA and blacklisted genomic regions via Samtools. PCR duplicates were marked and removed using Picard Tools' MarkDuplicates command. Deduplicated BAM files were then processed with XenoFilteR to remove contaminating mouse reads. The processed BAM files, having completed alignment, sorting, and filtering, served as input for subsequent analyses. For track visualization purposes, BAM files were transformed into BigWig format through deepTools' bamCoverage function, incorporating Tn5 transposase insertion site correction. Peak calling was performed on Tn5-adjusted insertion sites using MACS2 with the following parameters: -g hs (genome size), -q 0.05 (q-value threshold), -f BAMPE (paired-end BAM format), and -B (bedGraph output). Genomic tracks were visualized using pyGenomeTracks version 3.9.

#### Generation of TCR knock-out (TCR-KO) CD19 CAR T cells

Peripheral blood mononuclear cells (PBMCs) obtained from three healthy donors (55 F, 53 F, and 40 M, respectively; Gulf Coast Regional Blood Center, Houston, TX) were isolated using Ficoll-Paque (Cytiva) and cryopreserved in FBS with 10% DMSO  until use. Thawed PBMCs were resuspended in CTS OpTmizer T cell expansion SFM supplemented with CTS Immune Cell SR, and GlutaMAX (all from Gibco; hereafter referred to as CAR T cell medium) at 1×$10^6$ cells/mL, and were stimulated with 25 µL/mL ImmunoCult Human CD3/CD28/CD2 T cell activator (STEMCELL) and 200 IU/mL recombinant human interleukin-2 (rhIL2; GenScript) for 3 days (Day 0). On day 3, activated T cells were resuspended in Buffer T (Thermo) at 4×$10^6$ cells/100 µL, and ribonucleoproteins (RNPs) that consisted of 200 pmol *Streptococcus pyogenes* Cas9 and 500 pmol sgRNA targeting *TRAC* (both from Synthego) were electroporated into cells using 100 µL Neon reaction tips (Invitrogen) with 1400 V, 10 ms, 3 pulses. Cells were subsequently plated in OptiMEM (Gibco) containing 25% v/v CD19 CAR lentiviral supernatant with 10 µg/mL Vectofusin-1 (Miltenyi Biotec) at 1×$10^6$ cells/mL and spinoculated at 400×g for 2 hours. On day 4, cells were resuspended in CAR T cell medium supplemented with 200 IU/mL rhIL2. On day 6, TCR-positive cells were depleted using a biotin-conjugated anti-human TCRα/ß antibody (clone IP26, BioLegend) and MACS systems (Miltenyi Biotec). Then, the retrieved TCR KO cells were expanded in G-Rex 6M plates (Wilson Wolf)by day 10, and cryopreserved until use. CAR positivity and TCR KO rate were evaluated by flow cytometry using VioBright B515 anti-CD19 CAR FMC 63 idiotype antibody (clone REA1297, Miltenyi Biotec) and Alexa Fluor 647 anti-TCRα/ß antibody (clone IP26, BioLegend) together with ViaDye Red (Cytek). Co-culture assay with HT cells (ATCC) and CAR T cells at 1:1 E:T ratio was conducted using Incucyte SX5 (Sartorius) according to the manufacturer instructions with minor modifications.

For lentiviral supernatant production, CD19-28z CAR plasmid and 3rd generation packaging mix (abm) were co-transfected into HEK293T cells (ATCC) using Lipofectamine 3000 (Invitrogen) according to the manufacturer's protocol with minor modifications. Viral supernatant was harvested 2 days after transfection, filtered through a 0.45 µm PVDF membrane (Millipore), and cryopreserved at -80°C until use.

#### SUPPLEMENTARY FIGURES

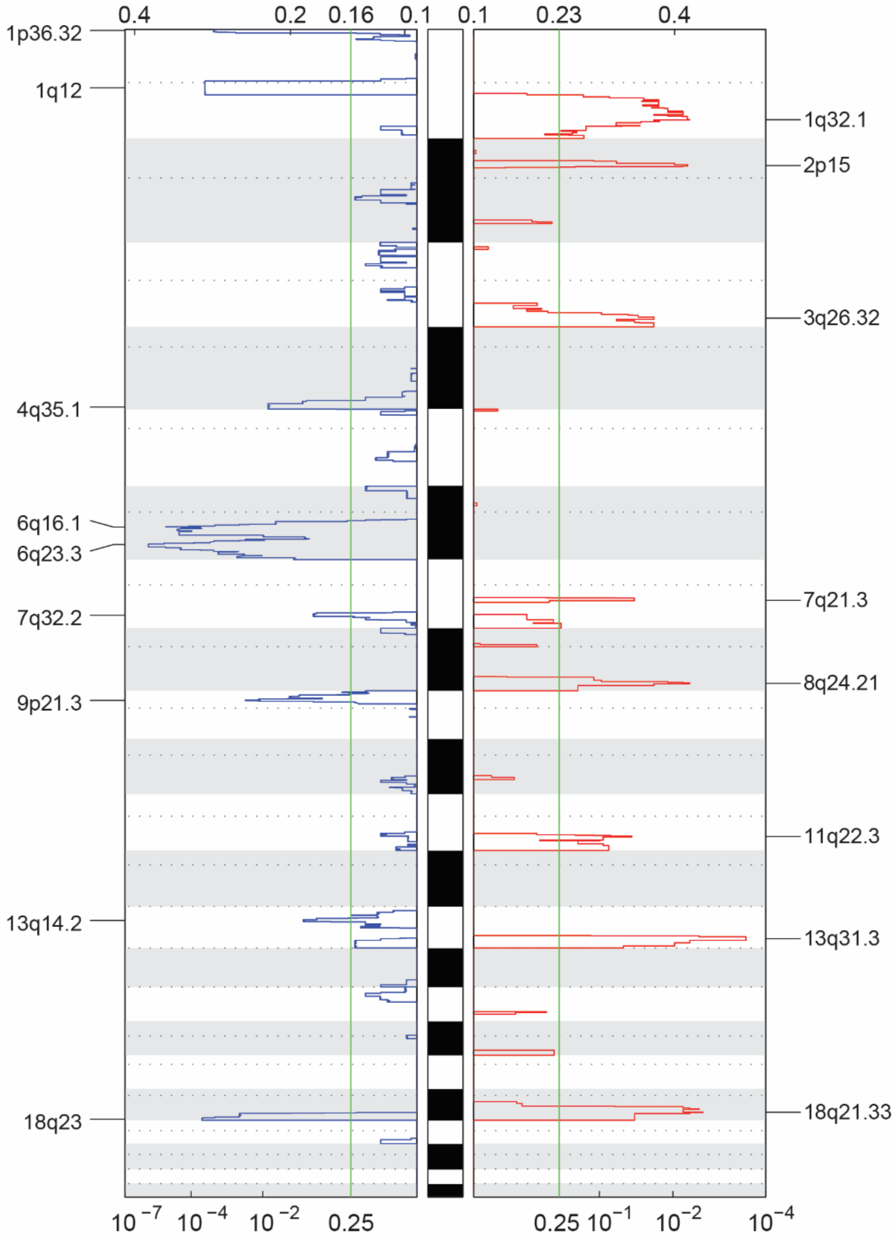


**Figure S1: GISTIC analysis of DNA copy number from 80 cases of LBCL PDX.**

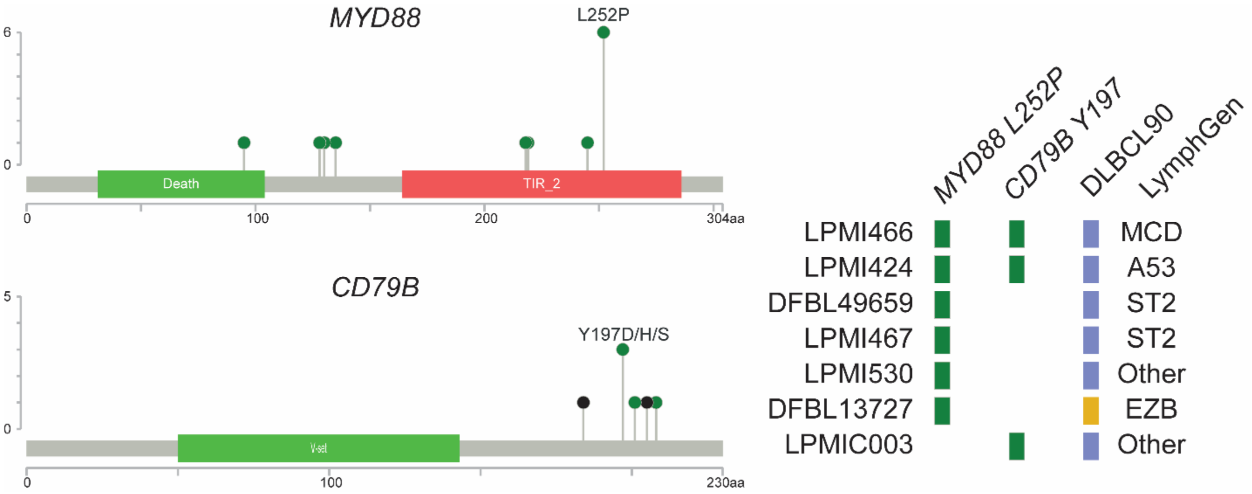


**Figure S2: Lollipop plot of *MYD88* and *CD79B* mutations from PDX models.**

### LPMI010

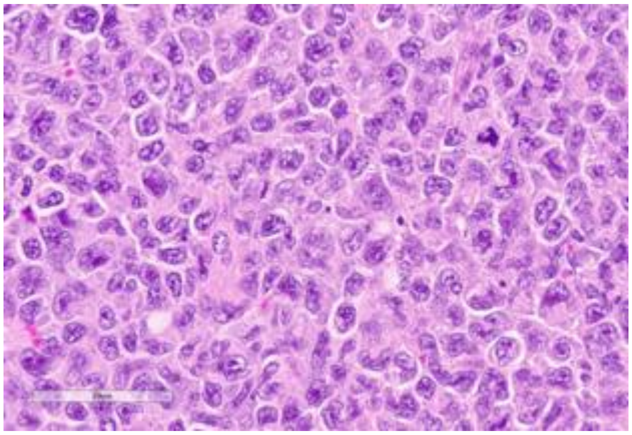

Source: MDACC

**Clinical Information**

Originating Biopsy Diagnosis: DLBCL, NOS

Prior Indolent Lymphoma: None

Sex: Male

Race: White or Caucasian

Ethnicity: Not Hispanic or Latino

Age at diagnosis: 22

Age at PDX biopsy: 23

### LoTs Pre Biopsy (best response): 1 (PD)

### LoTs Post Biopsy (best response): 1 (NA)

Therapies: Chemotherapy; Chemotherapy

**Immunohistochemical Staining:**

Hans COO call: Non-GCB

BCL6 (-)

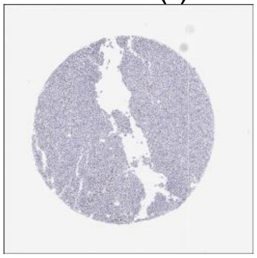

CD10 (-)

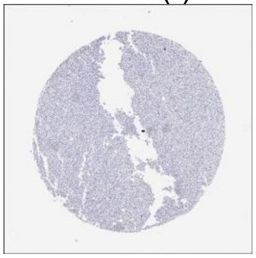

MUM1 (+)

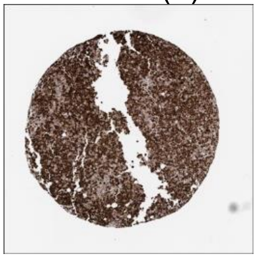

MYC (-)

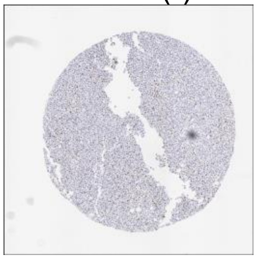

CD20 (-)

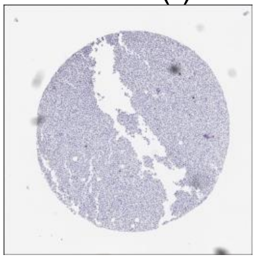

CD19 (-)

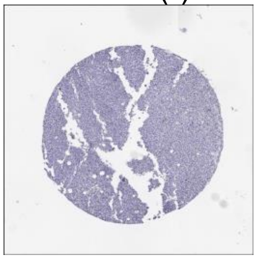

**DLBCL90 call:** ABC

**Genetic Characteristics:**

BCL2 FISH: Negative | MYC FISH: Negative | BCL6 FISH: Negative

Copy Number Alterations:

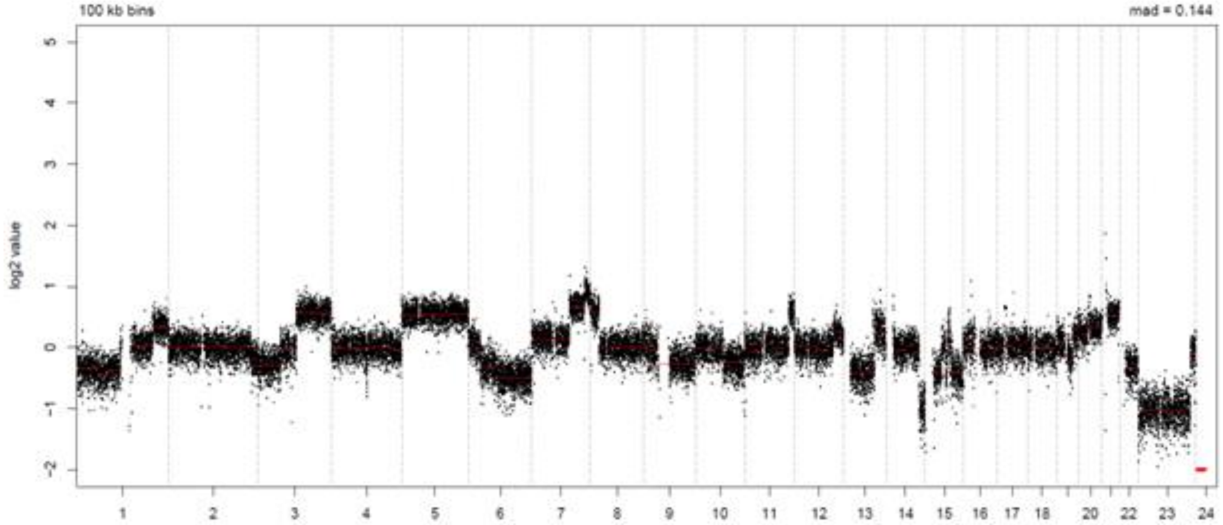


LymphGen Class (Confidence): ST2/A53

BN2: 0.22
EZB: 0.04
N1: 0.07
MCD: 0.00
ST2: 0.99
A53: 0.99

### LPMI011

Source: MDACC

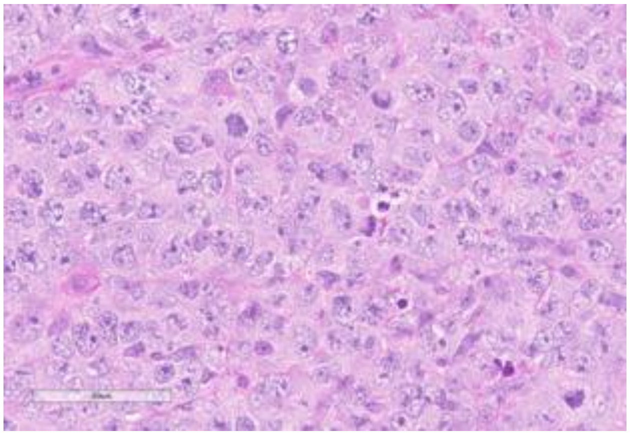

**Clinical Information**

Originating Biopsy Diagnosis: PMBCL

Prior Indolent Lymphoma: None

Sex: Male

Race: Black or African American

Ethnicity: Hispanic or Latino

Age at diagnosis: 37

Age at PDX biopsy: 39

### LoTs Pre Biopsy (best response): 4 (PR)

### LoTs Post Biopsy (best response): 2 (PD)

Therapies: Chemotherapy; Chemotherapy; Chemotherapy; CART; CART; Immunotherapy

**Immunohistochemical Staining:**

Hans COO call: GCB

BCL6 (+)

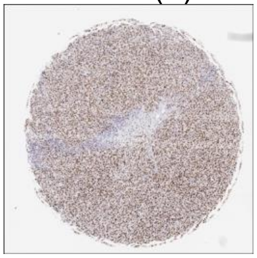

CD10 (+)

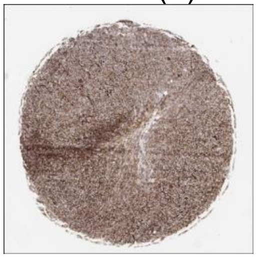

MUM1 (+)

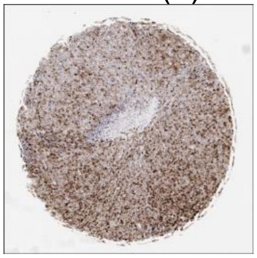

MYC (+)

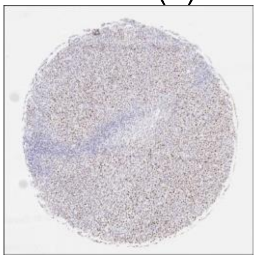

CD20 (+)

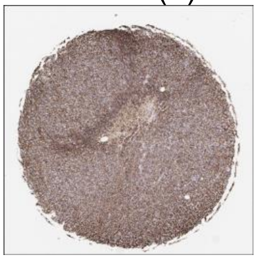

CD19 (+)

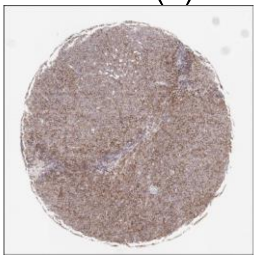

**DLBCL90 call:** PMBL

**Genetic Characteristics:**

BCL2 FISH: Negative | MYC FISH: Negative | BCL6 FISH: Positive

Copy Number Alterations:

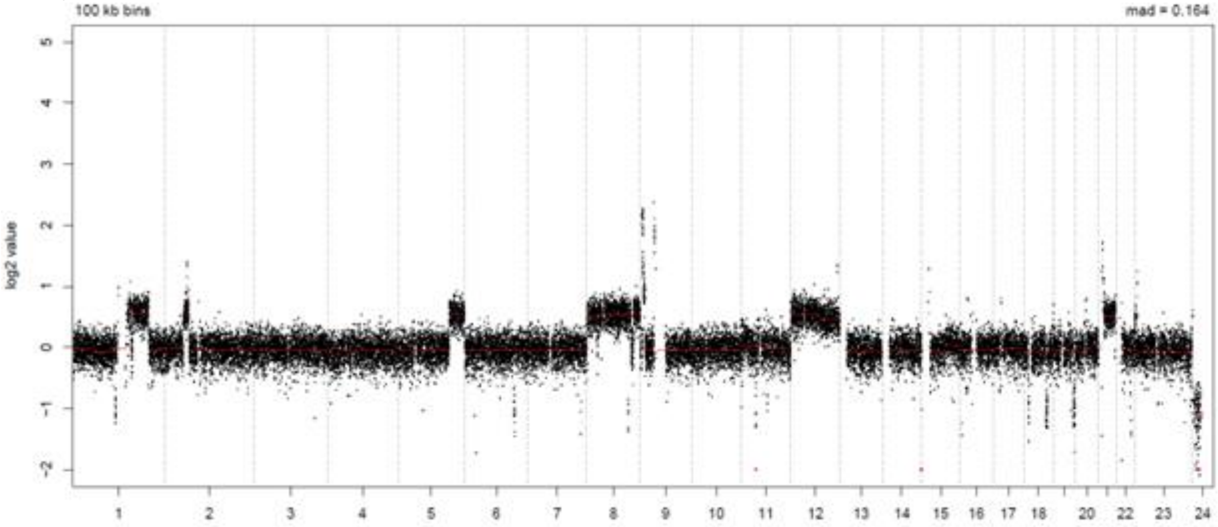


LymphGen Class (Confidence): BN2/ST2

BN2: 0.91 | EZB: 0.04 | N1: 0.07
MCD: 0.06 | ST2: 0.99 | A53: 0.00

### LPMI014

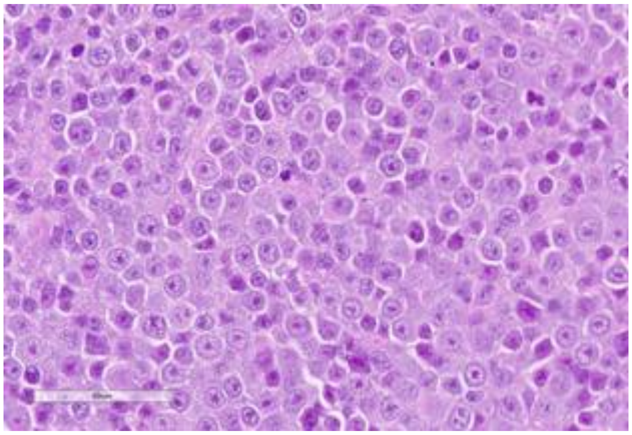

Source: MDACC

**Clinical Information**

<u>Originating Biopsy Diagnosis</u>: DLBCL, GCB

<u>Prior Indolent Lymphoma</u>: None

<u>Sex</u>: Male

<u>Race</u>: Other

<u>Ethnicity</u>: Hispanic or Latino

<u>Age at diagnosis</u>: 58

<u>Age at PDX biopsy</u>: 60

<u># LoTs Pre Biopsy</u> (best response): 3 (CR)

<u># LoTs Post Biopsy</u> (best response): 6 (PD)

<u>Therapies</u>: Chemotherapy; Chemotherapy; CART; Immunotherapy; Chemotherapy; Targeted; Targeted; Targeted; Targeted

**Immunohistochemical Staining:**

Hans COO call: GCB

BCL6 (+)

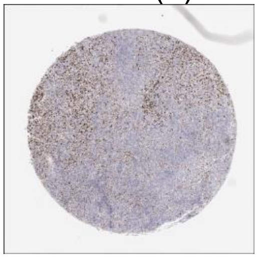

CD10 (+)

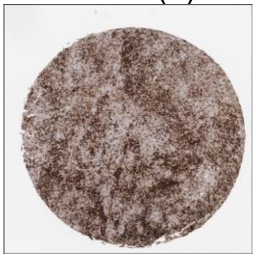

MUM1 (+)

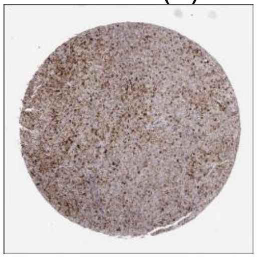

MYC (+)

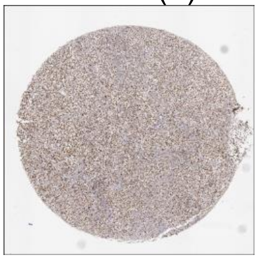

CD20 (+)

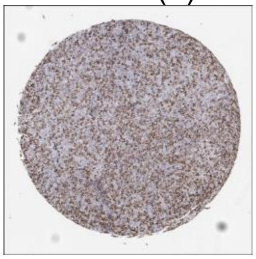

CD19 (+)

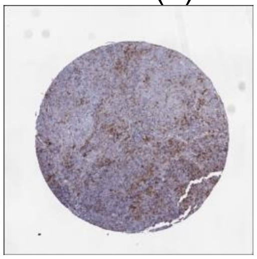

**DLBCL90 call:** GCB, DZsig+

**Genetic Characteristics:**

BCL2 FISH: Negative

MYC FISH: Negative

BCL6 FISH: Negative

Copy Number Alterations:

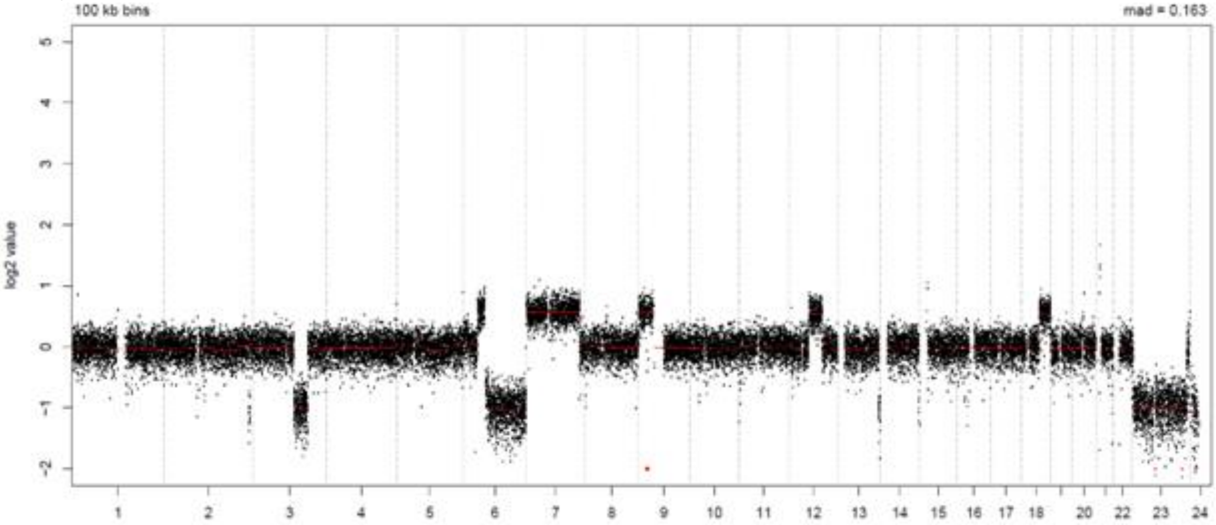


LymphGen Class (Confidence): BN2

BN2: 0.53

EZB: 0.34

N1: 0.07

MCD: 0.00

ST2: 0.24

A53: 0.06

### LPMI020

Source: MDACC

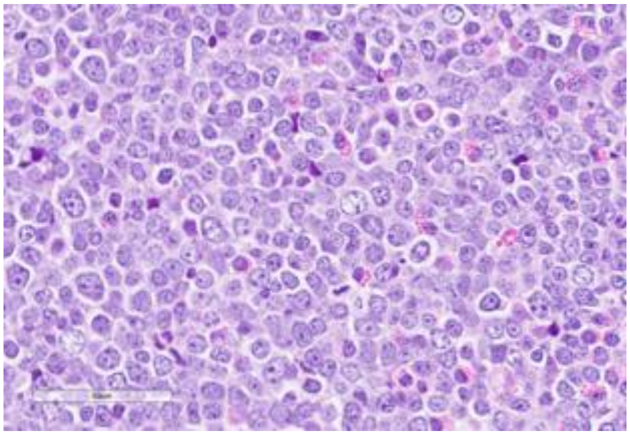

**Clinical Information**

Originating Biopsy Diagnosis: DLBCL, NOS

Prior Indolent Lymphoma: None

Sex: Female

Race: White or Caucasian

Ethnicity: Not Hispanic or Latino

Age at diagnosis: 74

Age at PDX biopsy: 80

### LoTs Pre Biopsy (best response): 1 (CR)

### LoTs Post Biopsy (best response): 1 (CR)

Therapies: Chemotherapy; Chemotherapy

**Immunohistochemical Staining:**

Hans COO call: Non-GCB

BCL6 (+)

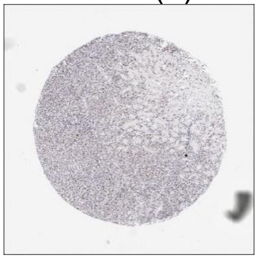

CD10 (-)

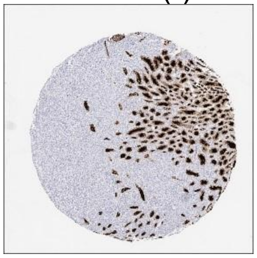

MUM1 (+)

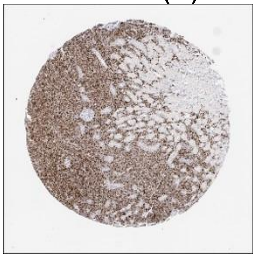

MYC (-)

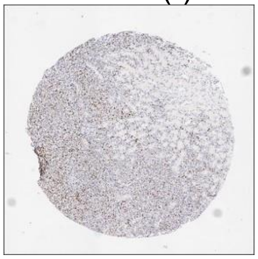

CD20 (+)

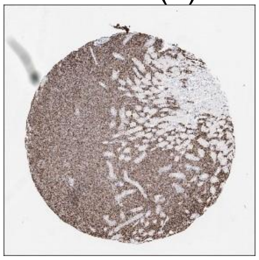

CD19 (+)

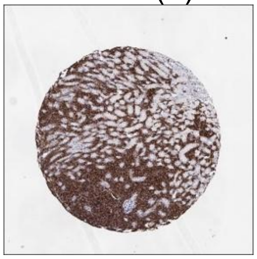

**DLBCL90 call:** ABC

**Genetic Characteristics:**

BCL2 FISH: Negative MYC FISH: Negative BCL6 FISH: Positive

Copy Number Alterations:

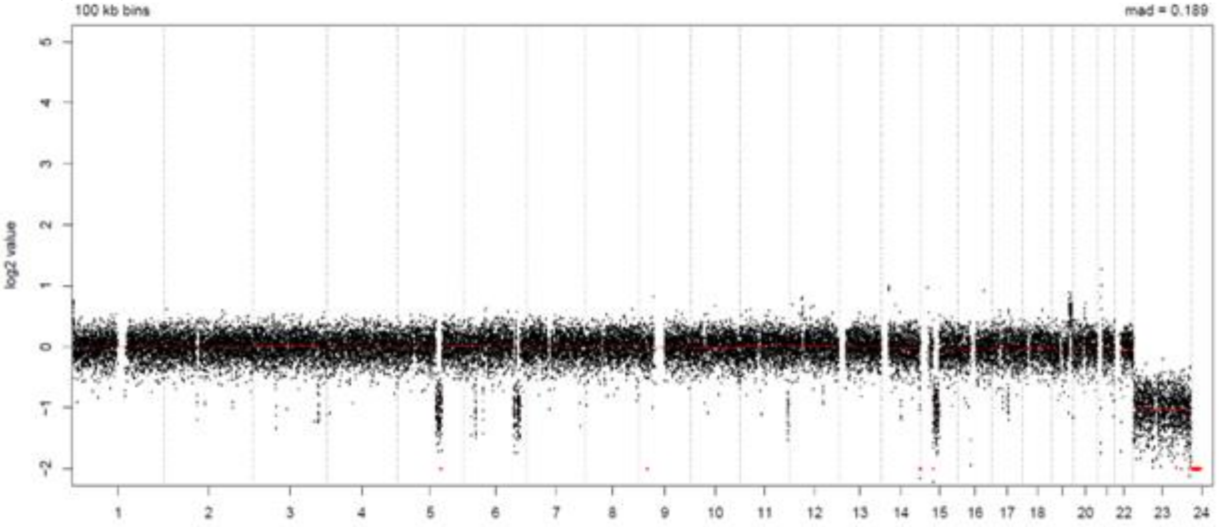


LymphGen Class (Confidence): BN2

BN2: 0.99 EZB: 0.00 N1: 0.07
MCD: 0.00 ST2: 0.01 A53: 0.00

### LPMI037

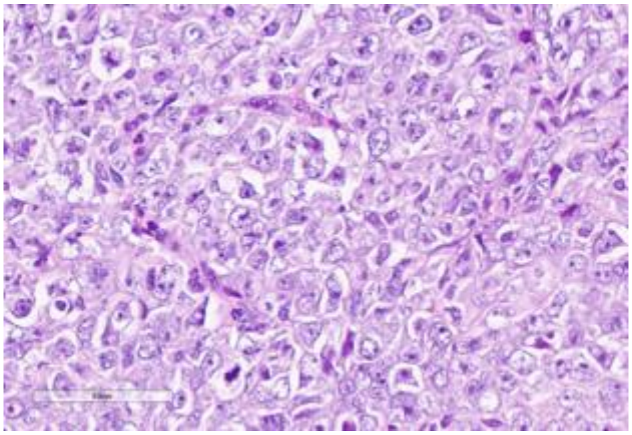

Source: MDACC

**Clinical Information**

Originating Biopsy Diagnosis: PMBCL

Prior Indolent Lymphoma: None

Sex: Male

Race: Black or African American

Ethnicity: Hispanic or Latino

Age at diagnosis: 37

Age at PDX biopsy: 39

### LoTs Pre Biopsy (best response): 5 (PD)

### LoTs Post Biopsy (best response): 1 (PD)

Therapies: Chemotherapy; Chemotherapy; Chemotherapy; CART; CART; Immunotherapy

**Immunohistochemical Staining:**

Hans COO call: GCB

BCL6 (+)

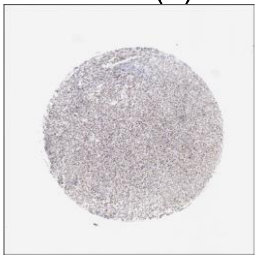

CD10 (+)

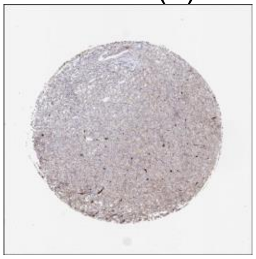

MUM1 (+)

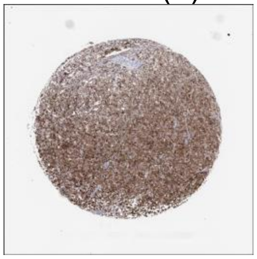

MYC (+)

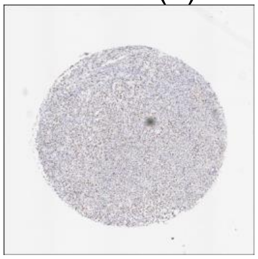

CD20 (+)

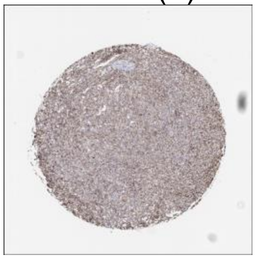

CD19 (+)

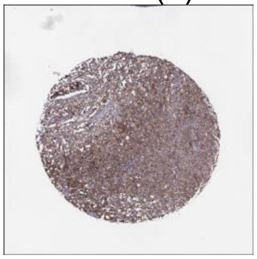

**DLBCL90 call:** PMBL

**Genetic Characteristics:**

BCL2 FISH: Negative

MYC FISH: Negative

BCL6 FISH: Positive

Copy Number Alterations:

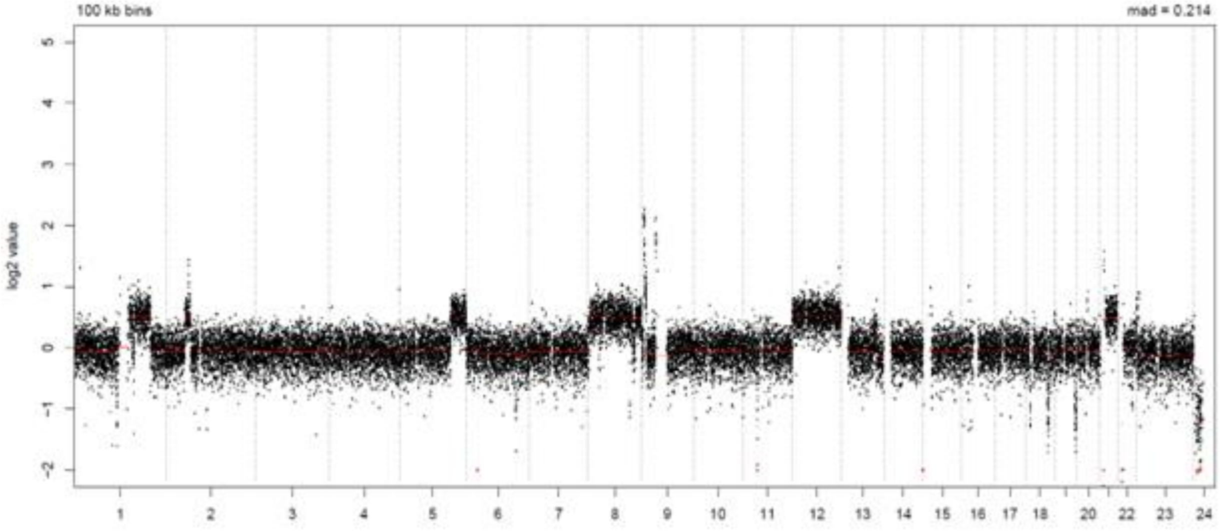


LymphGen Class (Confidence): BN2/ST2

BN2: 0.91

EZB: 0.48

N1: 0.07

MCD: 0.03

ST2: 0.99

A53: 0.00

### LPMI065

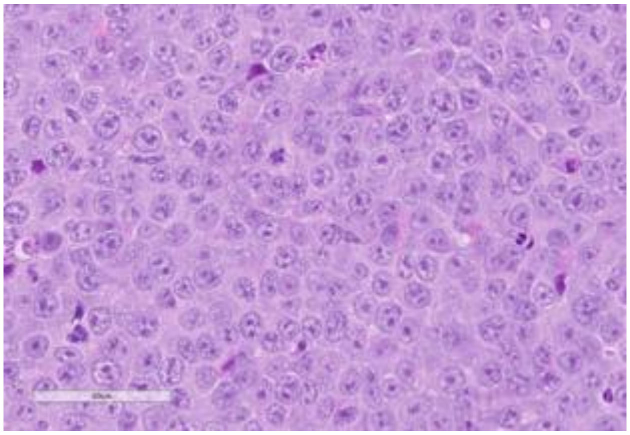

Source: MDACC

**Clinical Information**

Originating Biopsy Diagnosis: Grey Zone Lymphoma

Prior Indolent Lymphoma: None

Sex: Male

Race: White or Caucasian

Ethnicity: Not Hispanic or Latino

Age at diagnosis: 27

Age at PDX biopsy: 29

### LoTs Pre Biopsy (best response): 7 (PD)

### LoTs Post Biopsy (best response): 1 (PD)

Therapies: Chemotherapy; Chemotherapy; Chemotherapy; Immunotherapy; Chemotherapy; CART; Immunotherapy; ADC

**Immunohistochemical Staining:**

Hans COO call: Non-GCB

BCL6 (-)

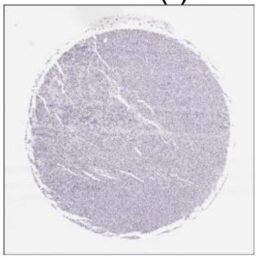

CD10 (-)

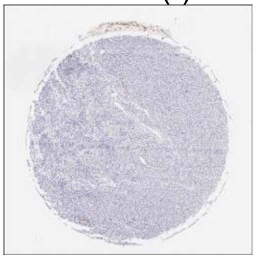

MUM1 (+)

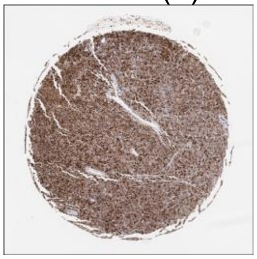

MYC (+)

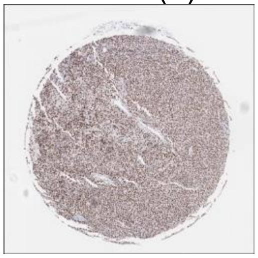

CD20 (+)

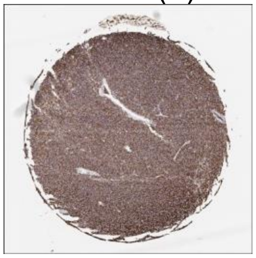

CD19 (+)

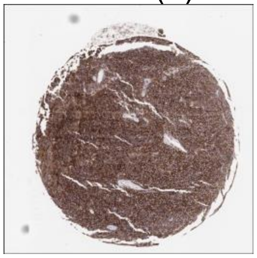

**DLBCL90 call:** PMBL

**Genetic Characteristics:**

| | | |
|---|---|---|
| BCL2 FISH: Negative | MYC FISH: Positive | BCL6 FISH: Negative |

Copy Number Alterations:

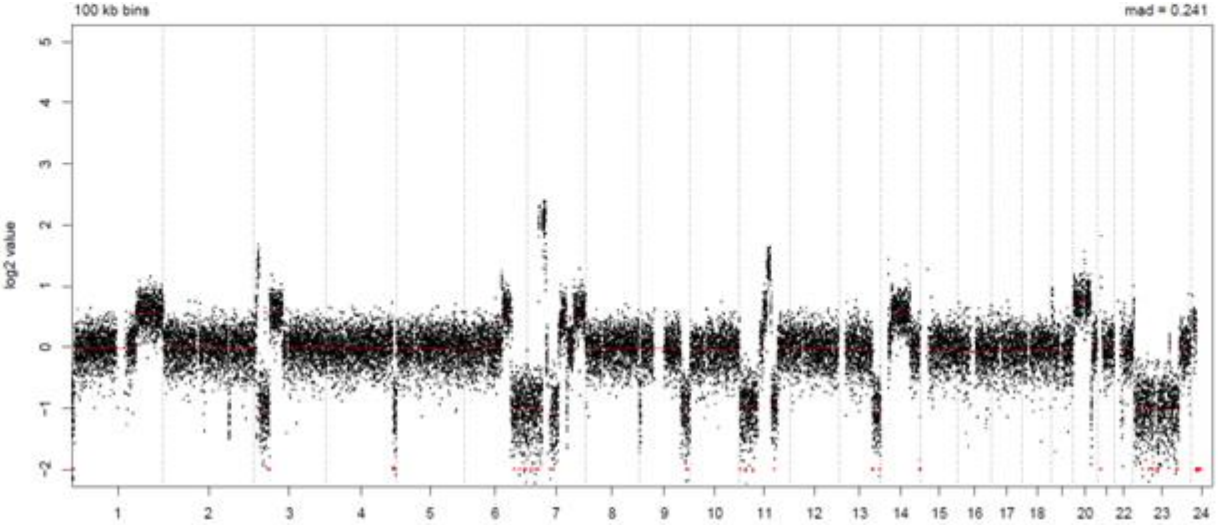


LymphGen Class (Confidence): ST2

| | | |
|---|---|---|
| BN2: 0.17 | EZB: 0.01 | N1: 0.07 |
| MCD: 0.00 | ST2: 0.98 | A53: 0.02 |

### LPMI147

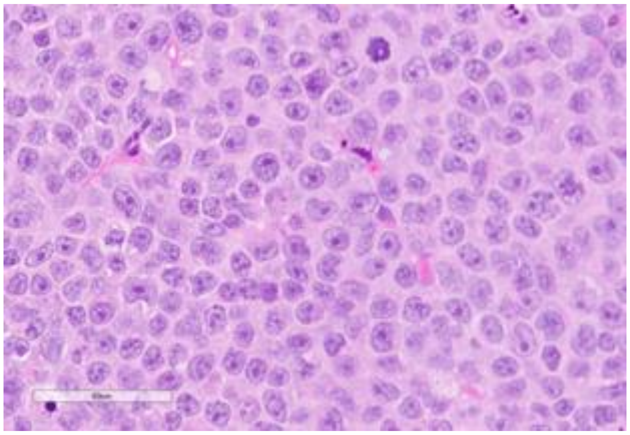

Source: MDACC

**Clinical Information**

Originating Biopsy Diagnosis: DLBCL, NOS

Prior Indolent Lymphoma: None

Sex: Female

Race: White or Caucasian

Ethnicity: Not Hispanic or Latino

Age at diagnosis: 68

Age at PDX biopsy: 78

### LoTs Pre Biopsy (best response): 5 (PD)

### LoTs Post Biopsy (best response): 1 (CR)

Therapies: Chemotherapy; Targeted; ADC; Chemotherapy; Immunotherapy; CART

**Immunohistochemical Staining:**

Hans COO call: GCB

BCL6 (+)

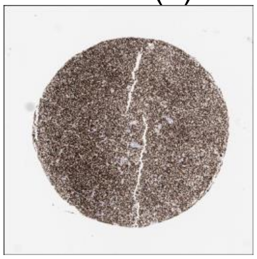

CD10 (+)

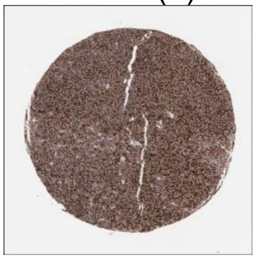

MUM1 (-)

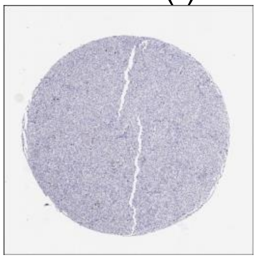

MYC (+)

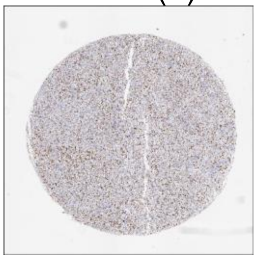

CD20 (+)

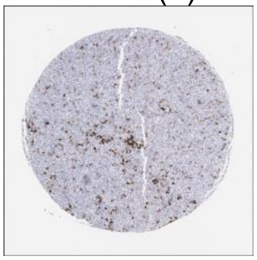

CD19 (+)

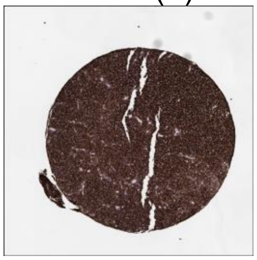

**DLBCL90 call:** GCB, DZsig+

**Genetic Characteristics:**

BCL2 FISH: Positive

MYC FISH: Negative

BCL6 FISH: Negative

Copy Number Alterations:

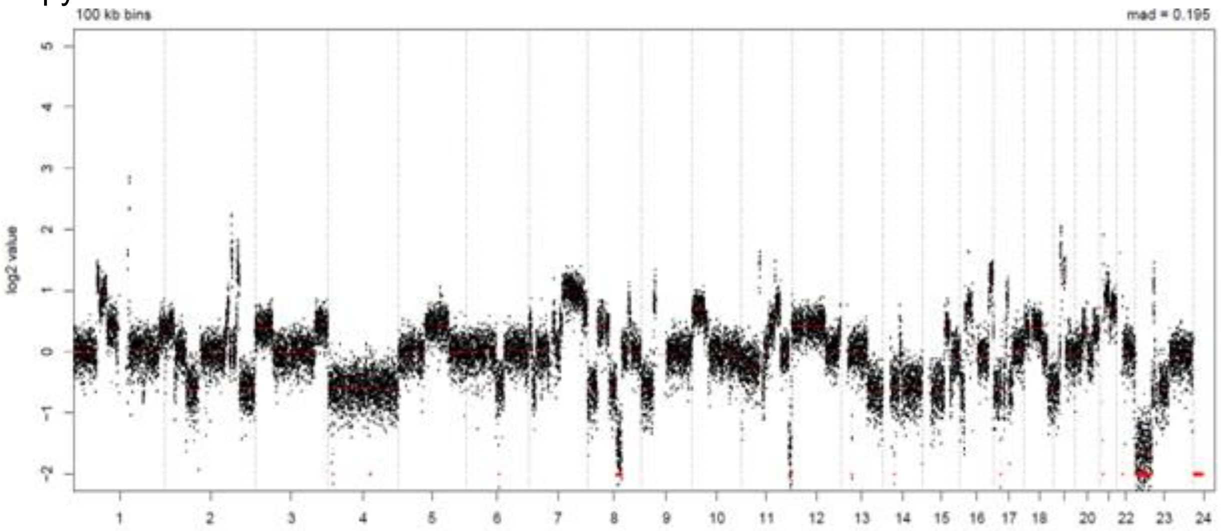


LymphGen Class (Confidence): EZB/ST2/A53

BN2: 0.12
MCD: 0.00

EZB: 0.99
ST2: 0.94

N1: 0.07
A53: 0.99

### LPMI195

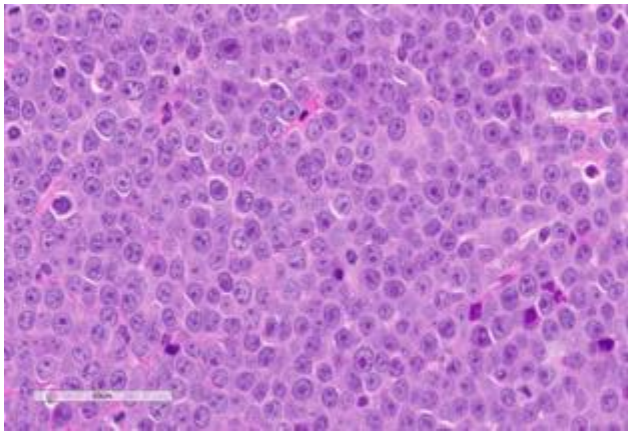

Source: MDACC

**Clinical Information**

Originating Biopsy Diagnosis: DLBCL, ABC

Prior Indolent Lymphoma: None

Sex: Female

Race: White or Caucasian

Ethnicity: Hispanic or Latino

Age at diagnosis: 59

Age at PDX biopsy: 59

### LoTs Pre Biopsy (best response): 0 (NA)

### LoTs Post Biopsy (best response): 1 (CR)

Therapies: Chemotherapy

**Immunohistochemical Staining:**

Hans COO call: Non-GCB

BCL6 (+)

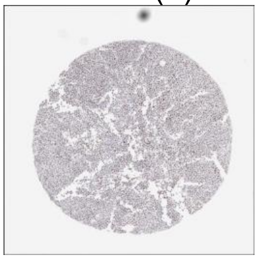

CD10 (-)

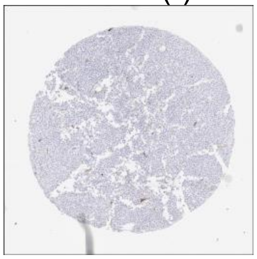

MUM1 (+)

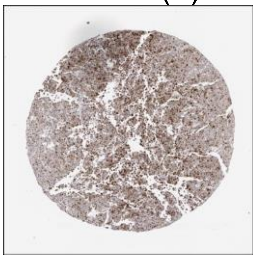

MYC (+)

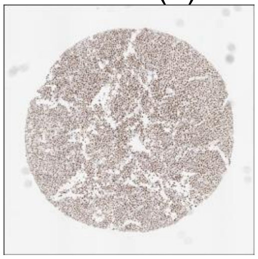

CD20 (+)

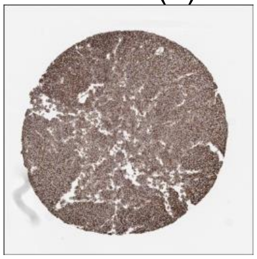

CD19 (+)

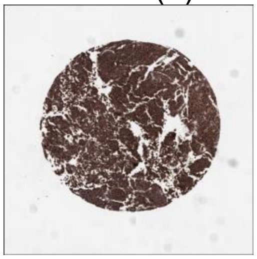

**DLBCL90 call:** ABC

**Genetic Characteristics:**

| | | |
|---|---|---|
| BCL2 FISH: Negative | MYC FISH: Positive | BCL6 FISH: Abnormal |

Copy Number Alterations:

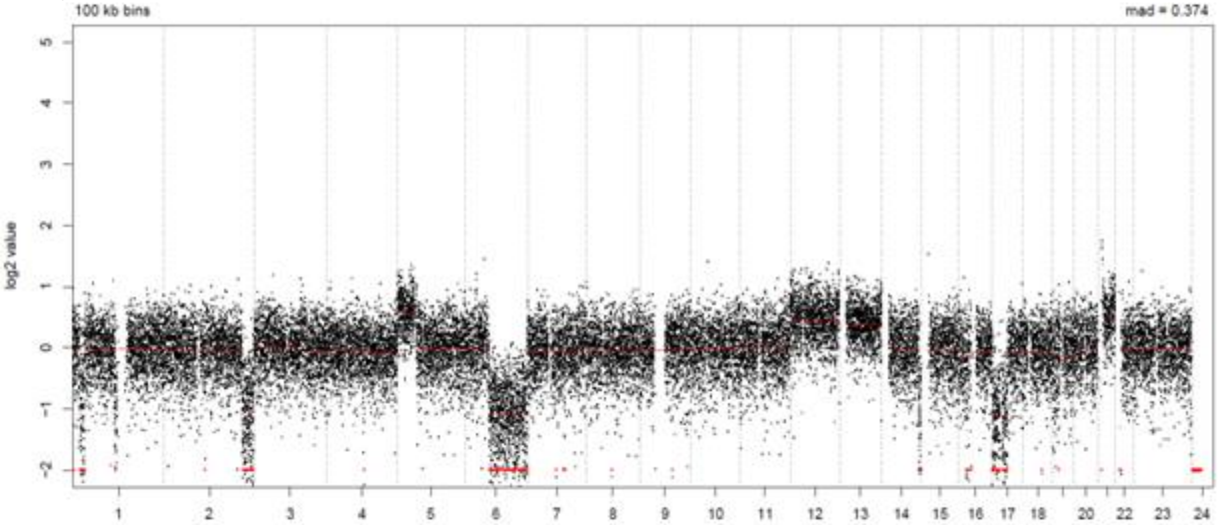


LymphGen Class (Confidence): A53

| | | |
|---|---|---|
| BN2: 0.34 | EZB: 0.72 | N1: 0.07 |
| MCD: 0.00 | ST2: 0.00 | A53: 0.98 |

### LPMI203

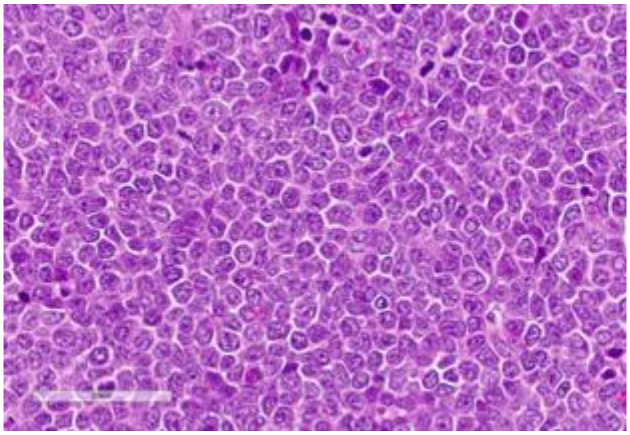

Source: MDACC

**Clinical Information**

Originating Biopsy Diagnosis: HGBCL

Prior Indolent Lymphoma: None

Sex: Female

Race: White or Caucasian

Ethnicity: Not Hispanic or Latino

Age at diagnosis: 67

Age at PDX biopsy: 67

### LoTs Pre Biopsy (best response): 2 (CR)

### LoTs Post Biopsy (best response): 2 (PD)

Therapies: Chemotherapy; CART; Immunotherapy; Chemotherapy

**Immunohistochemical Staining:**

Hans COO call: GCB

BCL6 (-)

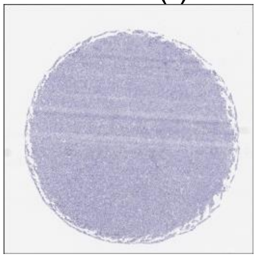

CD10 (+)

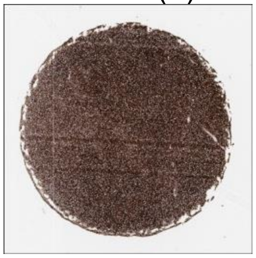

MUM1 (+)

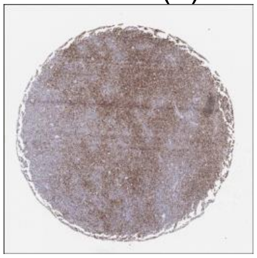

MYC (+)

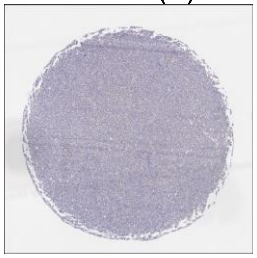

CD20 (+)

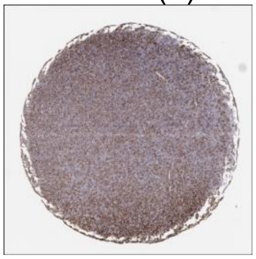

CD19 (+)

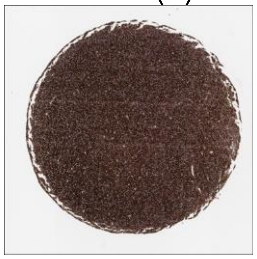

**DLBCL90 call:** UNCLASS, DZsig+

**Genetic Characteristics:**

BCL2 FISH: Positive

MYC FISH: Positive

BCL6 FISH: Negative

Copy Number Alterations:

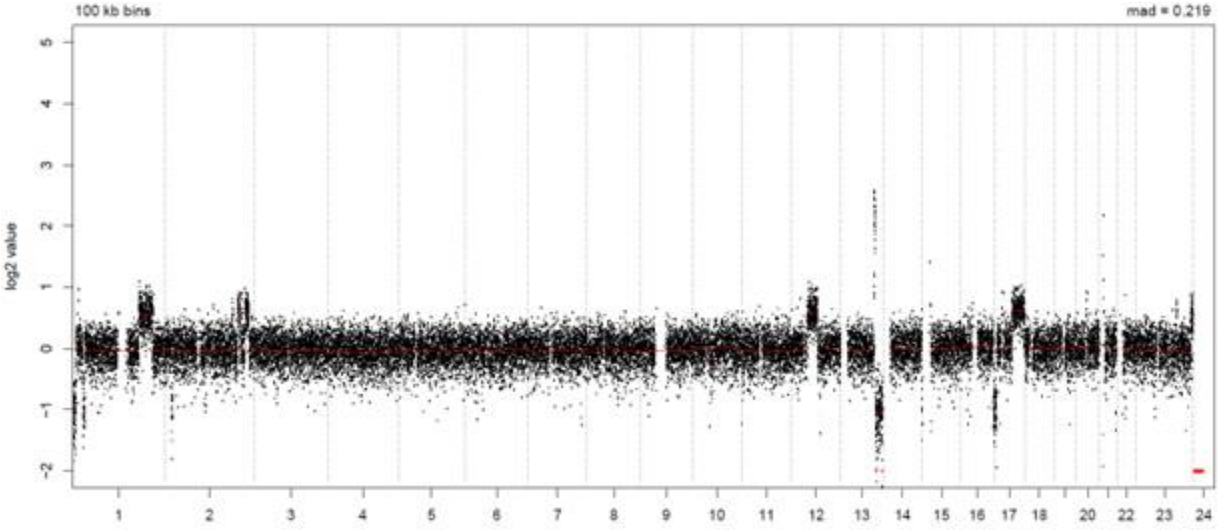


LymphGen Class (Confidence): EZB

BN2: 0.01

EZB: 0.83

N1: 0.07

MCD: 0.00

ST2: 0.15

A53: 0.02

### LPMI204

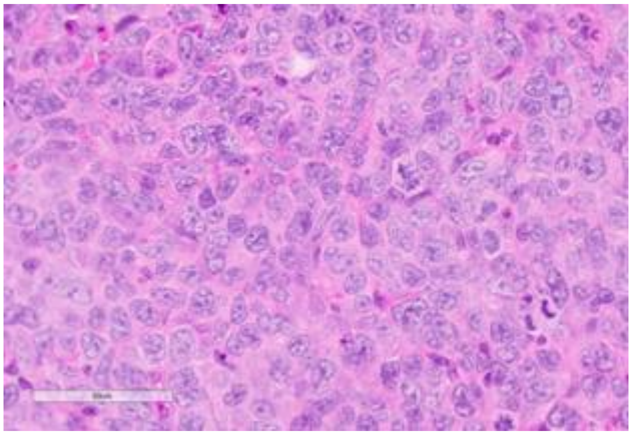

Source: MDACC

**Clinical Information**

<u>Originating Biopsy Diagnosis</u>: DLBCL, NOS

<u>Prior Indolent Lymphoma</u>: Follicular Lymphoma

<u>Sex</u>: Male

<u>Race</u>: White or Caucasian

<u>Ethnicity</u>: Not Hispanic or Latino

<u>Age at diagnosis</u>: 47

<u>Age at PDX biopsy</u>: 59

<u># LoTs Pre Biopsy</u> (best response): 9 (PD)

<u># LoTs Post Biopsy</u> (best response): 1 (PD)

<u>Therapies</u>: Chemotherapy; Chemotherapy; Chemotherapy; Chemotherapy; Targeted; Chemotherapy; RT; Chemotherapy; CART; Targeted

**Immunohistochemical Staining:**

Hans COO call: GCB

BCL6 (+)

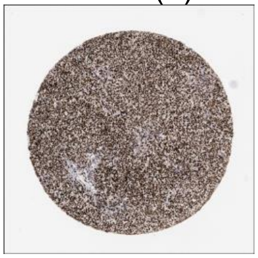

CD10 (+)

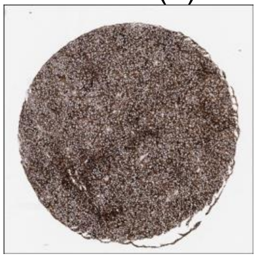

MUM1 (+)

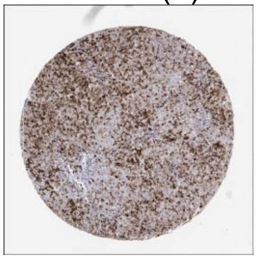

MYC (-)

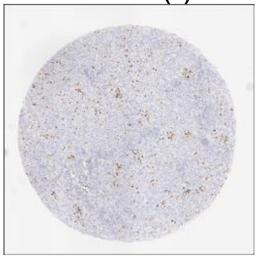

CD20 (+)

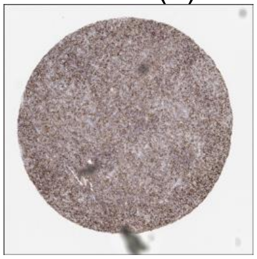

CD19 (+)

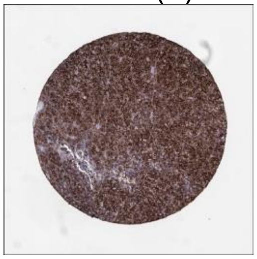

**DLBCL90 call:** GCB, DZsig+

**Genetic Characteristics:**

| | | |
|---|---|---|
| BCL2 FISH: Positive | MYC FISH: Negative | BCL6 FISH: Negative |

Copy Number Alterations:

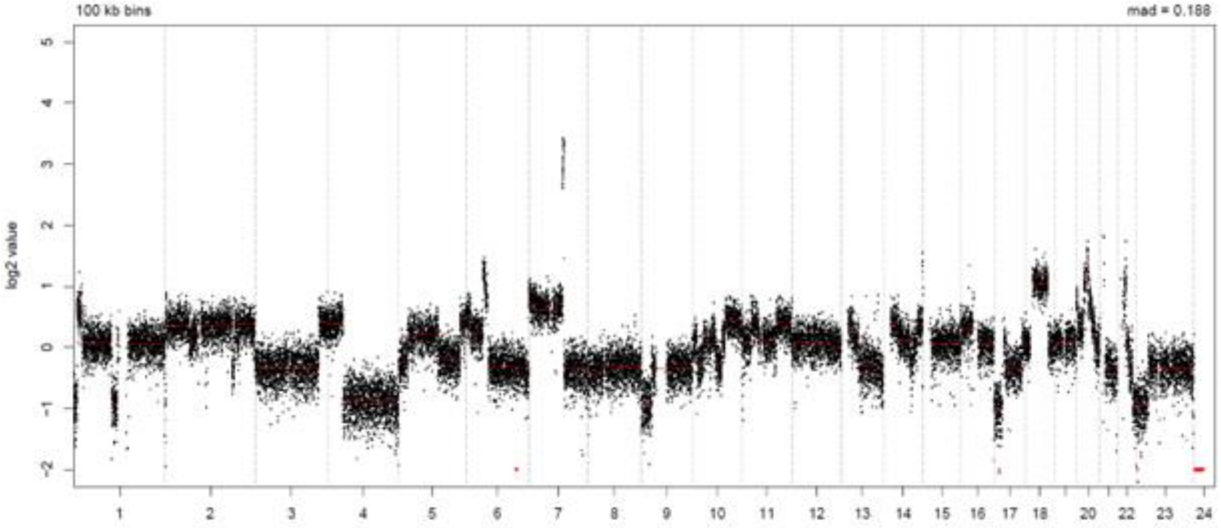


LymphGen Class (Confidence): EZB/A53

| | | |
|---|---|---|
| BN2: 0.07 | EZB: 0.99 | N1: 0.07 |
| MCD: 0.23 | ST2: 0.00 | A53: 0.99 |

### LPMI242

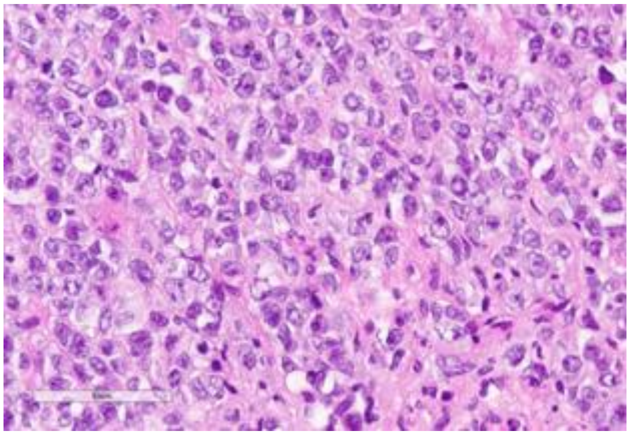

Source: MDACC

**Clinical Information**

Originating Biopsy Diagnosis: DLBCL, NOS

Prior Indolent Lymphoma: None

Sex: Male

Race: White or Caucasian

Ethnicity: Not Hispanic or Latino

Age at diagnosis: 51

Age at PDX biopsy: 52

### LoTs Pre Biopsy (best response): 4 (PD)

### LoTs Post Biopsy (best response): 5 (PR)

Therapies: Chemotherapy; Chemotherapy; Chemotherapy; ADC; CART; Chemotherapy; Chemotherapy; Targeted; Transplant

**Immunohistochemical Staining:**

Hans COO call: Non-GCB

BCL6 (+)

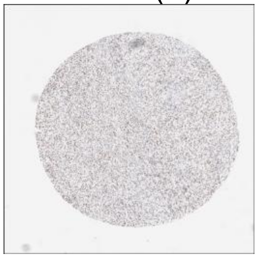

CD10 (+)

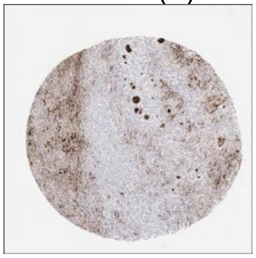

MUM1 (+)

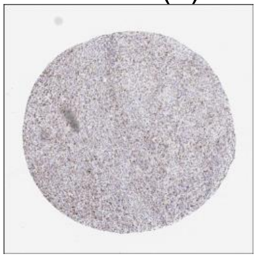

MYC (-)

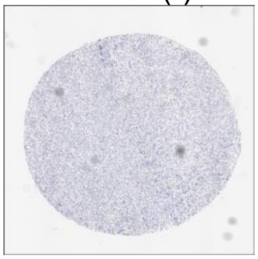

CD20 (+)

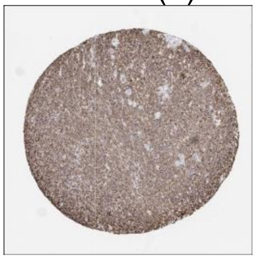

CD19 (+)

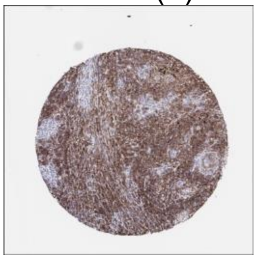

**DLBCL90 call:** GCB

**Genetic Characteristics:**

BCL2 FISH: Negative

MYC FISH: Positive

BCL6 FISH: Positive

Copy Number Alterations:

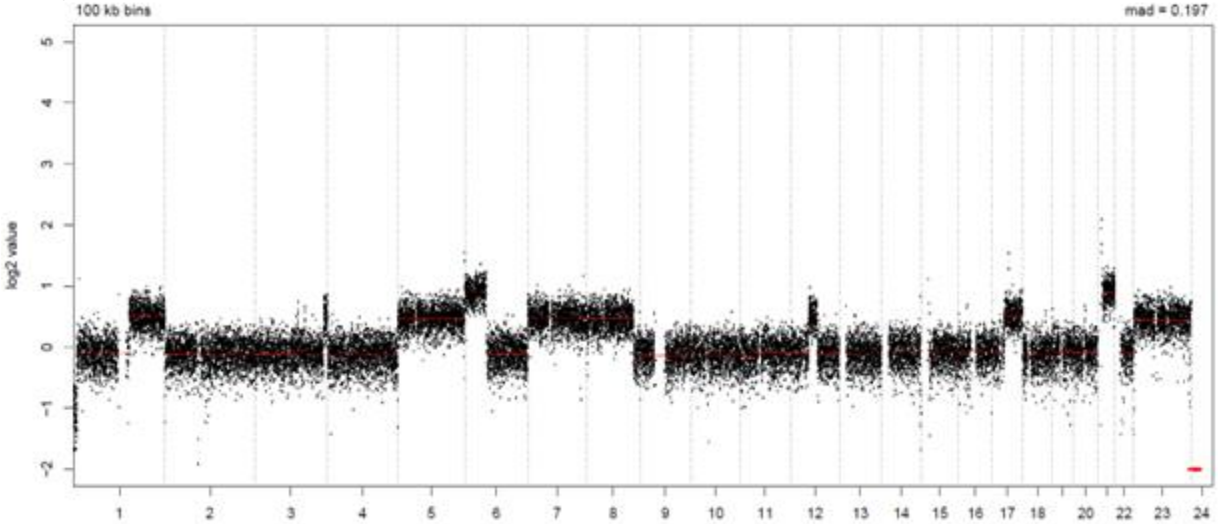


LymphGen Class (Confidence): EZB

| | | |
|---|---|---|
| BN2: 0.18 | EZB: 0.95 | N1: 0.07 |
| MCD: 0.00 | ST2: 0.00 | A53: 0.00 |

### LPMI251

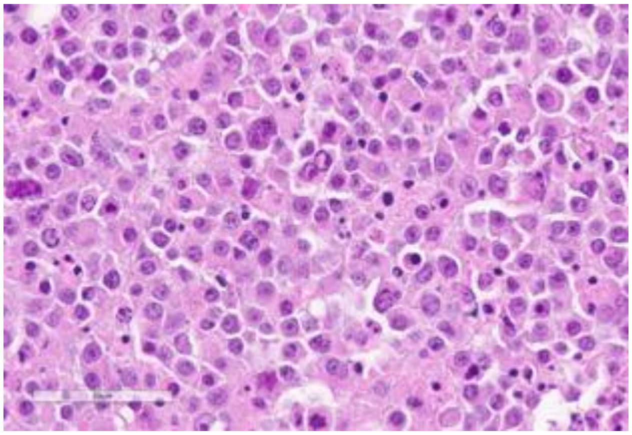

Source: MDACC

**Clinical Information**

Originating Biopsy Diagnosis: HGBCL

Prior Indolent Lymphoma: None

Sex: Male

Race: White or Caucasian

Ethnicity: Not Hispanic or Latino

Age at diagnosis: 64

Age at PDX biopsy: 65

### LoTs Pre Biopsy (best response): 2 (PD)

### LoTs Post Biopsy (best response): 1 (PD)

Therapies: Chemotherapy; CART; RT

**Immunohistochemical Staining:**

Hans COO call: Non-GCB

BCL6 (-)

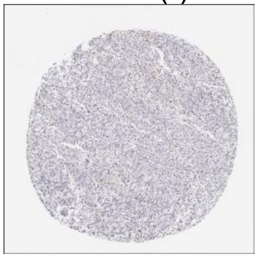

CD10 (-)

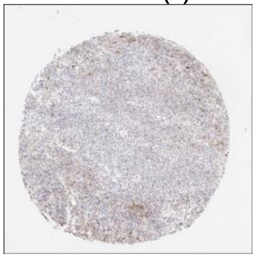

MUM1 (+)

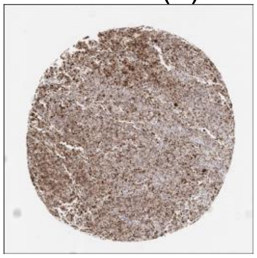

MYC (-)

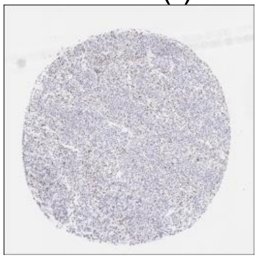

CD20 (+)

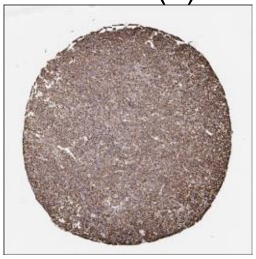

CD19 (+)

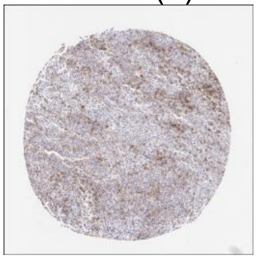

**DLBCL90 call:** GCB, DZsig+

**Genetic Characteristics:**

BCL2 FISH: Negative | MYC FISH: Negative | BCL6 FISH: Negative

Copy Number Alterations:

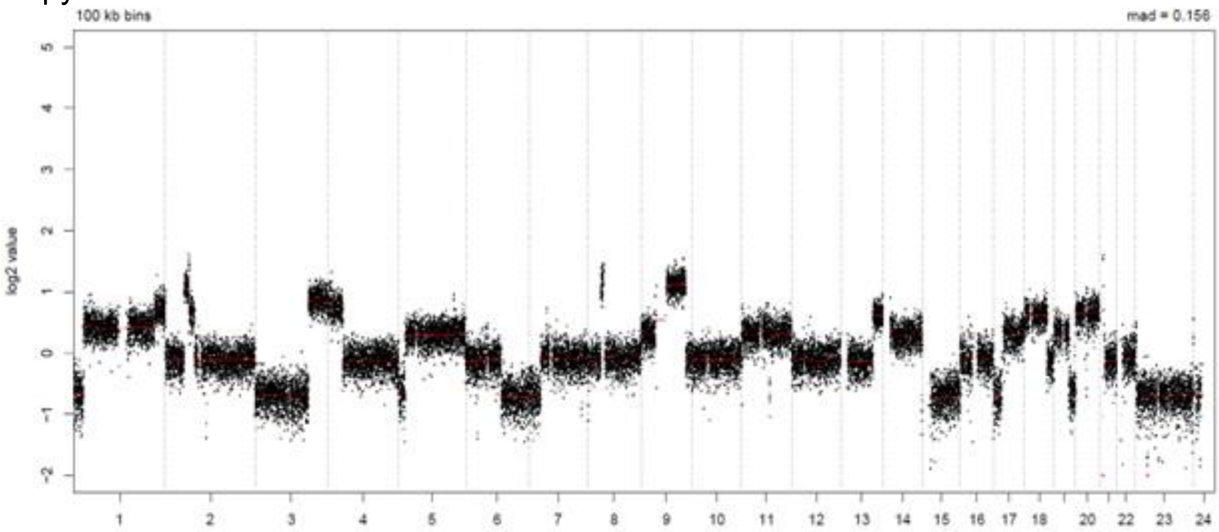


LymphGen Class (Confidence): Other

BN2: 0.22
EZB: 0.43
N1: 0.07
MCD: 0.00
ST2: 0.01
A53: 0.03

### LPMI300

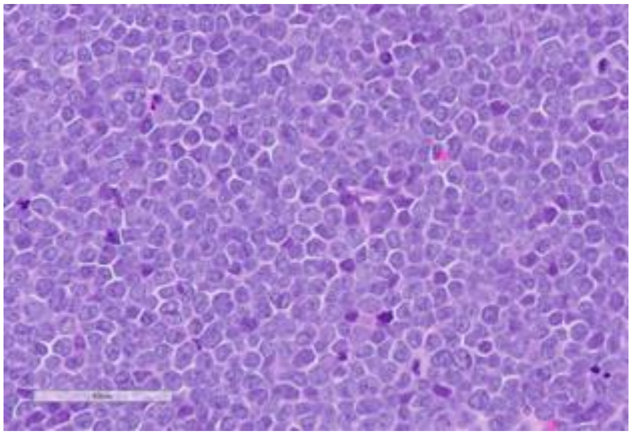

Source: MDACC

**Clinical Information**

Originating Biopsy Diagnosis: HGBCL

Prior Indolent Lymphoma: None

Sex: Female

Race: White or Caucasian

Ethnicity: Not Hispanic or Latino

Age at diagnosis: 30

Age at PDX biopsy: 31

### LoTs Pre Biopsy (best response): 4 (PD)

### LoTs Post Biopsy (best response): 2 (PD)

Therapies: Chemotherapy; Chemotherapy; Chemotherapy; CART; Targeted; Targeted

**Immunohistochemical Staining:**

Hans COO call: GCB

BCL6 (-)

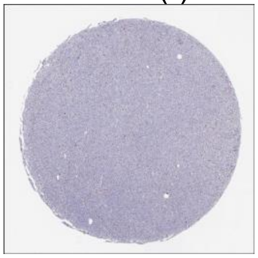

CD10 (+)

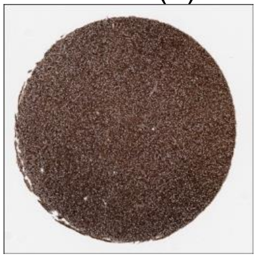

MUM1 (+)

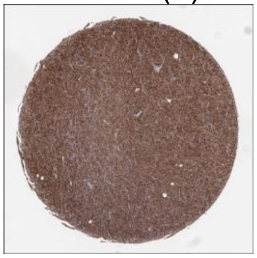

MYC (+)

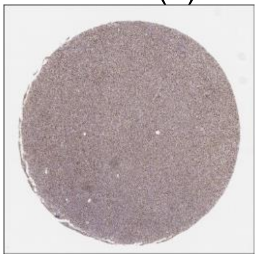

CD20 (+)

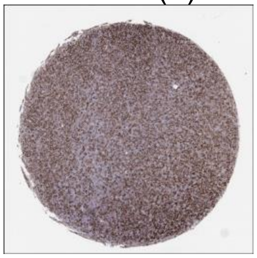

CD19 (+)

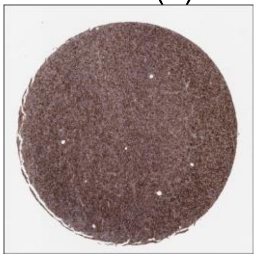

**DLBCL90 call:** ABC

**Genetic Characteristics:**

BCL2 FISH: Negative

MYC FISH: Positive

BCL6 FISH: Negative

Copy Number Alterations:

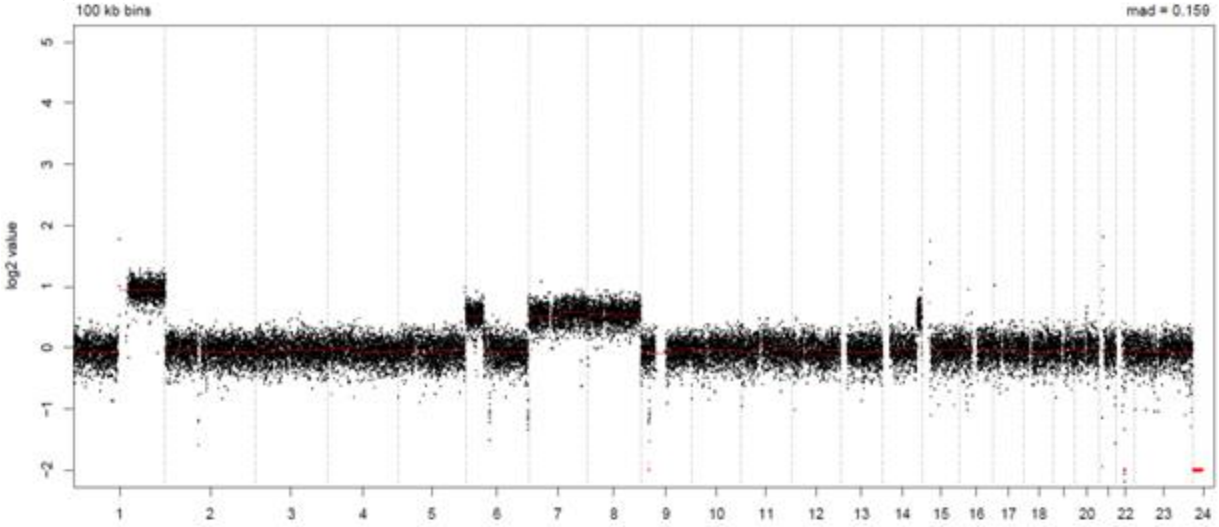


LymphGen Class (Confidence): Other

BN2: 0.01

EZB: 0.43

N1: 0.07

MCD: 0.00

ST2: 0.01

A53: 0.00

### LPMI322

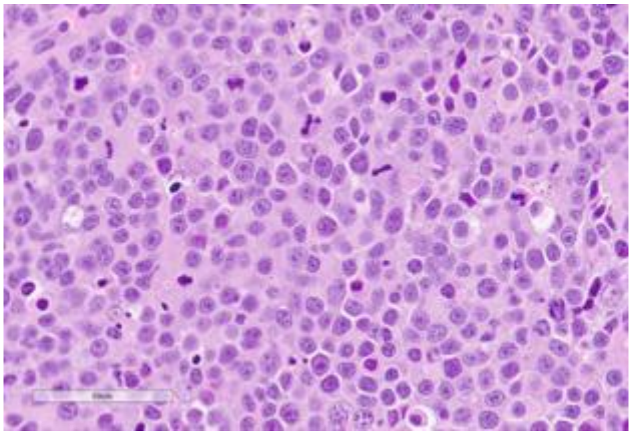

Source: MDACC

**Clinical Information**

Originating Biopsy Diagnosis: DLBCL, NOS

Prior Indolent Lymphoma: None

Sex: Male

Race: White or Caucasian

Ethnicity: Hispanic or Latino

Age at diagnosis: 80

Age at PDX biopsy: 81

### LoTs Pre Biopsy (best response): 3 (PD)

### LoTs Post Biopsy (best response): 0 (NA)

Therapies: Chemotherapy; Chemotherapy; CART

**Immunohistochemical Staining:**

Hans COO call: GCB

BCL6 (+)

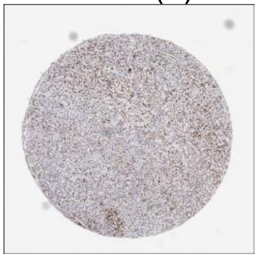

CD10 (+)

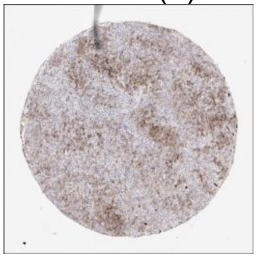

MUM1 (+)

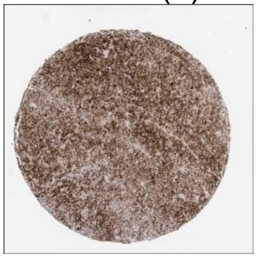

MYC (+)

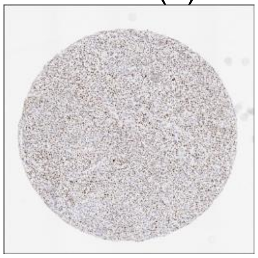

CD20 (+)

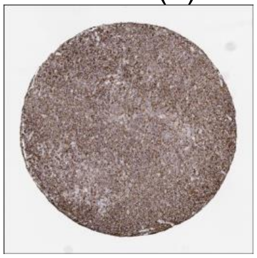

CD19 (+)

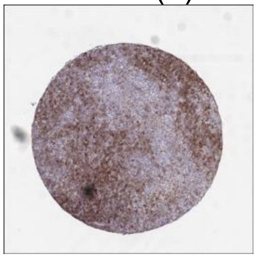

**DLBCL90 call:** ABC

**Genetic Characteristics:**

| | | |
|---|---|---|
| BCL2 FISH: Negative | MYC FISH: Negative | BCL6 FISH: Negative |

Copy Number Alterations:

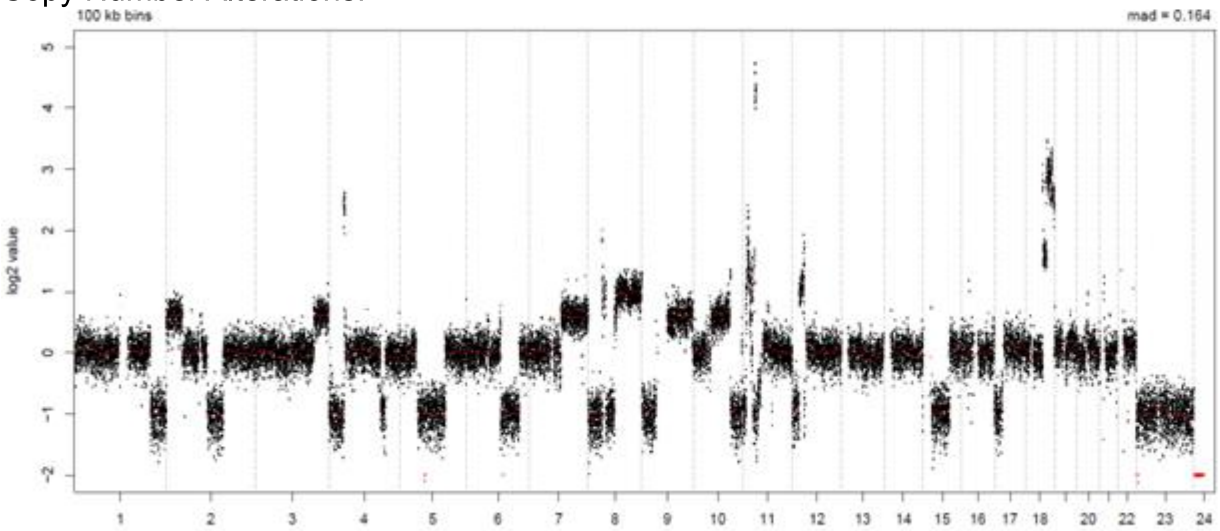


LymphGen Class (Confidence): ST2

| | | |
|---|---|---|
| BN2: 0.01 | EZB: 0.23 | N1: 0.07 |
| MCD: 0.73 | ST2: 0.95 | A53: 0.12 |

### LPMI329

Source: MDACC

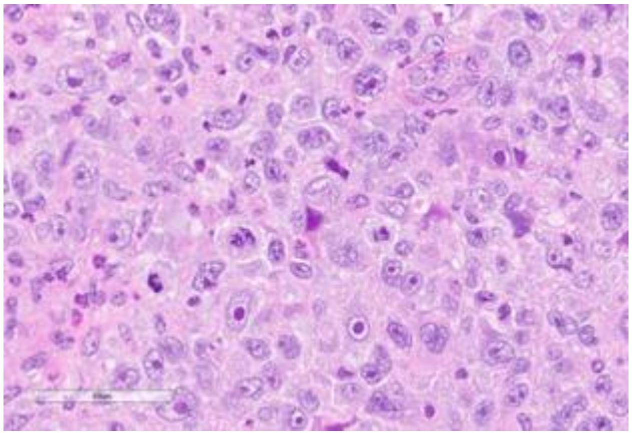

**Clinical Information**

Originating Biopsy Diagnosis: Grey Zone Lymphoma

Prior Indolent Lymphoma: None

Sex: Female

Race: White or Caucasian

Ethnicity: Not Hispanic or Latino

Age at diagnosis: 67

Age at PDX biopsy: 71

### LoTs Pre Biopsy (best response): 5 (CR)

### LoTs Post Biopsy (best response): 2 (CR)

Therapies: Chemotherapy; Chemotherapy; ADC; Chemotherapy; Immunotherapy; Chemotherapy; ADC

**Immunohistochemical Staining:**

Hans COO call: Non-GCB

BCL6 (+)

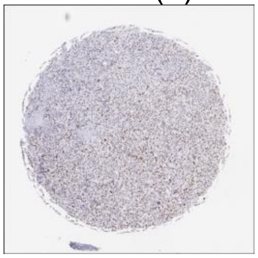

CD10 (-)

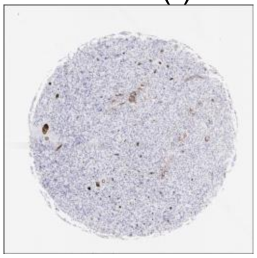

MUM1 (+)

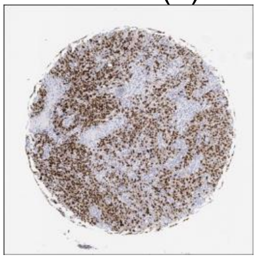

MYC (+)

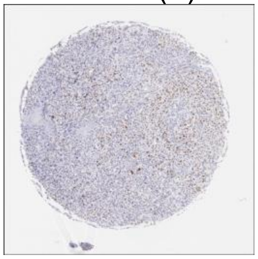

CD20 (-)

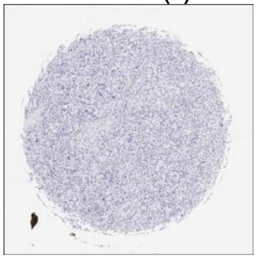

CD19 (-)

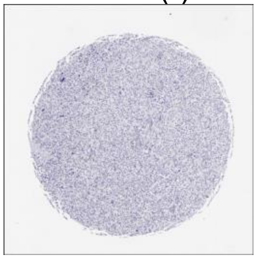

**DLBCL90 call:** PMBL

**Genetic Characteristics:**

BCL2 FISH: Positive

MYC FISH: Negative

BCL6 FISH: Negative

Copy Number Alterations:

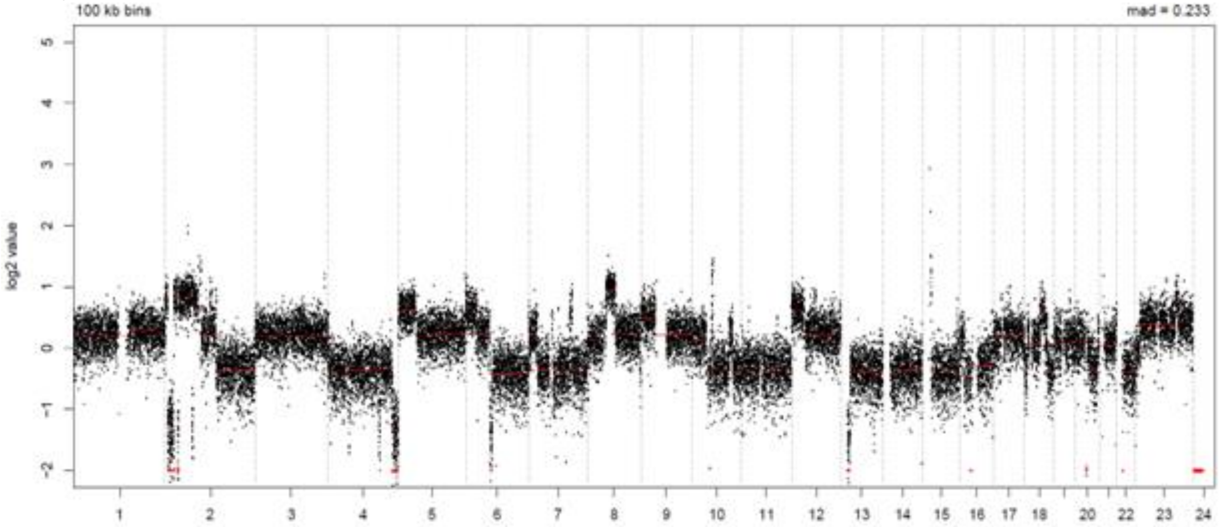


LymphGen Class (Confidence): EZB

BN2: 0.01

EZB: 0.63

N1: 0.07

MCD: 0.00

ST2: 0.00

A53: 0.18

### LPMI331

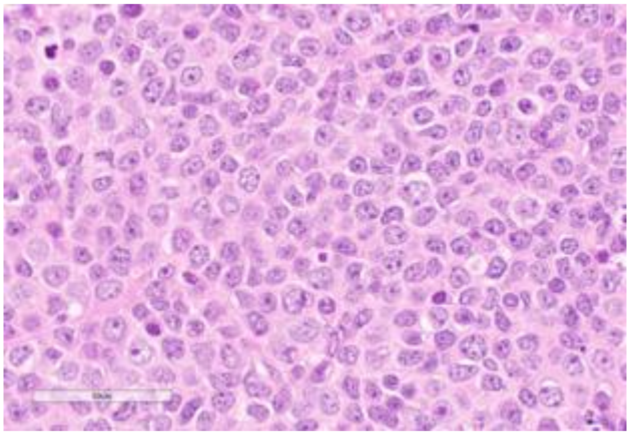

Source: MDACC

**Clinical Information**

<u>Originating Biopsy Diagnosis</u>: DLBCL, GCB

<u>Prior Indolent Lymphoma</u>: None

<u>Sex</u>: Male

<u>Race</u>: White or Caucasian

<u>Ethnicity</u>: Not Hispanic or Latino

<u>Age at diagnosis</u>: 57

<u>Age at PDX biopsy</u>: 57

<u># LoTs Pre Biopsy</u> (best response): 1 (PR)

<u># LoTs Post Biopsy</u> (best response): 5 (SD)

<u>Therapies</u>: CART; Chemotherapy; Chemotherapy; Transplant; Immunotherapy; Chemotherapy

**Immunohistochemical Staining:**

Hans COO call: GCB

BCL6 (+)

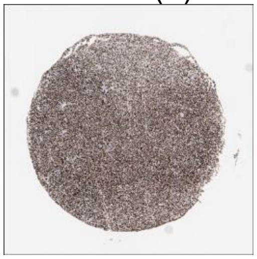

CD10 (+)

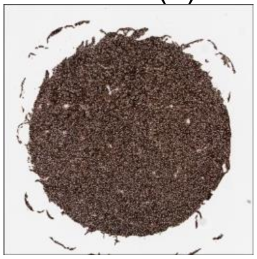

MUM1 (+)

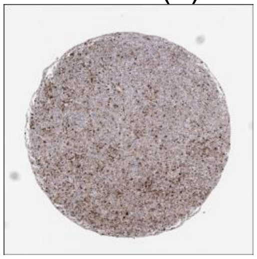

MYC (+)

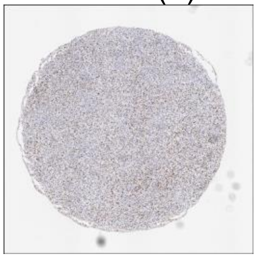

CD20 (+)

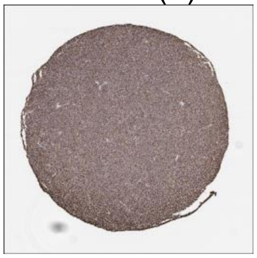

CD19 (+)

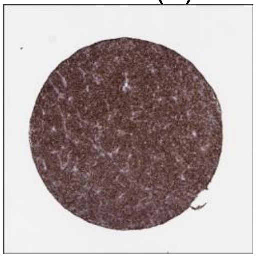

**DLBCL90 call:** GCB

**Genetic Characteristics:**

BCL2 FISH: Positive

MYC FISH: Abnormal

BCL6 FISH: Negative

Copy Number Alterations:

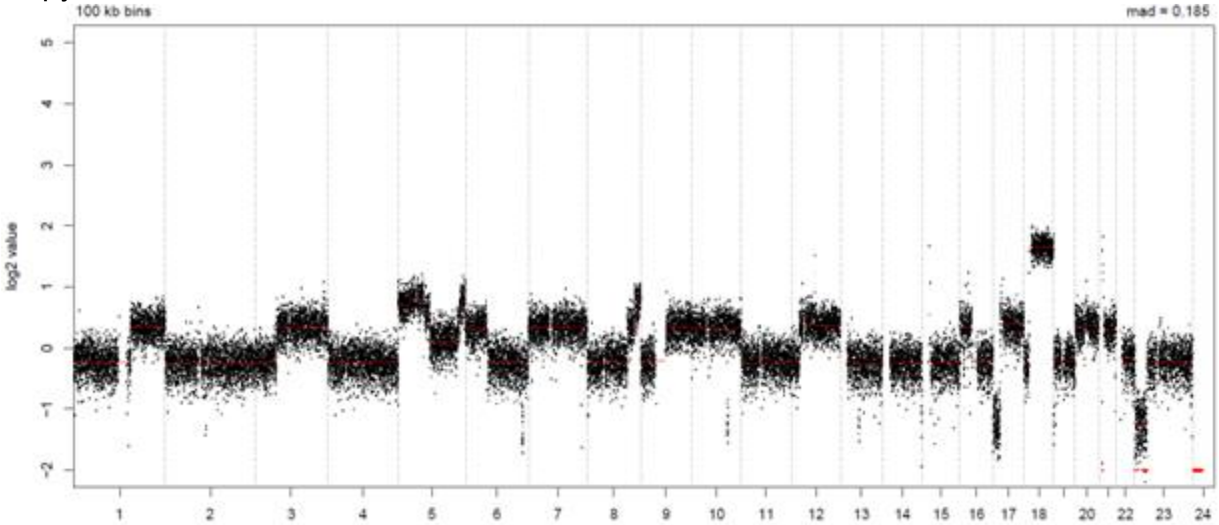


LymphGen Class (Confidence): EZB

BN2: 0.01

MCD: 0.00

EZB: 0.97

ST2: 0.00

N1: 0.07

A53: 0.00

### LPMI343

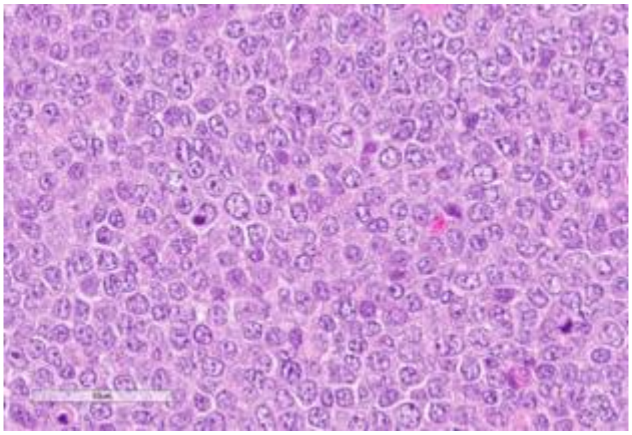

Source: MDACC

**Clinical Information**

Originating Biopsy Diagnosis: DLBCL, GCB

Prior Indolent Lymphoma: Marginal Zone Lymphoma

Sex: Male

Race: White or Caucasian

Ethnicity: Not Hispanic or Latino

Age at diagnosis: 69

Age at PDX biopsy: 71

### LoTs Pre Biopsy (best response): 8 (PR)

### LoTs Post Biopsy (best response): 0 (NA)

Therapies: Chemotherapy; Targeted; Targeted; Chemotherapy; Targeted; Other Cellular Therapy; Targeted; CART

**Immunohistochemical Staining:**

Hans COO call: GCB

BCL6 (+)

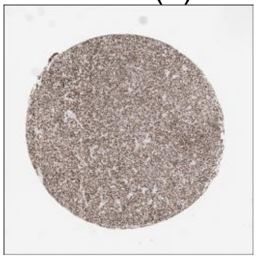

CD10 (+)

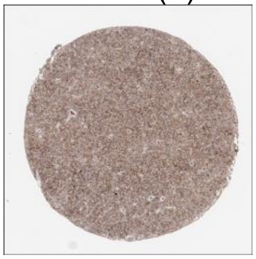

MUM1 (+)

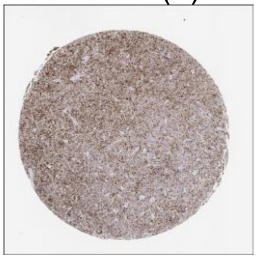

MYC (+)

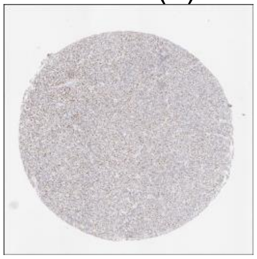

CD20 (+)

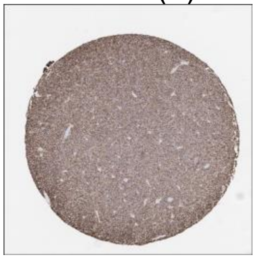

CD19 (+)

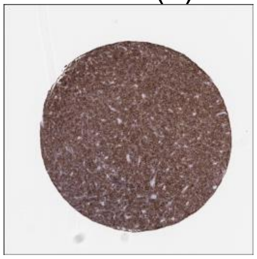

**DLBCL90 call:** ABC

**Genetic Characteristics:**

| | | |
|---|---|---|
| BCL2 FISH: Negative | MYC FISH: Negative | BCL6 FISH: Negative |

Copy Number Alterations:

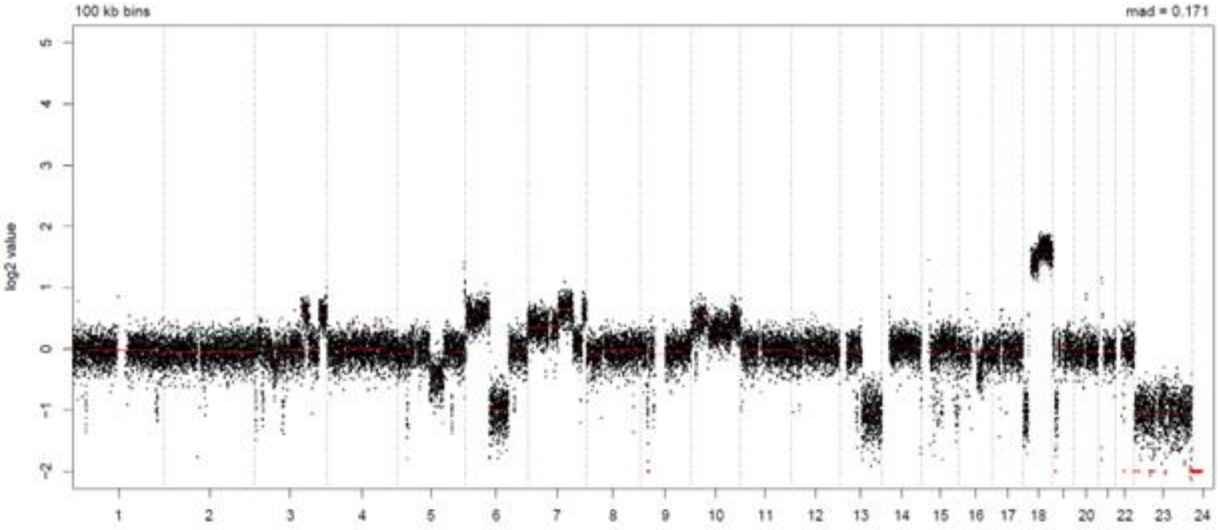


LymphGen Class (Confidence): ST2

| | | |
|---|---|---|
| BN2: 0.13 | EZB: 0.08 | N1: 0.07 |
| MCD: 0.12 | ST2: 0.86 | A53: 0.00 |

### LPMI351

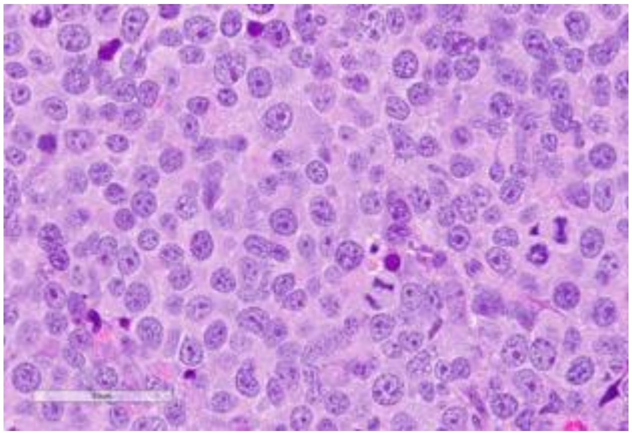

Source: MDACC

**Clinical Information**

Originating Biopsy Diagnosis: DLBCL, ABC

Prior Indolent Lymphoma: None

Sex: Male

Race: White or Caucasian

Ethnicity: Not Hispanic or Latino

Age at diagnosis: 91

Age at PDX biopsy: 91

### LoTs Pre Biopsy (best response): 1 (PD)

### LoTs Post Biopsy (best response): 1 (PD)

Therapies: Chemotherapy; Targeted

**Immunohistochemical Staining:**

Hans COO call: Non-GCB

BCL6 (+)

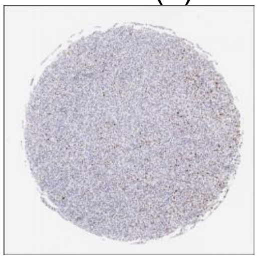

CD10 (-)

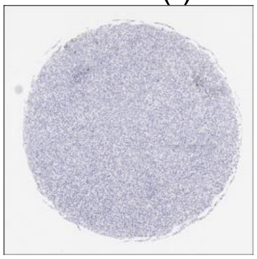

MUM1 (+)

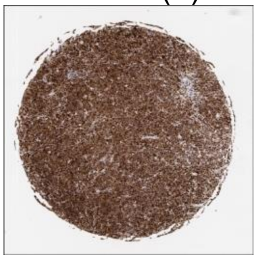

MYC (+)

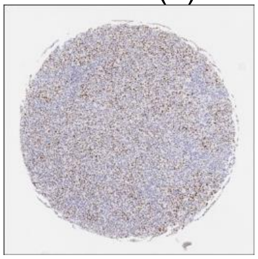

CD20 (+)

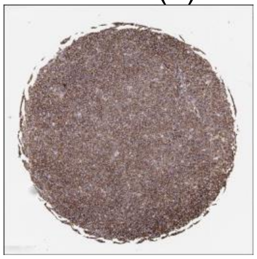

CD19 (+)

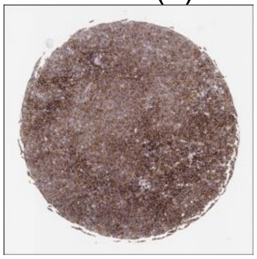

**DLBCL90 call:** ABC

**Genetic Characteristics:**

| | | |
|---|---|---|
| BCL2 FISH: Negative | MYC FISH: Negative | BCL6 FISH: Negative |

Copy Number Alterations:

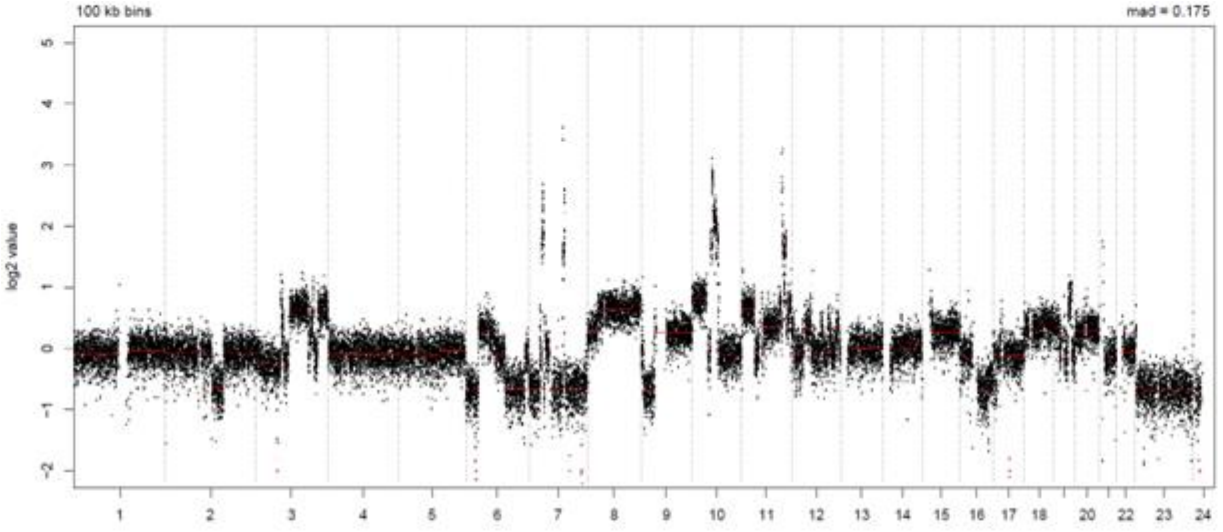


LymphGen Class (Confidence): Other

| | | |
|---|---|---|
| BN2: 0.25 | EZB: 0.12 | N1: 0.07 |
| MCD: 0.00 | ST2: 0.00 | A53: 0.00 |

### LPMI372

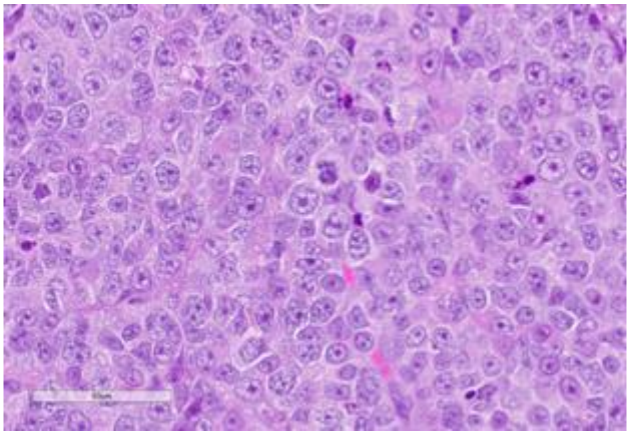

Source: MDACC

**Clinical Information**

<u>Originating Biopsy Diagnosis</u>: BCL

<u>Prior Indolent Lymphoma</u>: Follicular Lymphoma

<u>Sex</u>: Male

<u>Race</u>: Black or African American

<u>Ethnicity</u>: Not Hispanic or Latino

<u>Age at diagnosis</u>: 68

<u>Age at PDX biopsy</u>: 69

<u># LoTs Pre Biopsy</u> (best response): 5 (CR)

<u># LoTs Post Biopsy</u> (best response): 1 (PD)

<u>Therapies</u>: Chemotherapy; Chemotherapy; Chemotherapy; Targeted; CART; Immunotherapy

**Immunohistochemical Staining:**

Hans COO call: GCB

BCL6 (-)

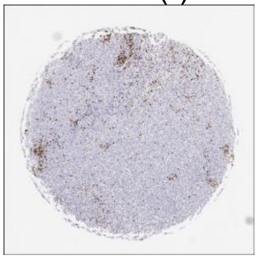

CD10 (+)

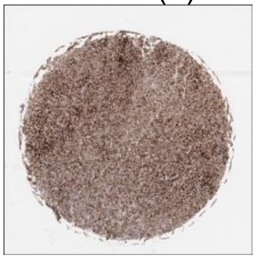

MUM1 (+)

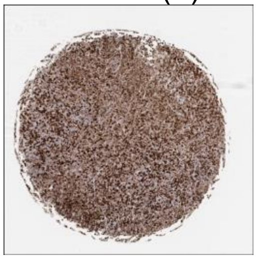

MYC (+)

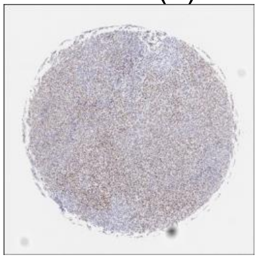

CD20 (+)

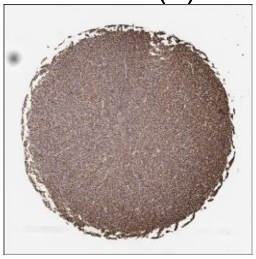

CD19 (+)

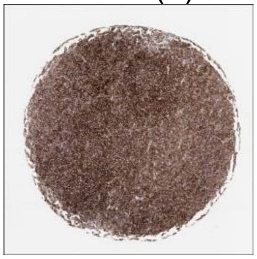

**DLBCL90 call:** GCB

**Genetic Characteristics:**

BCL2 FISH: Positive

MYC FISH: Positive

BCL6 FISH: Negative

Copy Number Alterations:

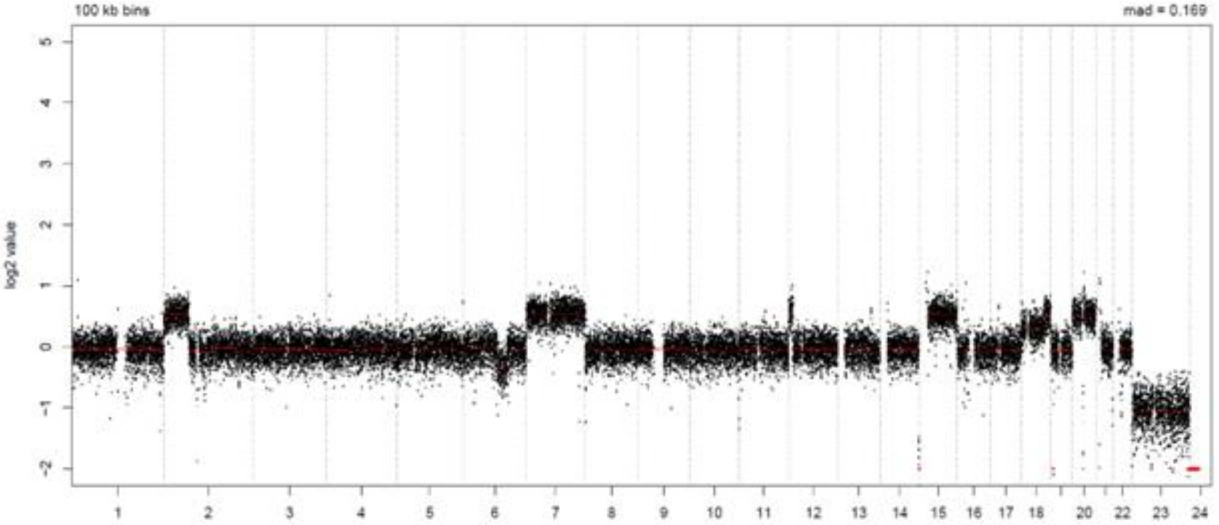


LymphGen Class (Confidence): EZB

BN2: 0.01

EZB: 0.94

N1: 0.07

MCD: 0.00

ST2: 0.36

A53: 0.00

### LPMI374

Source: MDACC

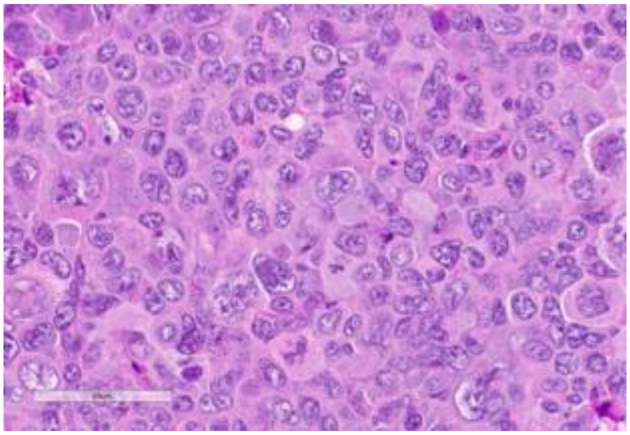

**Clinical Information**

Originating Biopsy Diagnosis: DLBCL, NOS

Prior Indolent Lymphoma: Follicular Lymphoma

Sex: Male

Race: White or Caucasian

Ethnicity: Not Hispanic or Latino

Age at diagnosis: 74

Age at PDX biopsy: 76

### LoTs Pre Biopsy (best response): 1 (PR)

### LoTs Post Biopsy (best response): 2 (PD)

Therapies: Chemotherapy; Chemotherapy; Targeted

**Immunohistochemical Staining:**

Hans COO call: GCB

BCL6 (+)

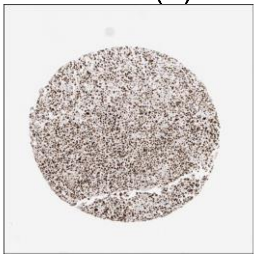

CD10 (+)

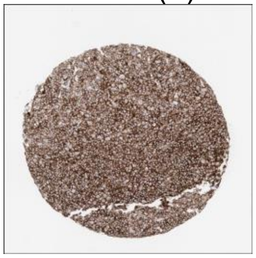

MUM1 (+)

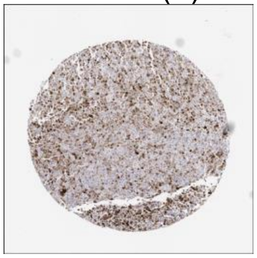

MYC (-)

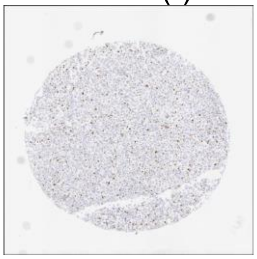

CD20 (+)

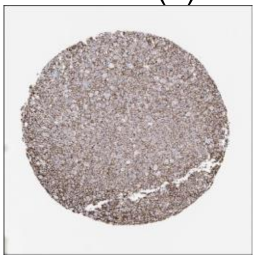

CD19 (+)

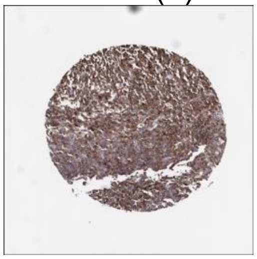

**DLBCL90 call:** GCB

**Genetic Characteristics:**

BCL2 FISH: Positive    MYC FISH: Negative    BCL6 FISH: Negative

Copy Number Alterations:

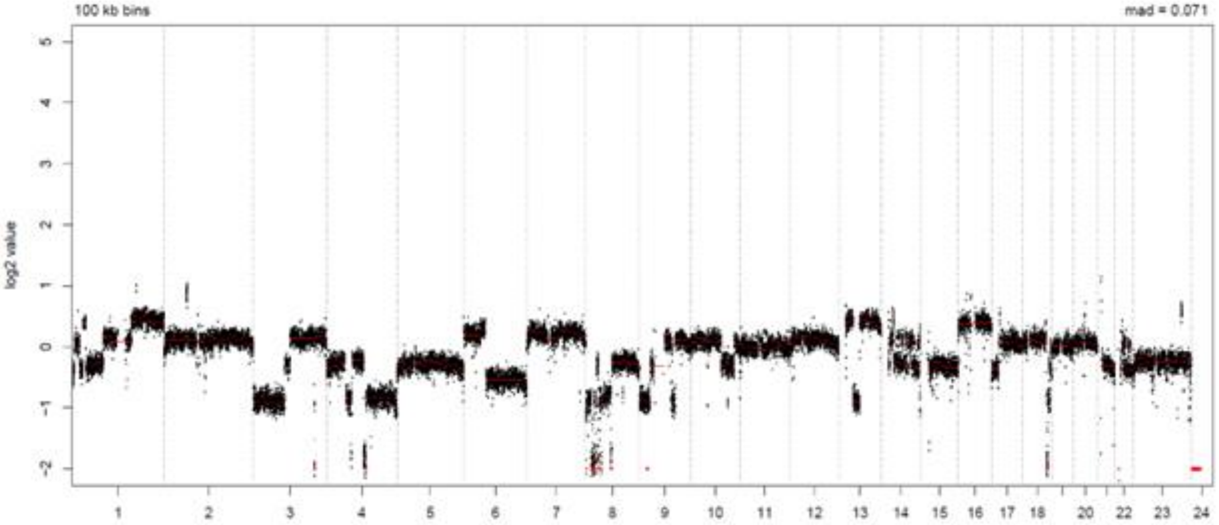


LymphGen Class (Confidence): EZB

BN2: 0.01    EZB: 0.99    N1: 0.07

MCD: 0.00    ST2: 0.00    A53: 0.57

### LPMI386

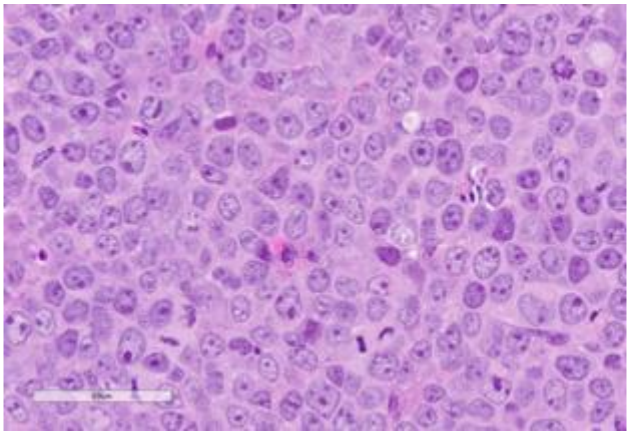

Source: MDACC

**Clinical Information**

Originating Biopsy Diagnosis: HGBCL

Prior Indolent Lymphoma: Follicular Lymphoma

Sex: Female

Race: Other

Ethnicity: Hispanic or Latino

Age at diagnosis: 46

Age at PDX biopsy: 52

### LoTs Pre Biopsy (best response): 10 (CR)

### LoTs Post Biopsy (best response): 0 (PD)

Therapies: Chemotherapy; Chemotherapy; Chemotherapy; Targeted; Other; CART; Targeted; Bispecific; Targeted; Chemotherapy

**Immunohistochemical Staining:**

Hans COO call: GCB

BCL6 (+)

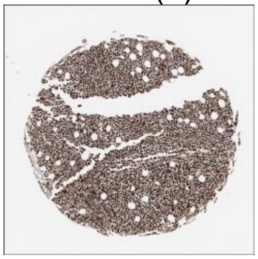

CD10 (+)

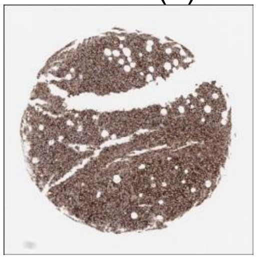

MUM1 (-)

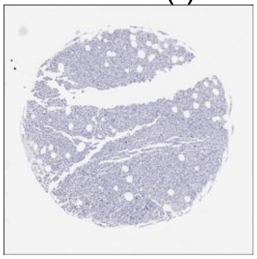

MYC (+)

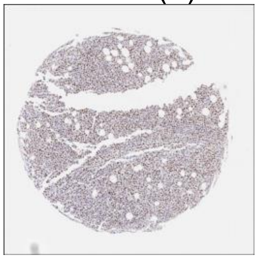

CD20 (-)

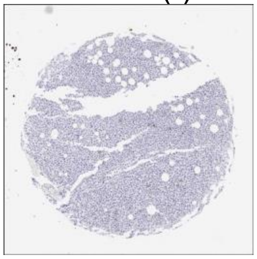

CD19 (-)

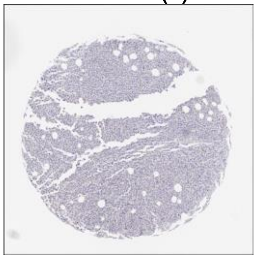

**DLBCL90 call:** GCB, DZsig+

**Genetic Characteristics:**

BCL2 FISH: Positive    MYC FISH: Negative    BCL6 FISH: Negative

Copy Number Alterations:

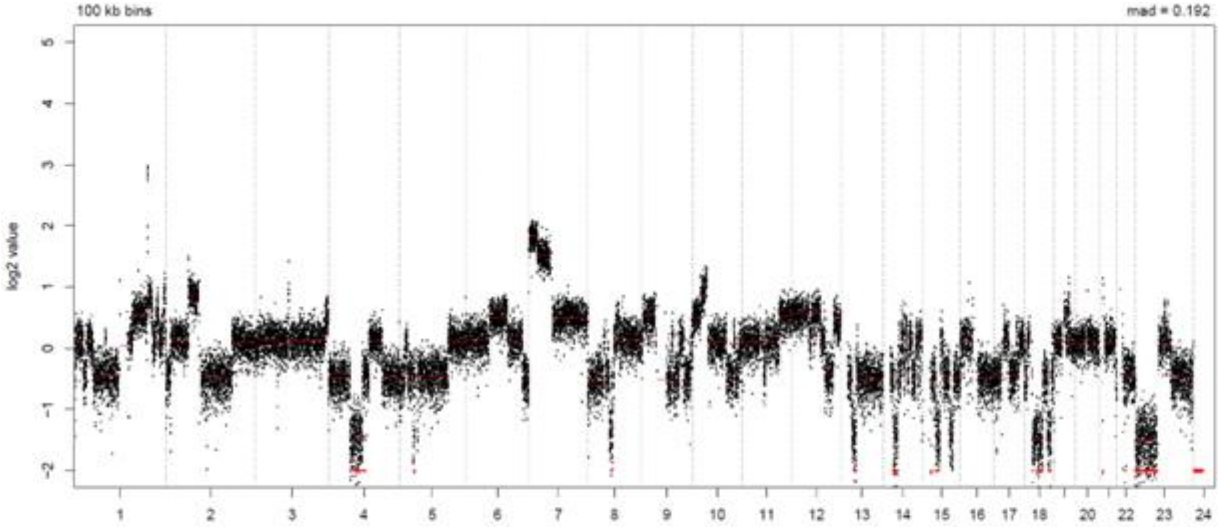


LymphGen Class (Confidence): EZB

BN2: 0.02    EZB: 0.99    N1: 0.07
MCD: 0.00    ST2: 0.16    A53: 0.05

### LPMI395

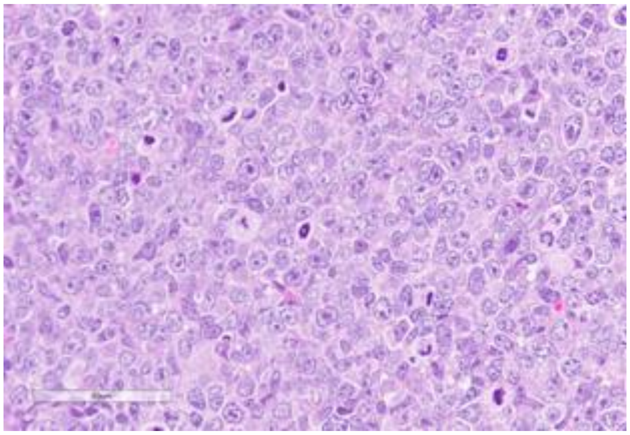

Source: MDACC

**Clinical Information**

Originating Biopsy Diagnosis: DLBCL, NOS

Prior Indolent Lymphoma: None

Sex: Male

Race: White or Caucasian

Ethnicity: Not Hispanic or Latino

Age at diagnosis: 71

Age at PDX biopsy: 71

### LoTs Pre Biopsy (best response): 2 (PR)

### LoTs Post Biopsy (best response): 1 (PR)

Therapies: Chemotherapy; RT; CART

**Immunohistochemical Staining:**

Hans COO call: Non-GCB

BCL6 (-)

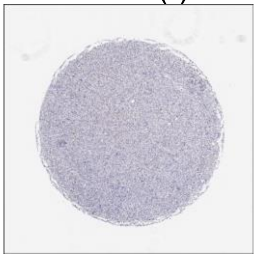

CD10 (-)

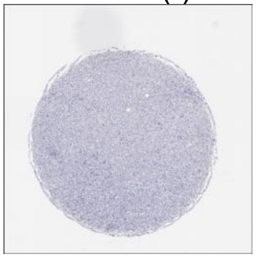

MUM1 (+)

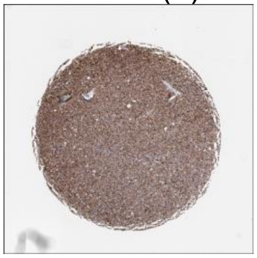

MYC (+)

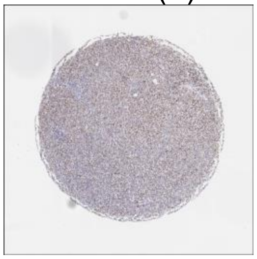

CD20 (+)

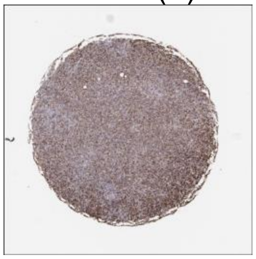

CD19 (+)

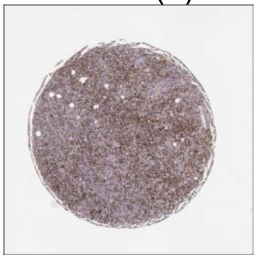

**DLBCL90 call:** ABC

**Genetic Characteristics:**

| | | |
|---|---|---|
| BCL2 FISH: Negative | MYC FISH: Positive | BCL6 FISH: Negative |

Copy Number Alterations:

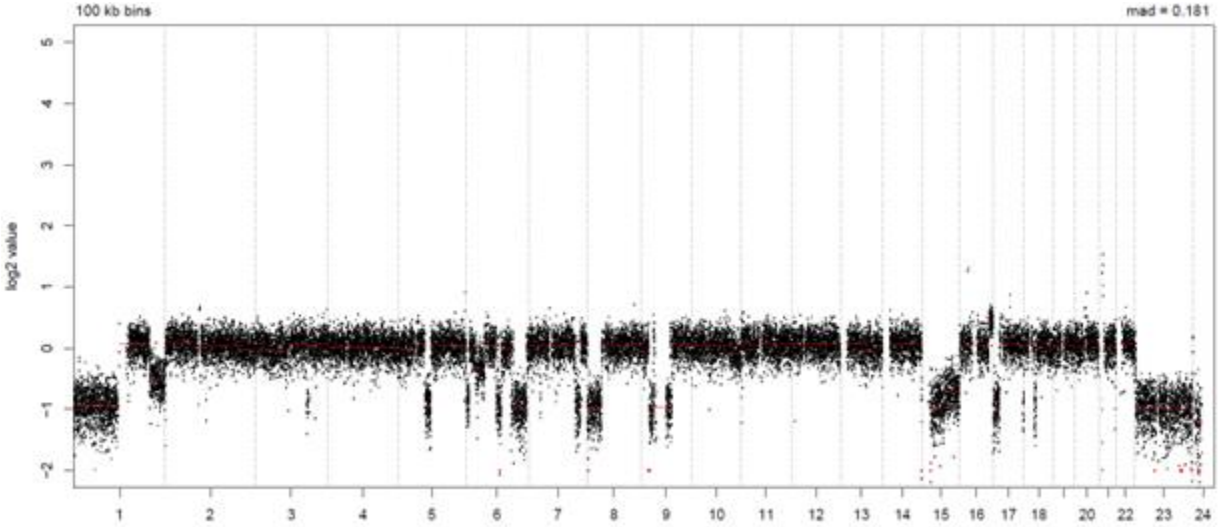


LymphGen Class (Confidence): Other

| | | |
|---|---|---|
| BN2: 0.04 | EZB: 0.26 | N1: 0.07 |
| MCD: 0.00 | ST2: 0.04 | A53: 0.01 |

### LPMI396

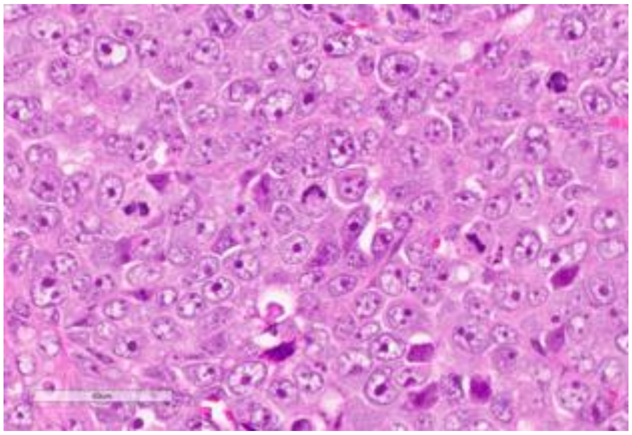

Source: MDACC

**Clinical Information**

Originating Biopsy Diagnosis: DLBCL, NOS

Prior Indolent Lymphoma: CLL/SLL

Sex: Male

Race: White or Caucasian

Ethnicity: Not Hispanic or Latino

Age at diagnosis: 74

Age at PDX biopsy: 75

### LoTs Pre Biopsy (best response): 3 (PR)

### LoTs Post Biopsy (best response): 2 (PD)

Therapies: Chemotherapy; Chemotherapy; Targeted; Chemotherapy; CART

**Immunohistochemical Staining:**

Hans COO call: Non-GCB

BCL6 (-)

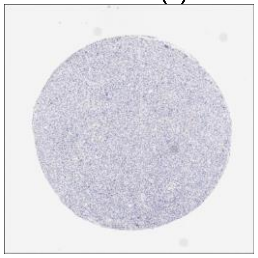

CD10 (-)

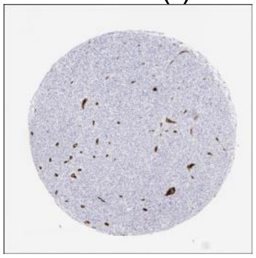

MUM1 (+)

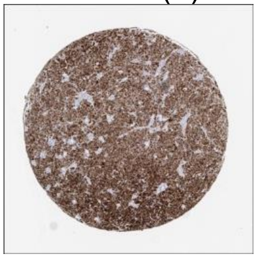

MYC (+)

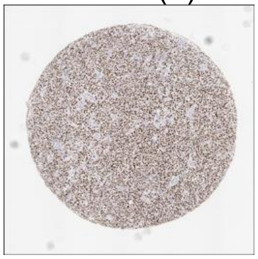

CD20 (+)

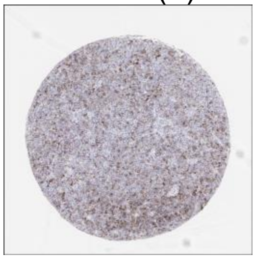

CD19 (-)

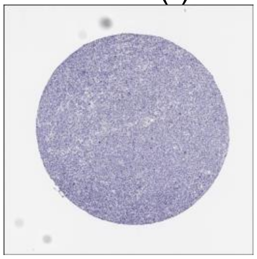

**DLBCL90 call:** ABC

**Genetic Characteristics:**

BCL2 FISH: Negative

MYC FISH: Positive

BCL6 FISH: Negative

Copy Number Alterations:

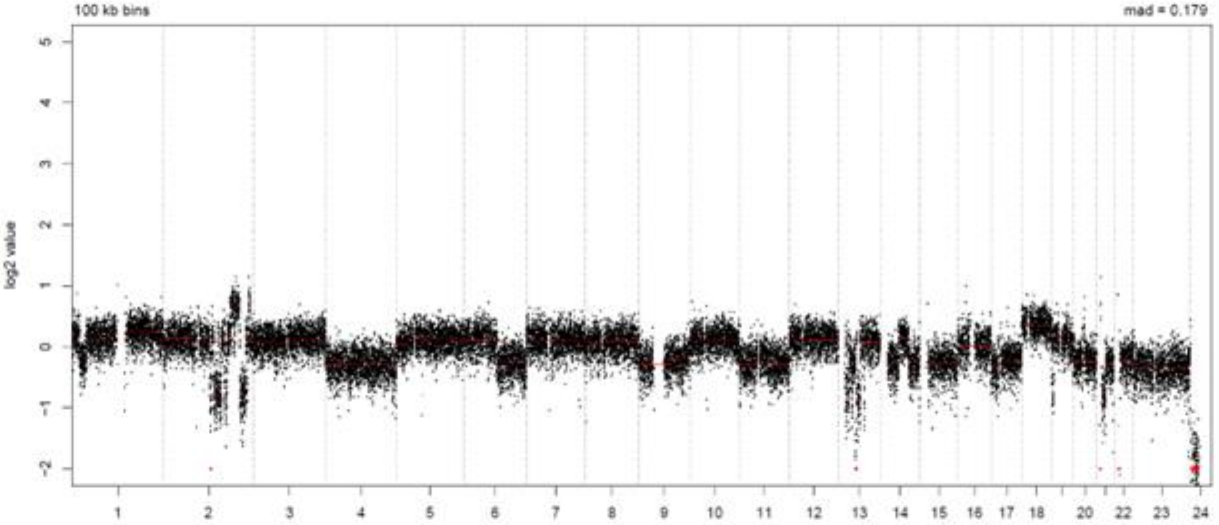


LymphGen Class (Confidence): ST2/A53

BN2: 0.12

EZB: 0.00

N1: 0.07

MCD: 0.03

ST2: 0.93

A53: 0.90

### LPMI406

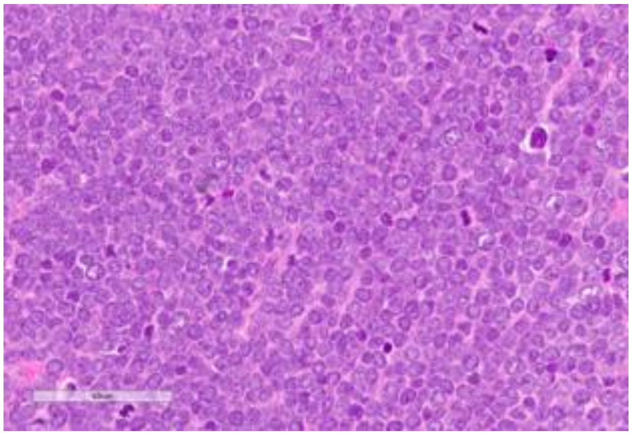

Source: MDACC

**Clinical Information**

<u>Originating Biopsy Diagnosis</u>: HGBCL

<u>Prior Indolent Lymphoma</u>: Follicular Lymphoma

<u>Sex</u>: Female

<u>Race</u>: Other

<u>Ethnicity</u>: Not Hispanic or Latino

<u>Age at diagnosis</u>: 54

<u>Age at PDX biopsy</u>: 55

<u># LoTs Pre Biopsy</u> (best response): 4 (CR)

<u># LoTs Post Biopsy</u> (best response): 2 (PD)

<u>Therapies</u>: Chemotherapy; Targeted; Chemotherapy; CART; Targeted; Chemotherapy

**Immunohistochemical Staining:**

Hans COO call: GCB

BCL6 (-)

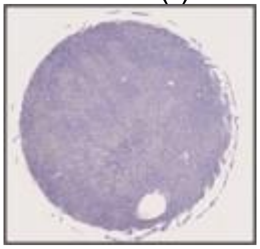

CD10 (+)

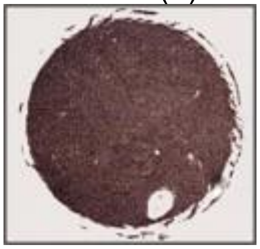

MUM1 (+)

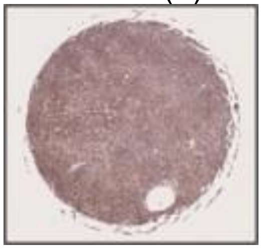

MYC (+)

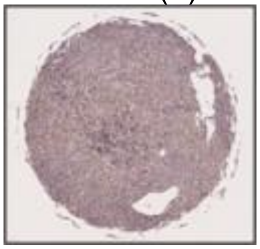

CD20 (+)

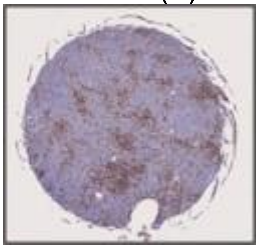

CD19 (+)

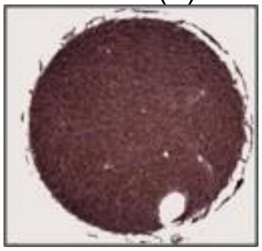

**DLBCL90 call:** ABC

**Genetic Characteristics:**

BCL2 FISH: Positive

MYC FISH: Positive

BCL6 FISH: Negative

Copy Number Alterations:

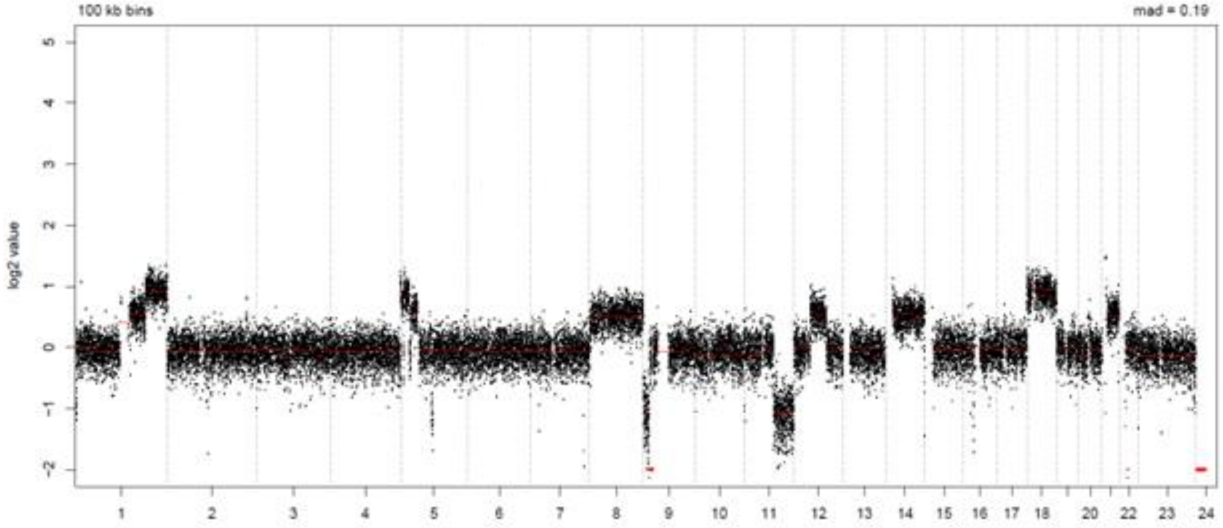


LymphGen Class (Confidence): EZB

BN2: 0.01

EZB: 0.99

N1: 0.07

MCD: 0.00

ST2: 0.00

A53: 0.00

### LPMI408

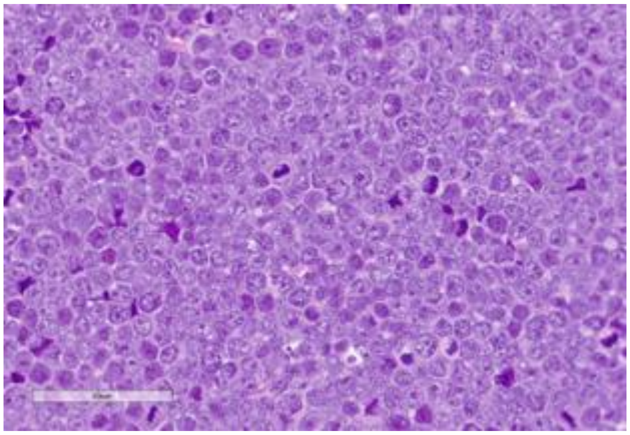

Source: MDACC

**Clinical Information**

<u>Originating Biopsy Diagnosis</u>: HGBCL

<u>Prior Indolent Lymphoma</u>: Follicular Lymphoma

<u>Sex</u>: Male

<u>Race</u>: Other

<u>Ethnicity</u>: Not Hispanic or Latino

<u>Age at diagnosis</u>: 51

<u>Age at PDX biopsy</u>: 52

<u># LoTs Pre Biopsy</u> (best response): 3 (CR)

<u># LoTs Post Biopsy</u> (best response): 3 (PD)

<u>Therapies</u>: Chemotherapy; Chemotherapy; CART; Chemotherapy; Targeted; Chemotherapy

**Immunohistochemical Staining:**

Hans COO call: GCB

BCL6 (-)

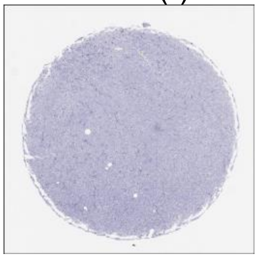

CD10 (+)

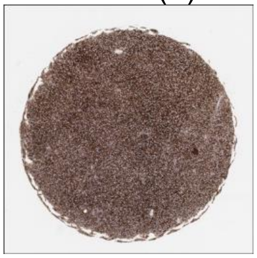

MUM1 (+)

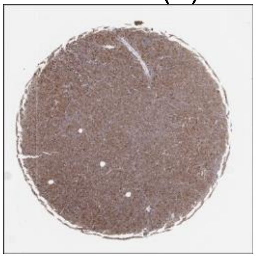

MYC (+)

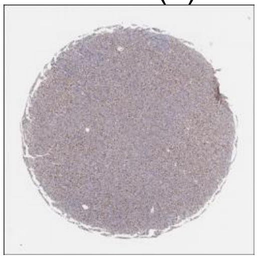

CD20 (+)

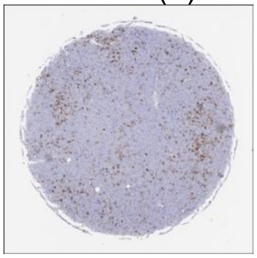

CD19 (+)

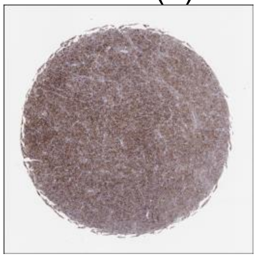

**DLBCL90 call:** ABC

**Genetic Characteristics:**

BCL2 FISH: Positive | MYC FISH: Negative | BCL6 FISH: Negative

Copy Number Alterations:

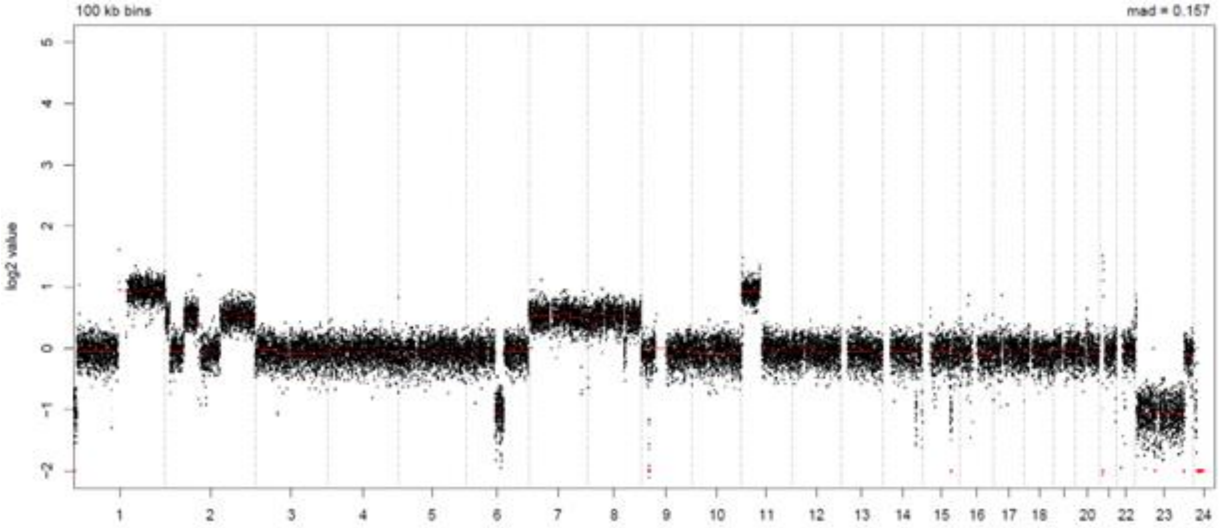


LymphGen Class (Confidence): EZB

BN2: 0.01
EZB: 0.99
N1: 0.07
MCD: 0.00
ST2: 0.00
A53: 0.00

### LPMI411

Source: MDACC

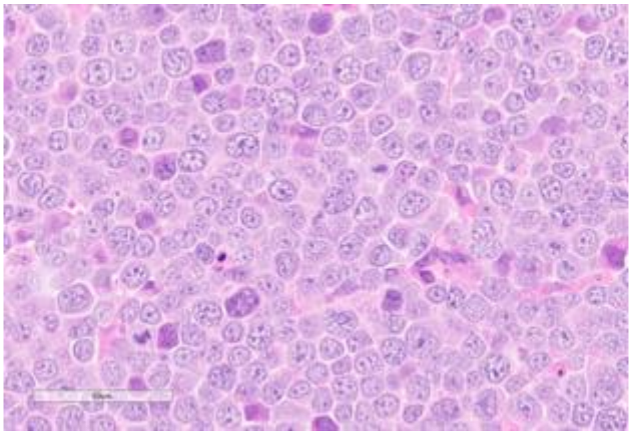

**Clinical Information**

Originating Biopsy Diagnosis: DLBCL, ABC

Prior Indolent Lymphoma: None

Sex: Male

Race: White or Caucasian

Ethnicity: Not Hispanic or Latino

Age at diagnosis: 39

Age at PDX biopsy: 41

### LoTs Pre Biopsy (best response): 5 (PR)

### LoTs Post Biopsy (best response): 4 (PD)

Therapies: Chemotherapy; Chemotherapy; Chemotherapy; Chemotherapy; CART; Targeted; Targeted; Chemotherapy; Targeted

**Immunohistochemical Staining:**

Hans COO call: Non-GCB

BCL6 (+)

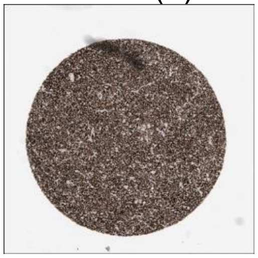

CD10 (-)

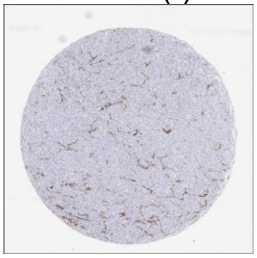

MUM1 (+)

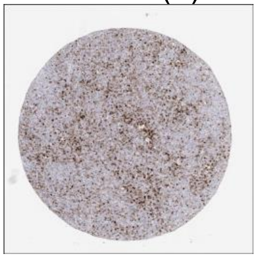

MYC (+)

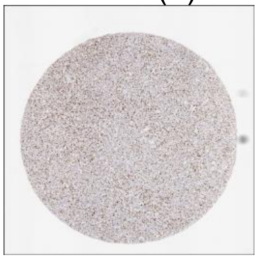

CD20 (+)

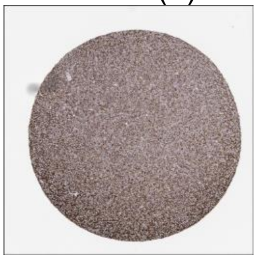

CD19 (+)

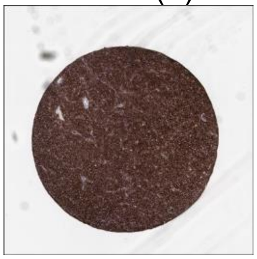

**DLBCL90 call:** UNCLASS

**Genetic Characteristics:**

BCL2 FISH: Negative MYC FISH: Negative BCL6 FISH: Negative

Copy Number Alterations:

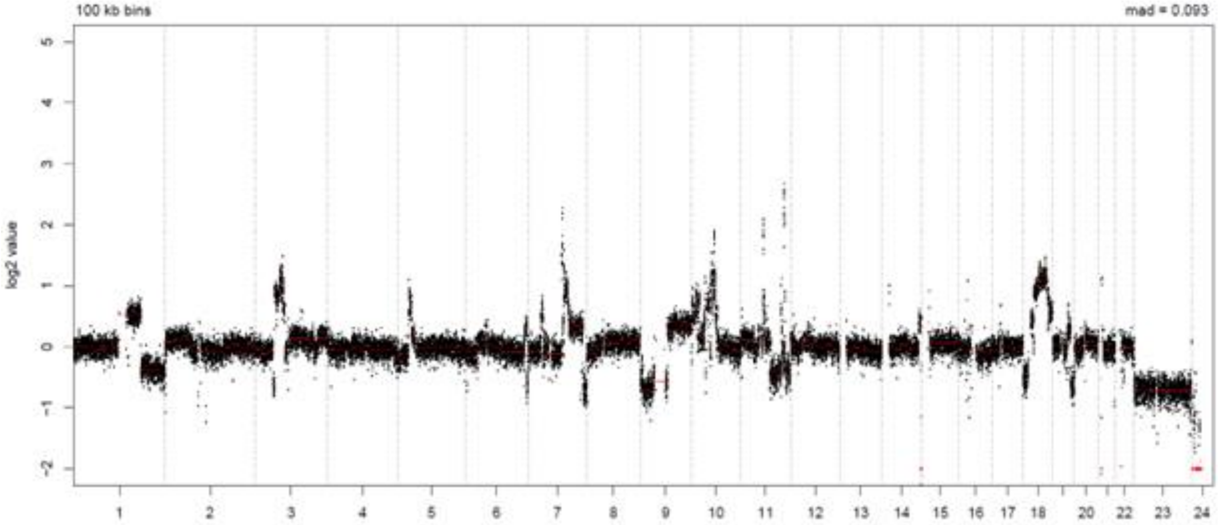


LymphGen Class (Confidence): Other

| | | |
|---|---|---|
| BN2: 0.07 | EZB: 0.08 | N1: 0.07 |
| MCD: 0.00 | ST2: 0.44 | A53: 0.00 |

### LPMI414

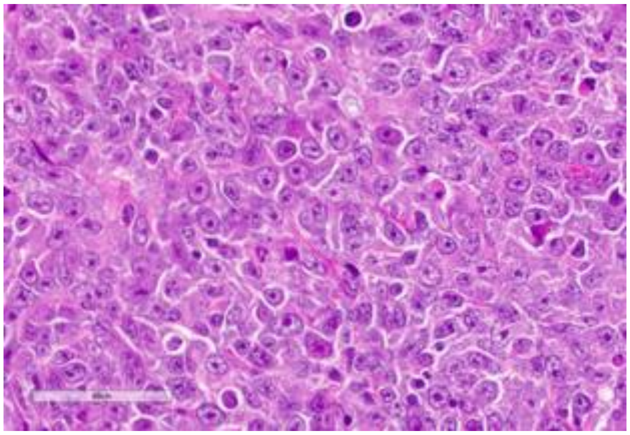

Source: MDACC

**Clinical Information**

Originating Biopsy Diagnosis: DLBCL, GCB

Prior Indolent Lymphoma: Follicular Lymphoma

Sex: Male

Race: White or Caucasian

Ethnicity: Hispanic or Latino

Age at diagnosis: 65

Age at PDX biopsy: 71

### LoTs Pre Biopsy (best response): 3 (PD)

### LoTs Post Biopsy (best response): 1 (CR)

Therapies: Chemotherapy; Chemotherapy; CART; Targeted

**Immunohistochemical Staining:**

Hans COO call: GCB

BCL6 (+)

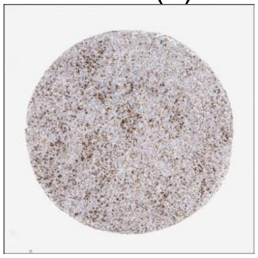

CD10 (+)

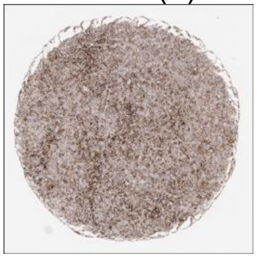

MUM1 (+)

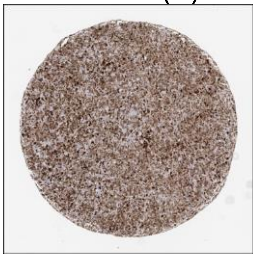

MYC (+)

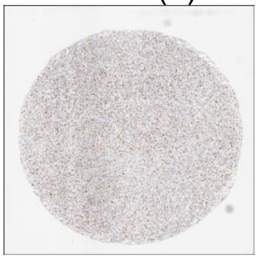

CD20 (+)

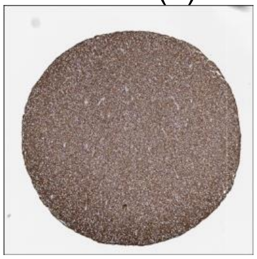

CD19 (+)

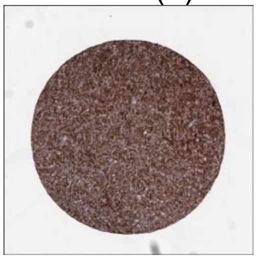

**DLBCL90 call:** ABC

**Genetic Characteristics:**

BCL2 FISH: Positive | MYC FISH: Positive | BCL6 FISH: Negative

Copy Number Alterations:

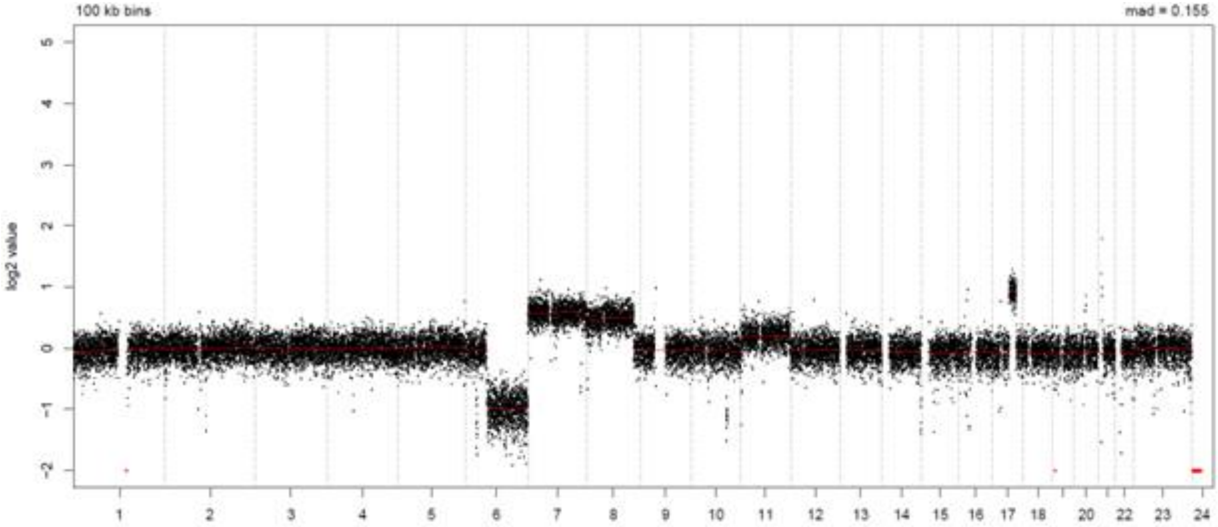


LymphGen Class (Confidence): EZB

BN2: 0.22
EZB: 0.90
N1: 0.07
MCD: 0.00
ST2: 0.03
A53: 0.00

### LPMI424

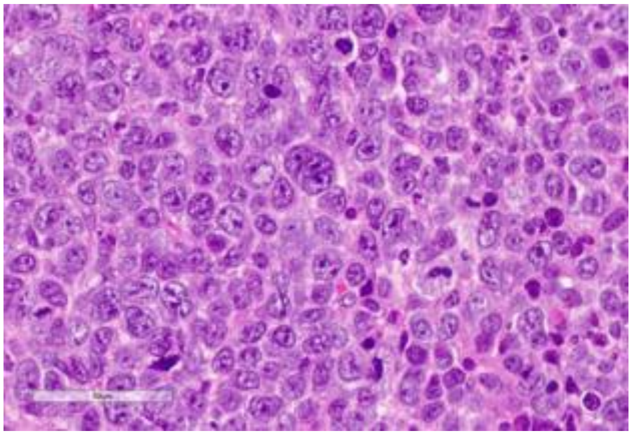

Source: MDACC

**Clinical Information**

<u>Originating Biopsy Diagnosis</u>: DLBCL, ABC

<u>Prior Indolent Lymphoma</u>: None

<u>Sex</u>: Male

<u>Race</u>: Other

<u>Ethnicity</u>: Hispanic or Latino

<u>Age at diagnosis</u>: 64

<u>Age at PDX biopsy</u>: 66

<u># LoTs Pre Biopsy</u> (best response): 5 (CR)

<u># LoTs Post Biopsy</u> (best response): 4 (CR)

<u>Therapies</u>: Chemotherapy; Chemotherapy; Chemotherapy; Chemotherapy; Chemotherapy; CART; CART; Targeted; Targeted

**Immunohistochemical Staining:**

Hans COO call: Non-GCB

BCL6 (-)

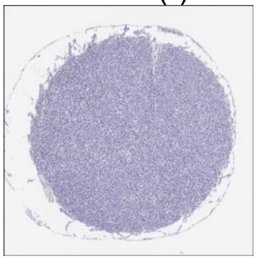

CD10 (-)

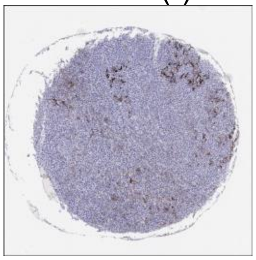

MUM1 (+)

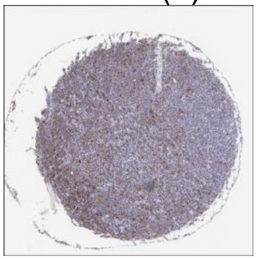

MYC (-)

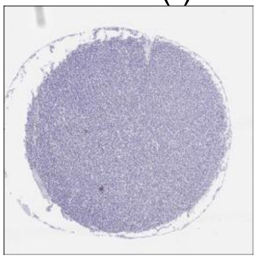

CD20 (+)

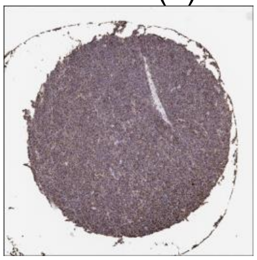

CD19 (+)

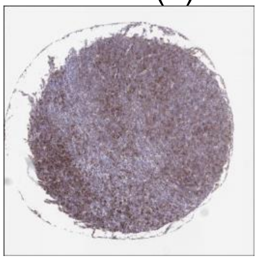

**DLBCL90 call:** ABC

**Genetic Characteristics:**

BCL2 FISH: Negative MYC FISH: Negative BCL6 FISH: Negative

Copy Number Alterations:

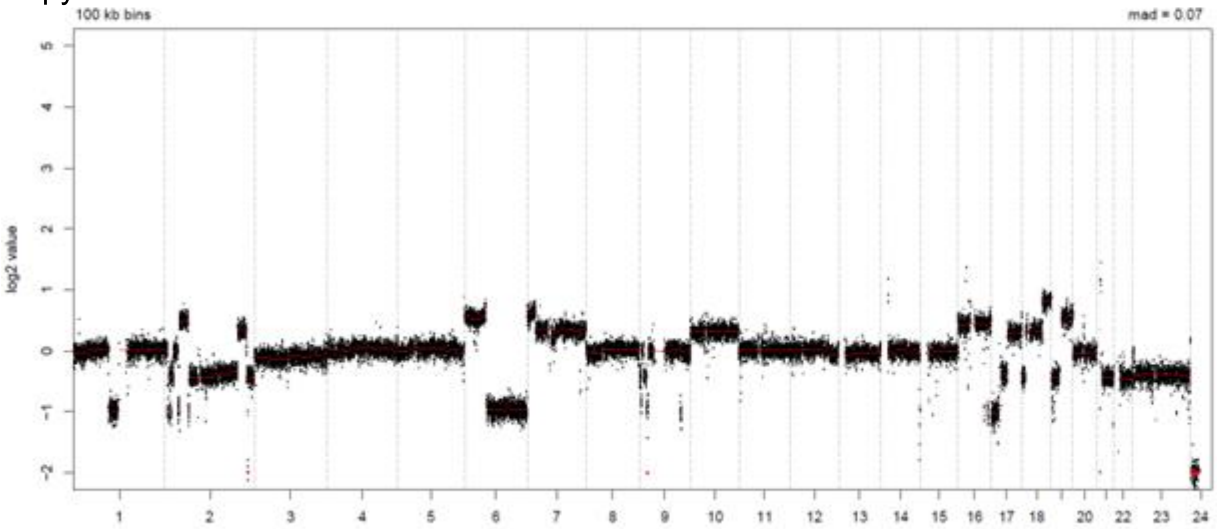


LymphGen Class (Confidence): A53

BN2: 0.01 EZB: 0.00 N1: 0.07

MCD: 0.09 ST2: 0.01 A53: 0.82

### LPMI431

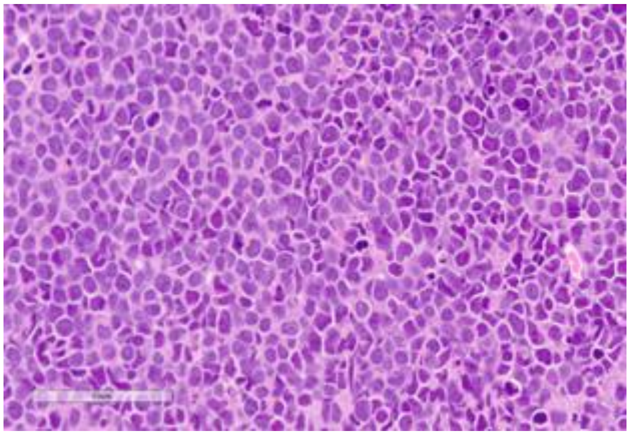

Source: MDACC

**Clinical Information**

Originating Biopsy Diagnosis: DLBCL, ABC

Prior Indolent Lymphoma: None

Sex: Female

Race: Asian

Ethnicity: Not Hispanic or Latino

Age at diagnosis: 66

Age at PDX biopsy: 67

### LoTs Pre Biopsy (best response): 4 (CR)

### LoTs Post Biopsy (best response): 1 (PD)

Therapies: Chemotherapy; Chemotherapy; Chemotherapy; CART; Targeted

**Immunohistochemical Staining:**

Hans COO call: GCB

BCL6 (-)

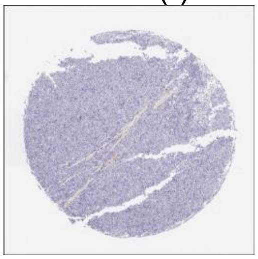

CD10 (+)

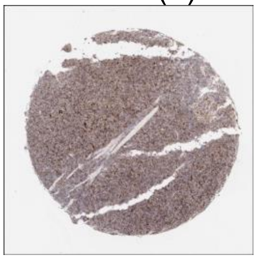

MUM1 (+)

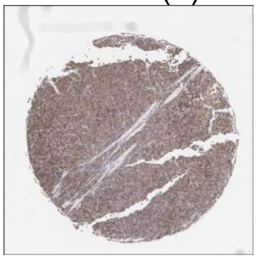

MYC (+)

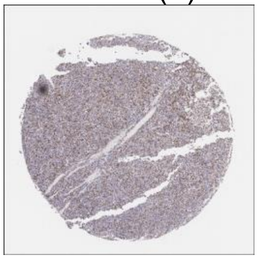

CD20 (+)

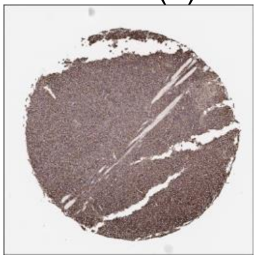

CD19 (+)

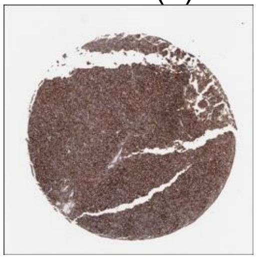

**DLBCL90 call:** ABC

**Genetic Characteristics:**

BCL2 FISH: Negative

MYC FISH: Negative

BCL6 FISH: Negative

Copy Number Alterations:

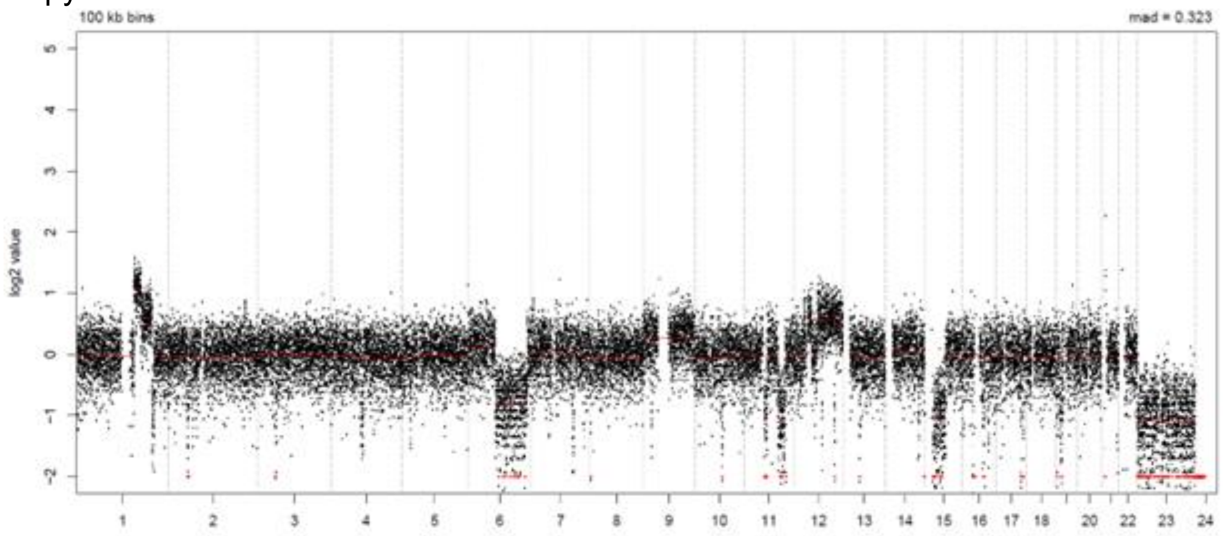


LymphGen Class (Confidence): Other

BN2: 0.02

EZB: 0.02

N1: 0.07

MCD: 0.20

ST2: 0.02

A53: 0.00

### LPMI462

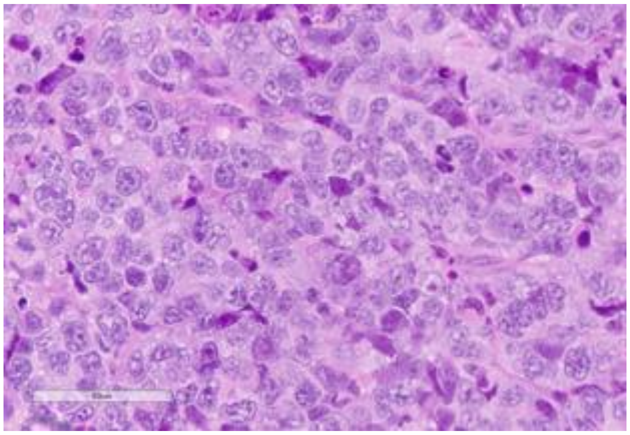

Source: MDACC

**Clinical Information**

Originating Biopsy Diagnosis: DLBCL, GCB

Prior Indolent Lymphoma: None

Sex: Female

Race: Other

Ethnicity: Hispanic or Latino

Age at diagnosis: 77

Age at PDX biopsy: 79

### LoTs Pre Biopsy (best response): 3 (SD)

### LoTs Post Biopsy (best response): 0 (SD)

Therapies: Chemotherapy; Chemotherapy; CART

**Immunohistochemical Staining:**

Hans COO call: GCB

BCL6 (+)

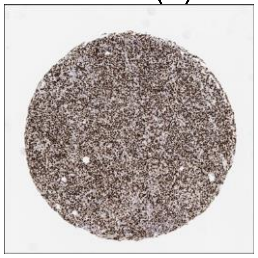

CD10 (+)

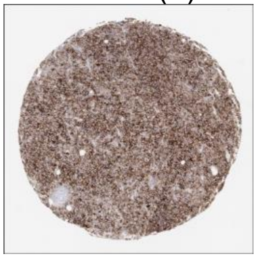

MUM1 (+)

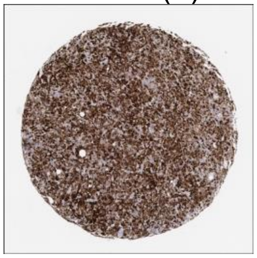

MYC (-)

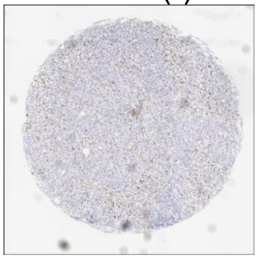

CD20 (+)

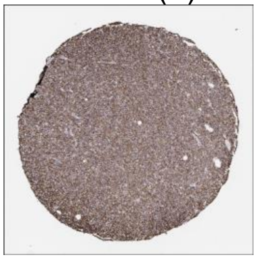

CD19 (+)

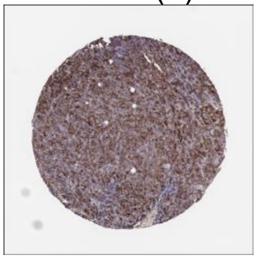

**DLBCL90 call:** GCB

**Genetic Characteristics:**

BCL2 FISH: Positive | MYC FISH: Positive | BCL6 FISH: Positive

Copy Number Alterations:

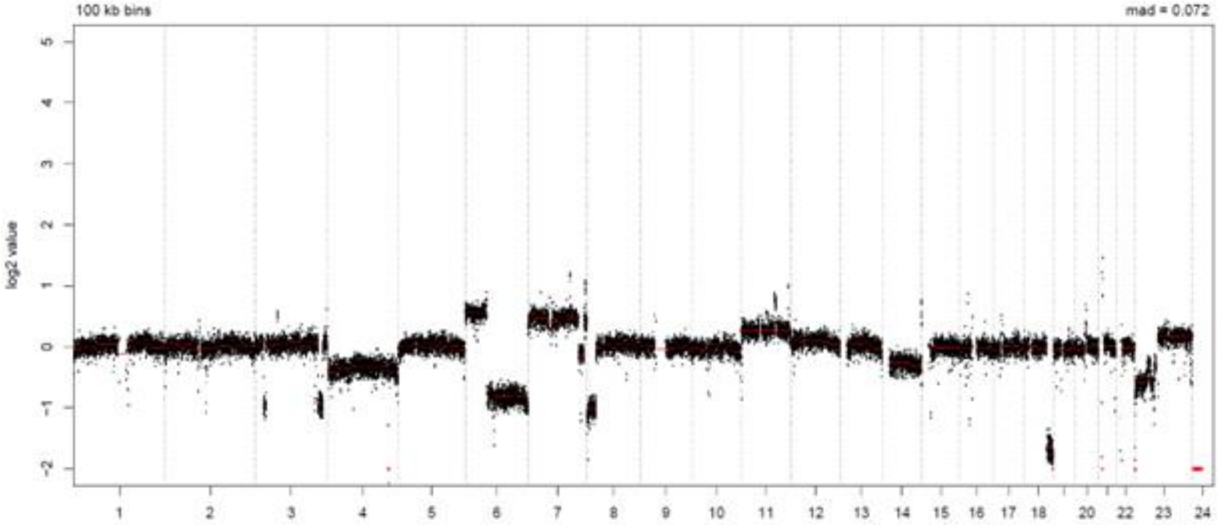


LymphGen Class (Confidence): EZB

BN2: 0.08 | EZB: 0.99 | N1: 0.07
MCD: 0.00 | ST2: 0.56 | A53: 0.00

### LPMI466

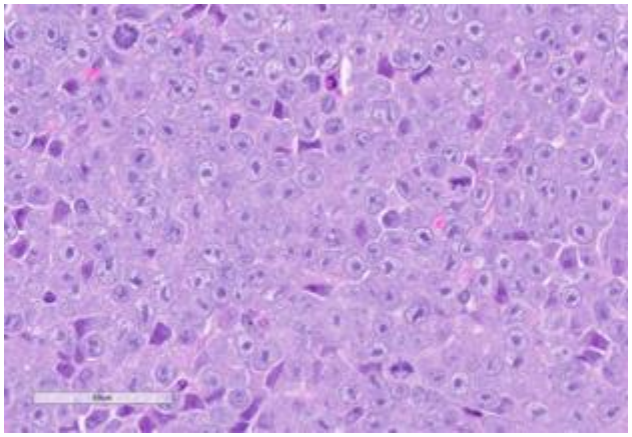

Source: MDACC

**Clinical Information**

Originating Biopsy Diagnosis: DLBCL, NOS

Prior Indolent Lymphoma: None

Sex: Male

Race: Asian

Ethnicity: Not Hispanic or Latino

Age at diagnosis: 51

Age at PDX biopsy: 57

### LoTs Pre Biopsy (best response): 7 (PD)

### LoTs Post Biopsy (best response): 2 (PD)

Therapies: Chemotherapy; CART; Transplant; Targeted; Targeted; Chemotherapy; Targeted; Targeted; Targeted

**Immunohistochemical Staining:**

Hans COO call: Non-GCB

BCL6 (-)

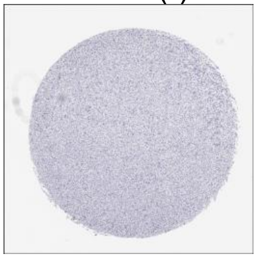

CD10 (-)

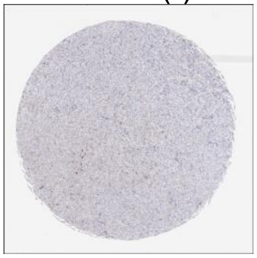

MUM1 (+)

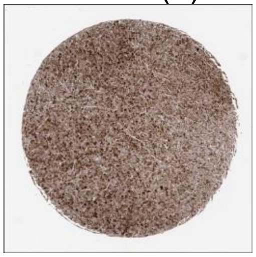

MYC (+)

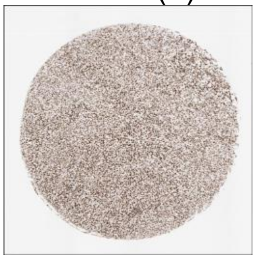

CD20 (+)

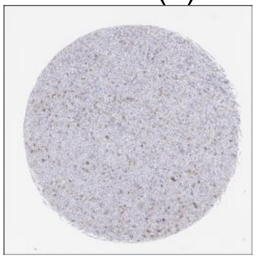

CD19 (-)

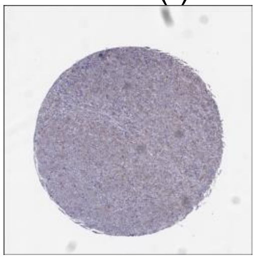

**DLBCL90 call:** ABC

**Genetic Characteristics:**

BCL2 FISH: Negative

MYC FISH: Positive

BCL6 FISH: Negative

Copy Number Alterations:

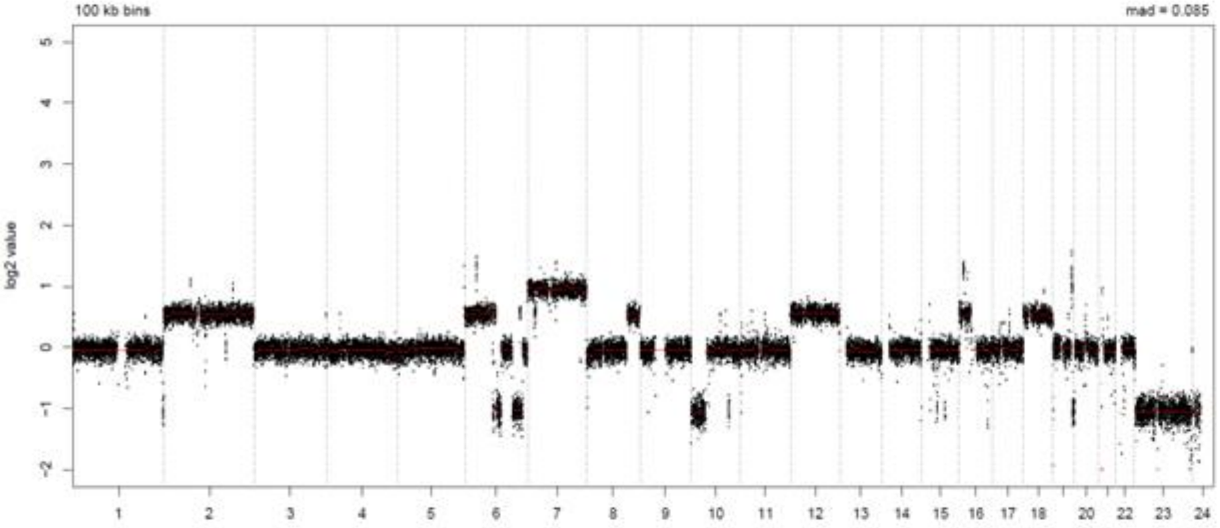


LymphGen Class (Confidence): MCD

| | | |
|---|---|---|
| BN2: 0.11 | EZB: 0.14 | N1: 0.07 |
| MCD: 0.88 | ST2: 0.03 | A53: 0.00 |

### LPMI467

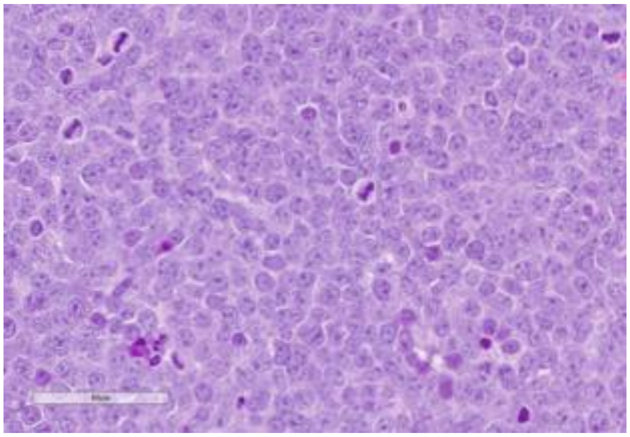

Source: MDACC

**Clinical Information**

Originating Biopsy Diagnosis: DLBCL, NOS

Prior Indolent Lymphoma: None

Sex: Female

Race: White or Caucasian

Ethnicity: Not Hispanic or Latino

Age at diagnosis: 66

Age at PDX biopsy: 68

### LoTs Pre Biopsy (best response): 5 (PD)

### LoTs Post Biopsy (best response): 2 (PD)

Therapies: Chemotherapy; CART; Targeted; RT; Other Cellular Therapy; RT; RT

**Immunohistochemical Staining:**

Hans COO call: Non-GCB

BCL6 (+)

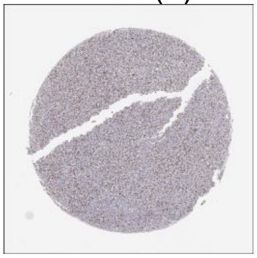

CD10 (-)

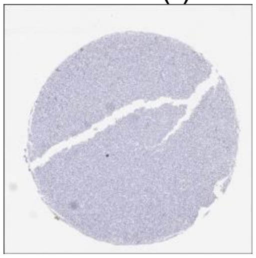

MUM1 (+)

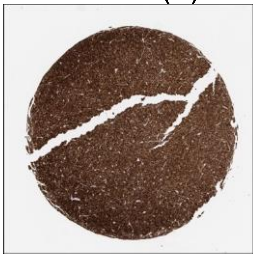

MYC (+)

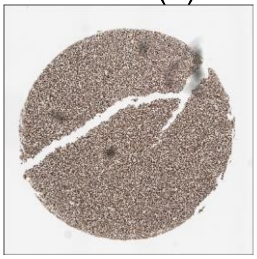

CD20 (+)

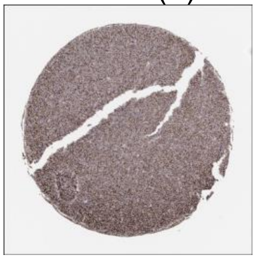

CD19 (-)

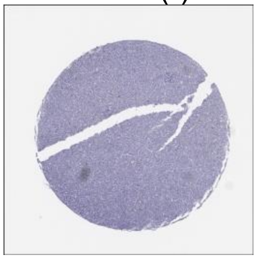

**DLBCL90 call:** ABC

**Genetic Characteristics:**

BCL2 FISH: Negative
MYC FISH: Abnormal
BCL6 FISH: Negative

Copy Number Alterations:

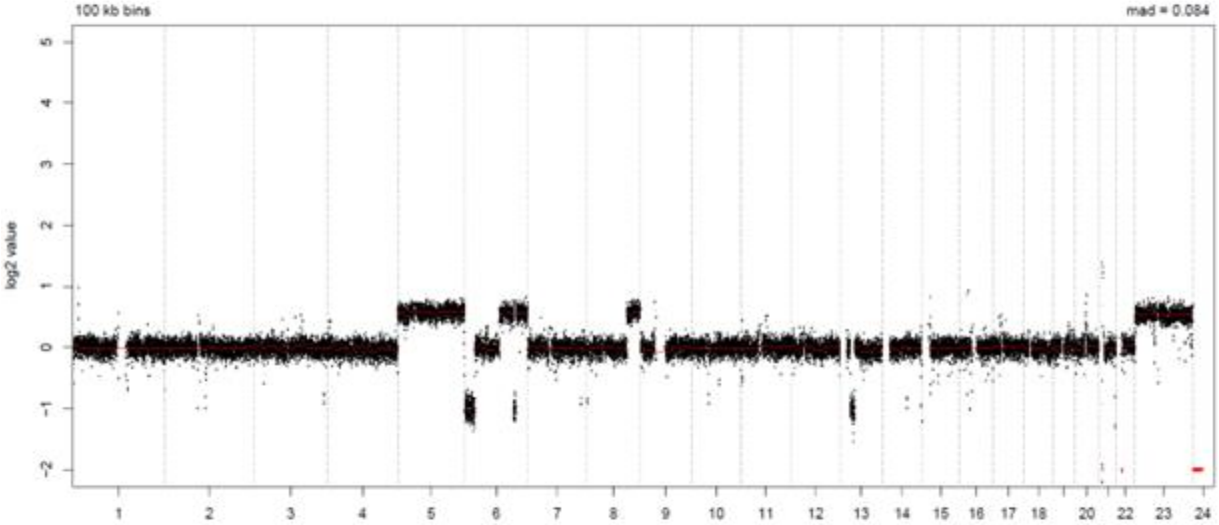


LymphGen Class (Confidence): ST2

| | | |
|---|---|---|
| BN2: 0.32 | EZB: 0.01 | N1: 0.07 |
| MCD: 0.05 | ST2: 0.87 | A53: 0.00 |

### LPMI478

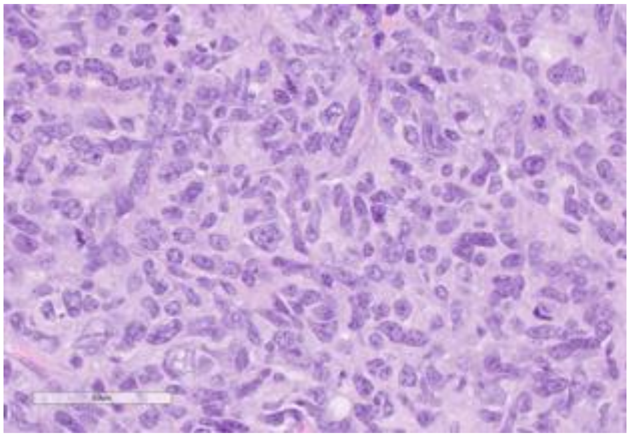

Source: MDACC

**Clinical Information**

Originating Biopsy Diagnosis: DLBCL, NOS

Prior Indolent Lymphoma: None

Sex: Male

Race: White or Caucasian

Ethnicity: Hispanic or Latino

Age at diagnosis: 49

Age at PDX biopsy: 49

### LoTs Pre Biopsy (best response): 1 (SD)

### LoTs Post Biopsy (best response): 2 (SD)

Therapies: Chemotherapy; Chemotherapy; CART

**Immunohistochemical Staining:**

Hans COO call: Non-GCB

BCL6 (+)

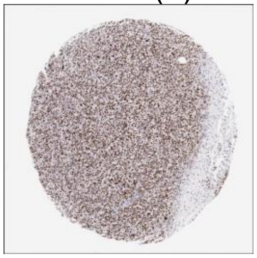

CD10 (-)

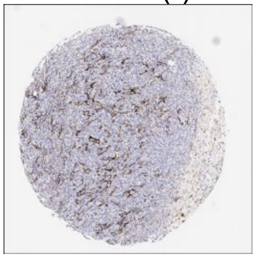

MUM1 (+)

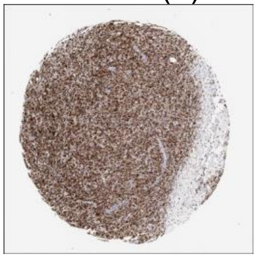

MYC (-)

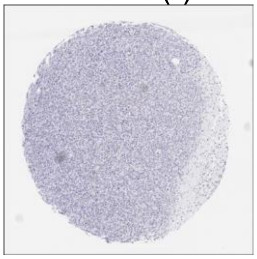

CD20 (+)

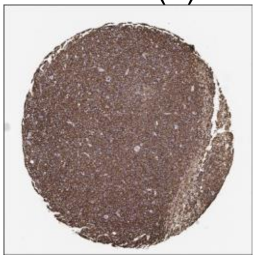

CD19 (+)

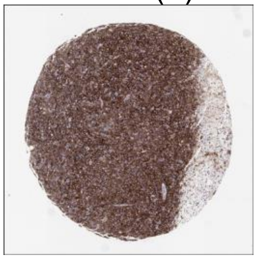

**DLBCL90 call:** ABC

**Genetic Characteristics:**

BCL2 FISH: Positive    MYC FISH: Negative    BCL6 FISH: Negative

Copy Number Alterations:

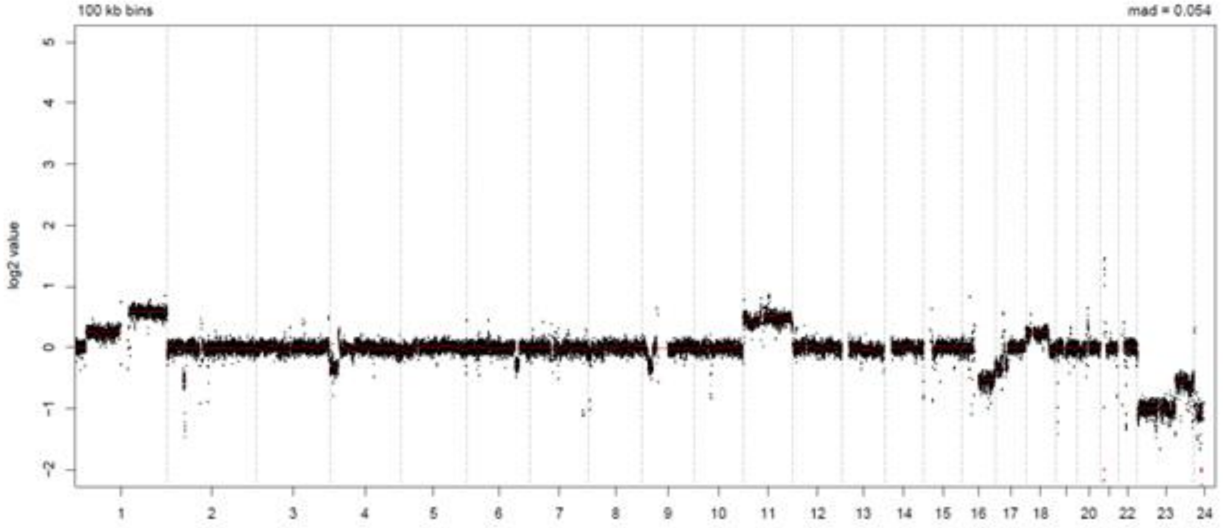


LymphGen Class (Confidence): EZB/ST2

BN2: 0.32    EZB: 0.99    N1: 0.07
MCD: 0.00    ST2: 0.99    A53: 0.00

### LPMI480

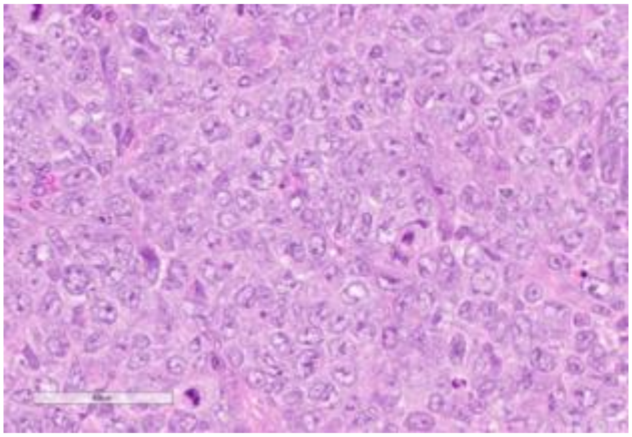

Source: MDACC

**Clinical Information**

Originating Biopsy Diagnosis: DLBCL, NOS

Prior Indolent Lymphoma: Follicular Lymphoma

Sex: Male

Race: Other

Ethnicity: Hispanic or Latino

Age at diagnosis: 43

Age at PDX biopsy: 56

### LoTs Pre Biopsy (best response): 6 (PD)

### LoTs Post Biopsy (best response): 0 (PR)

Therapies: Targeted; Targeted; Chemotherapy; Targeted; Targeted; Chemotherapy

**Immunohistochemical Staining:**

Hans COO call: GCB

BCL6 (+)

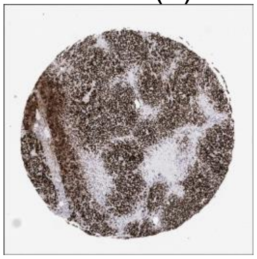

CD10 (+)

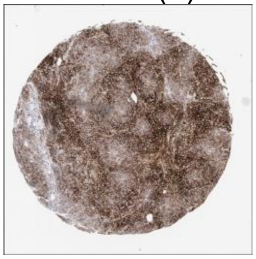

MUM1 (-)

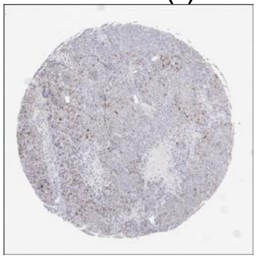

MYC (-)

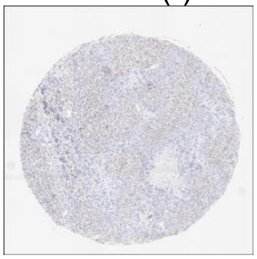

CD20 (+)

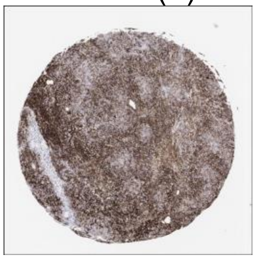

CD19 (+)

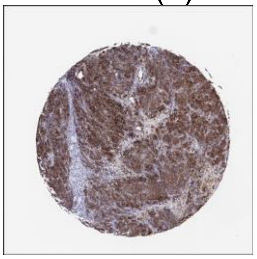

**DLBCL90 call:** GCB

**Genetic Characteristics:**

BCL2 FISH: Positive | MYC FISH: Negative | BCL6 FISH: Negative

Copy Number Alterations:

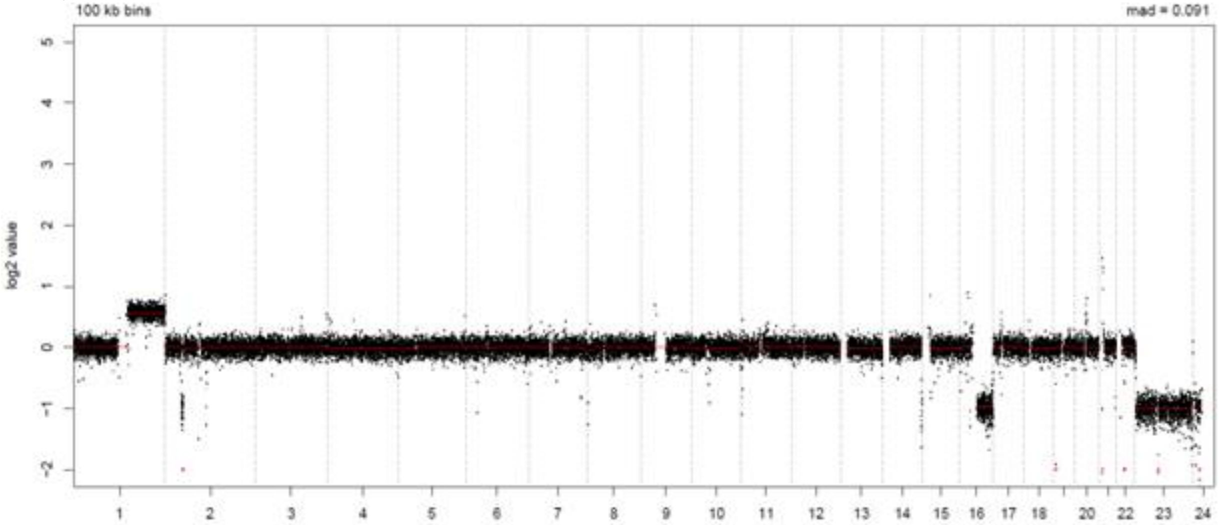


LymphGen Class (Confidence): BN2/ST2

BN2: 0.92
EZB: 0.86
N1: 0.07
MCD: 0.00
ST2: 0.99
A53: 0.00

### LPMI495

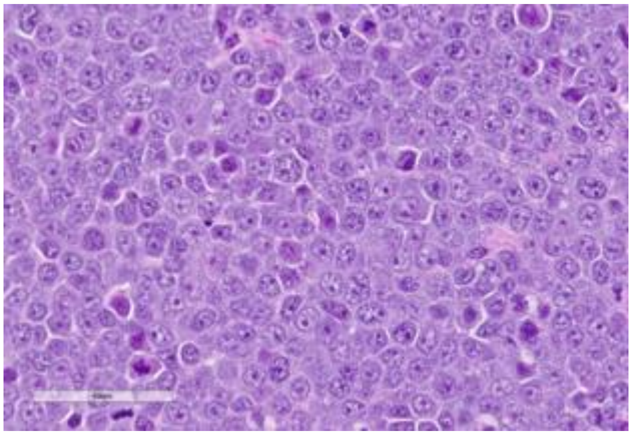

Source: MDACC

**Clinical Information**

Originating Biopsy Diagnosis: HGBCL

Prior Indolent Lymphoma: None

Sex: Male

Race: White or Caucasian

Ethnicity: Not Hispanic or Latino

Age at diagnosis: 75

Age at PDX biopsy: 75

### LoTs Pre Biopsy (best response): 1 (PR)

### LoTs Post Biopsy (best response): 7 (PD)

Therapies: Chemotherapy; Chemotherapy; CART; Chemotherapy; Immunotherapy; RT; Targeted; Targeted

**Immunohistochemical Staining:**

Hans COO call: GCB

BCL6 (-)

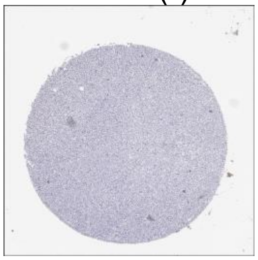

CD10 (+)

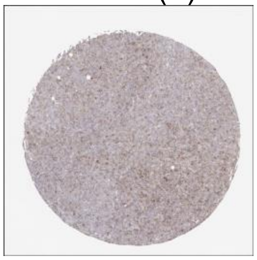

MUM1 (+)

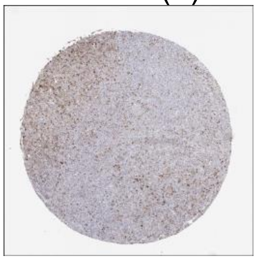

MYC (+)

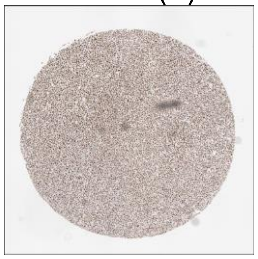

CD20 (+)

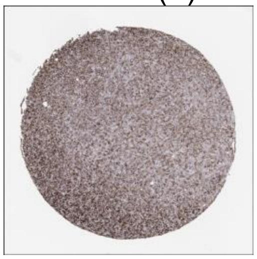

CD19 (+)

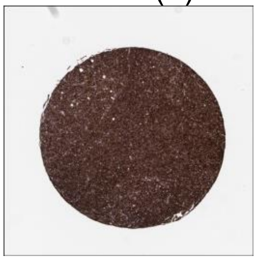

**DLBCL90 call:** ABC

**Genetic Characteristics:**

BCL2 FISH: Positive | MYC FISH: Positive | BCL6 FISH: Negative

Copy Number Alterations:

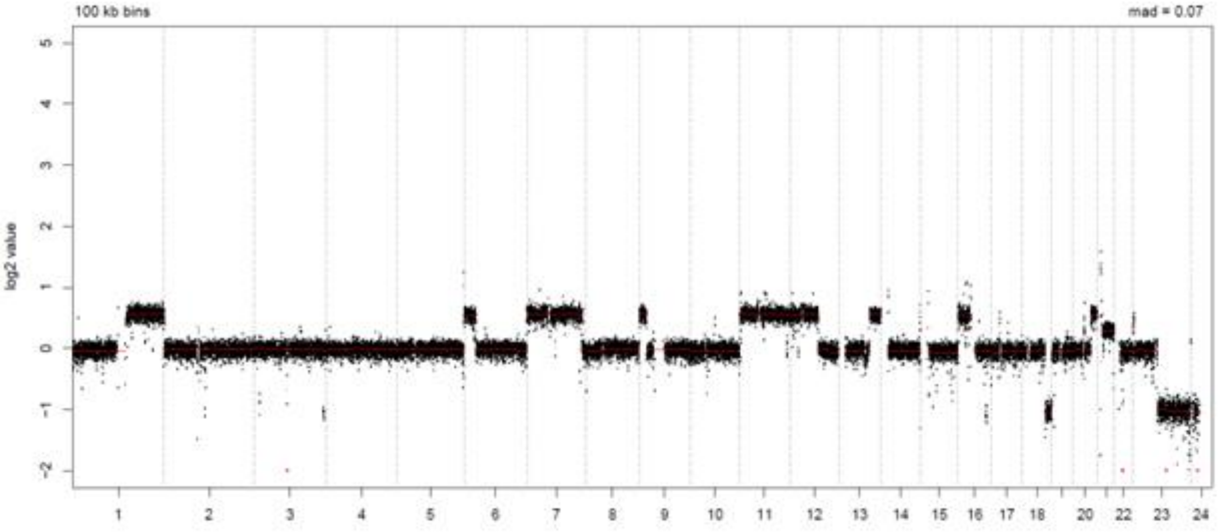


LymphGen Class (Confidence): EZB

BN2: 0.07 | EZB: 0.92 | N1: 0.07
MCD: 0.00 | ST2: 0.15 | A53: 0.00

### LPMI530

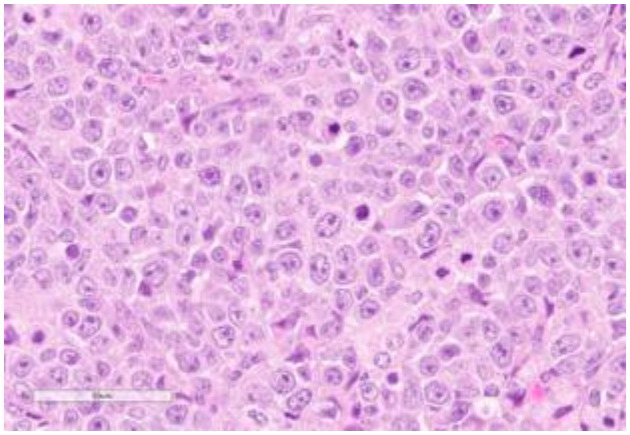

Source: MDACC

**Clinical Information**

Originating Biopsy Diagnosis: DLBCL, NOS

Prior Indolent Lymphoma: Follicular Lymphoma

Sex: Female

Race: Other

Ethnicity: Hispanic or Latino

Age at diagnosis: 54

Age at PDX biopsy: 58

### LoTs Pre Biopsy (best response): 1 (CR)

### LoTs Post Biopsy (best response): 1 (CR)

Therapies: Chemotherapy; Chemotherapy

**Immunohistochemical Staining:**

Hans COO call: Non-GCB

BCL6 (+)

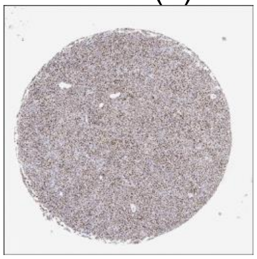

CD10 (-)

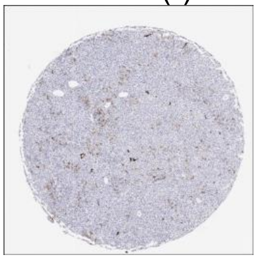

MUM1 (+)

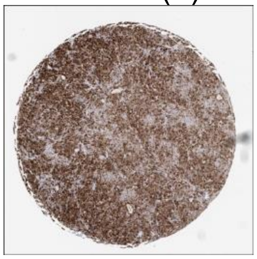

MYC (-)

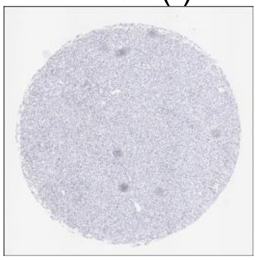

CD20 (+)

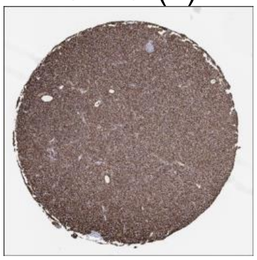

CD19 (+)

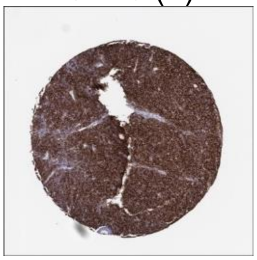

**DLBCL90 call:** ABC

**Genetic Characteristics:**

BCL2 FISH: Positive MYC FISH: Negative BCL6 FISH: Positive

Copy Number Alterations:

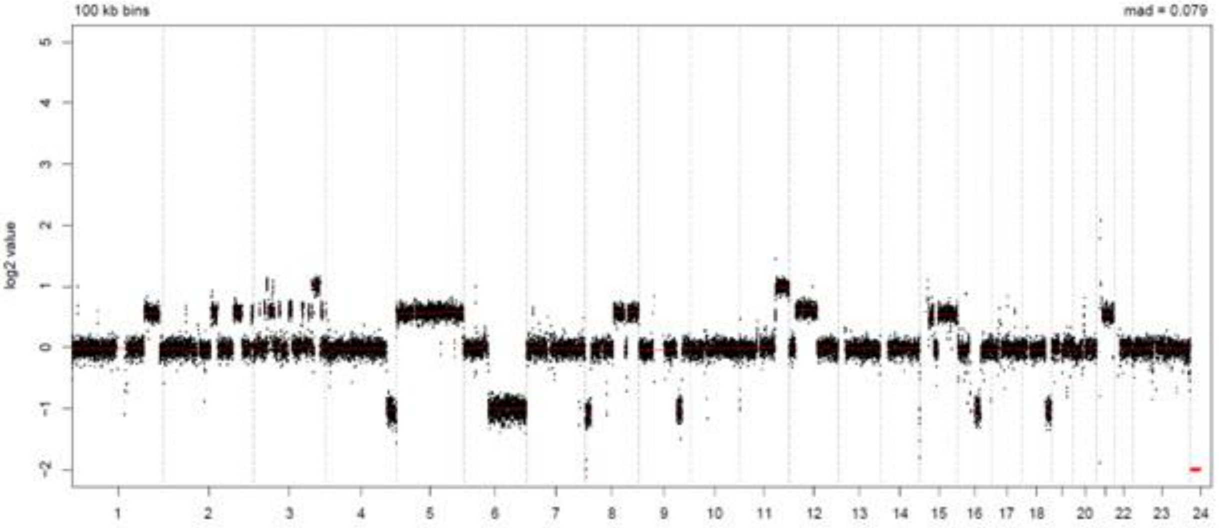


LymphGen Class (Confidence): Other

| | | |
|---|---|---|
| BN2: 0.54 | EZB: 0.63 | N1: 0.07 |
| MCD: 0.00 | ST2: 0.00 | A53: 0.00 |

### LPMI550

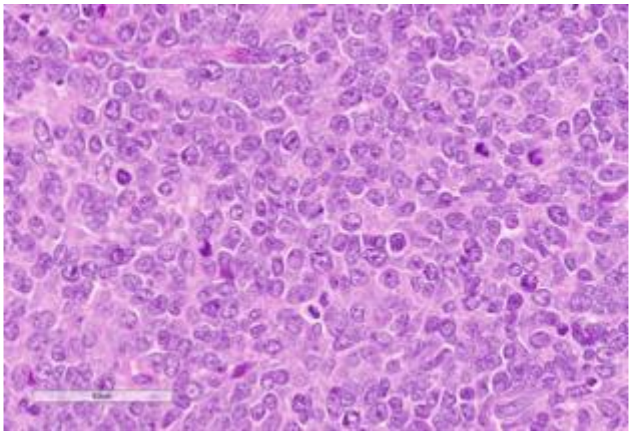

Source: MDACC

**Clinical Information**

Originating Biopsy Diagnosis: DLBCL, NOS

Prior Indolent Lymphoma: None

Sex: Male

Race: White or Caucasian

Ethnicity: Not Hispanic or Latino

Age at diagnosis: 53

Age at PDX biopsy: 61

### LoTs Pre Biopsy (best response): 2 (SD)

### LoTs Post Biopsy (best response): 3 (SD)

Therapies: Chemotherapy; Chemotherapy; Chemotherapy; Chemotherapy; CART

**Immunohistochemical Staining:**

Hans COO call: GCB

BCL6 (+)

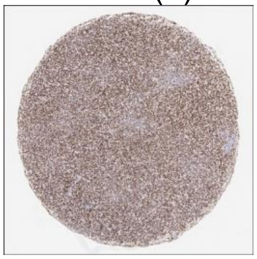

CD10 (+)

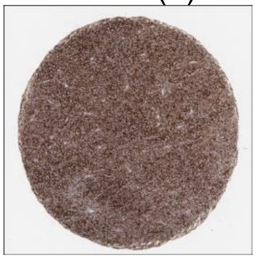

MUM1 (-)

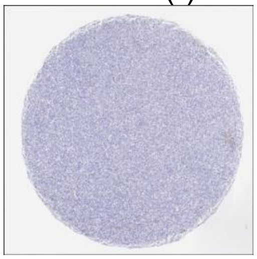

MYC (-)

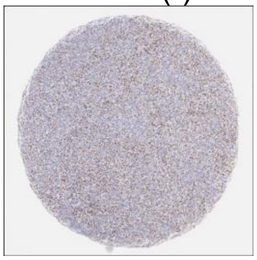

CD20 (+)

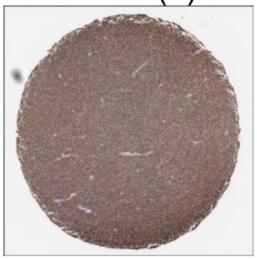

CD19 (+)

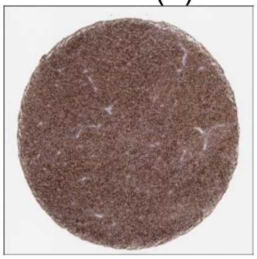

**DLBCL90 call:** GCB, DZsig+

**Genetic Characteristics:**

| | | |
|---|---|---|
| BCL2 FISH: Positive | MYC FISH: Negative | BCL6 FISH: Negative |

Copy Number Alterations:

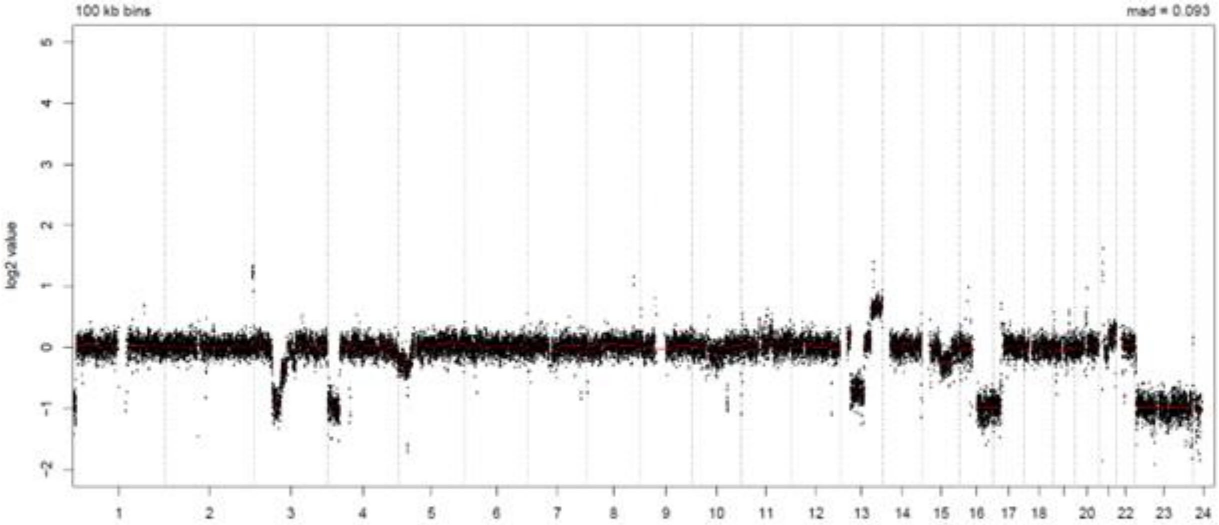


LymphGen Class (Confidence): EZB

| | | |
|---|---|---|
| BN2: 0.01 | EZB: 0.99 | N1: 0.07 |
| MCD: 0.00 | ST2: 0.01 | A53: 0.00 |

### LPMI556

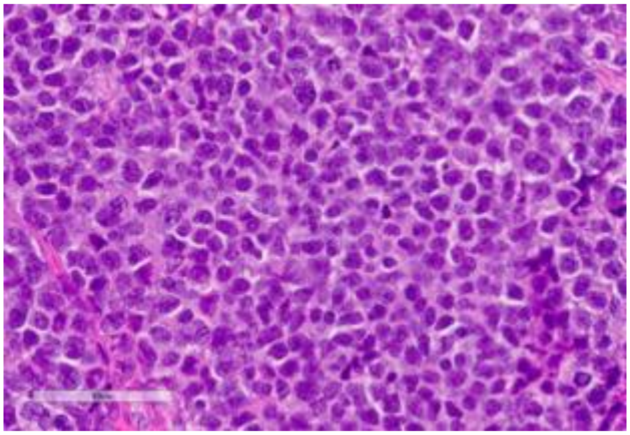

Source: MDACC

**Clinical Information**

Originating Biopsy Diagnosis: HGBCL

Prior Indolent Lymphoma: Follicular Lymphoma

Sex: Male

Race: White or Caucasian

Ethnicity: Not Hispanic or Latino

Age at diagnosis: 67

Age at PDX biopsy: 82

### LoTs Pre Biopsy (best response): 1 (CR)

### LoTs Post Biopsy (best response): 2 (CR)

Therapies: Targeted; Chemotherapy; Targeted

**Immunohistochemical Staining:**

Hans COO call: GCB

BCL6 (+)

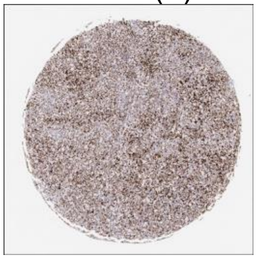

CD10 (+)

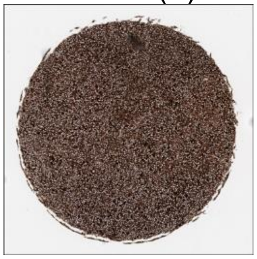

MUM1 (-)

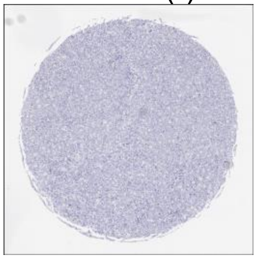

MYC (-)

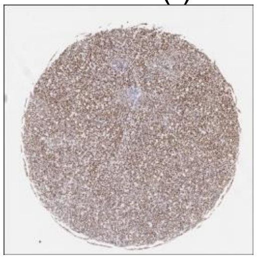

CD20 (+)

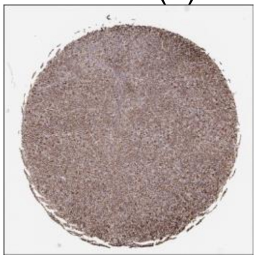

CD19 (+)

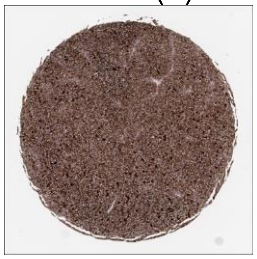

**DLBCL90 call:** GCB

**Genetic Characteristics:**

BCL2 FISH: Positive
MYC FISH: Positive
BCL6 FISH: Negative

Copy Number Alterations:

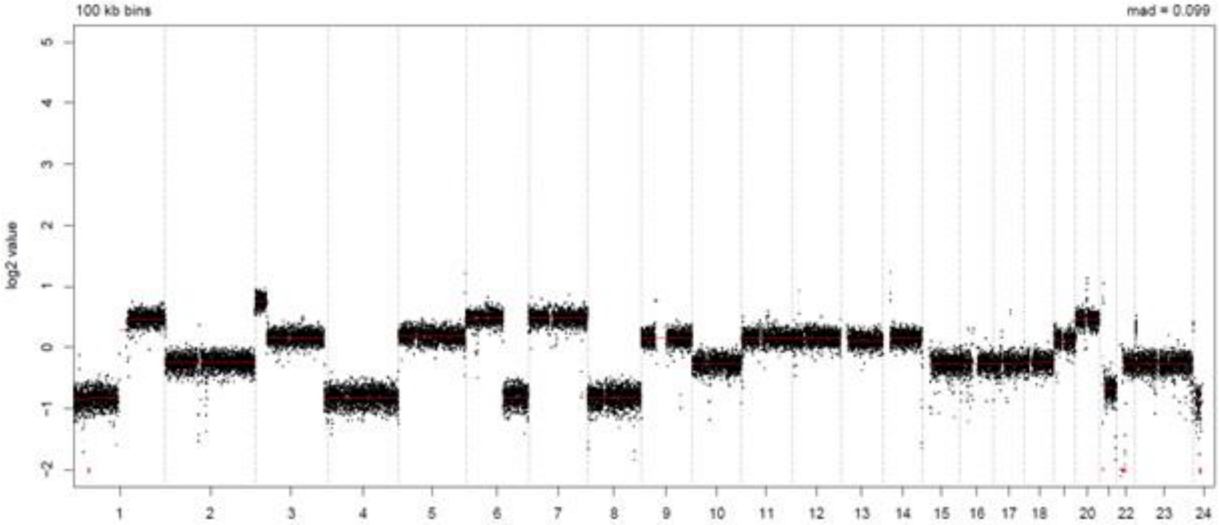


LymphGen Class (Confidence): Other

BN2: 0.01
EZB: 0.63
N1: 0.07
MCD: 0.00
ST2: 0.03
A53: 0.81

### LPMI580

Source: MDACC

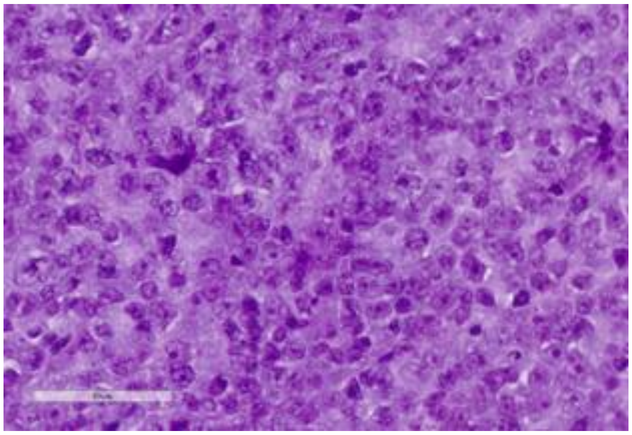

**Clinical Information**

Originating Biopsy Diagnosis: DLBCL, GCB

Prior Indolent Lymphoma: None

Sex: Male

Race: White or Caucasian

Ethnicity: Not Hispanic or Latino

Age at diagnosis: 64

Age at PDX biopsy: 64

### LoTs Pre Biopsy (best response): 0 (NA)

### LoTs Post Biopsy (best response): 1 (CR)

Therapies: Chemotherapy

**Immunohistochemical Staining:**

Hans COO call: GCB

BCL6 (+)

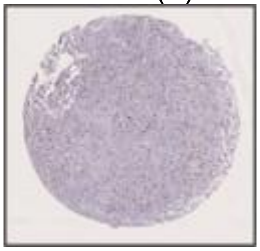

CD10 (+)

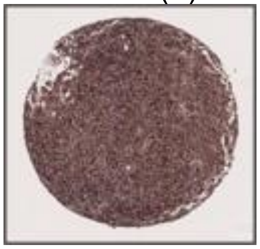

MUM1 (-)

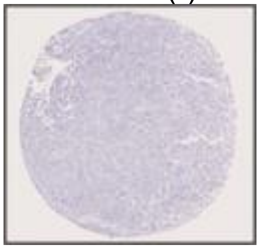

MYC (+)

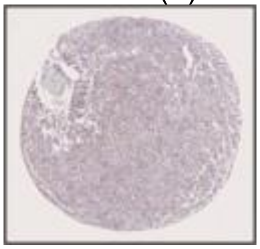

CD20 (+)

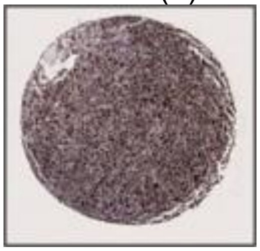

CD19 (+)

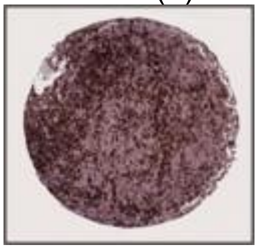

**DLBCL90 call:** GCB, DZsig+

**Genetic Characteristics:**

| | | |
|---|---|---|
| BCL2 FISH: Positive | MYC FISH: Positive | BCL6 FISH: Negative |

Copy Number Alterations:

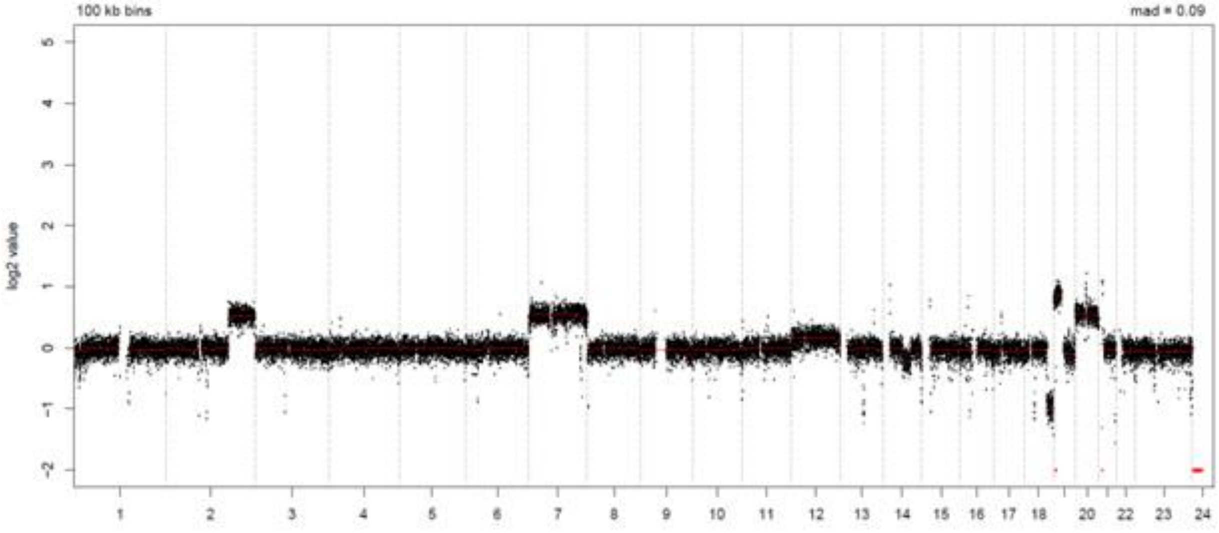


LymphGen Class (Confidence): EZB/ST2

| | | |
|---|---|---|
| BN2: 0.01 | EZB: 0.93 | N1: 0.07 |
| MCD: 0.00 | ST2: 0.95 | A53: 0.00 |

### LPMI581

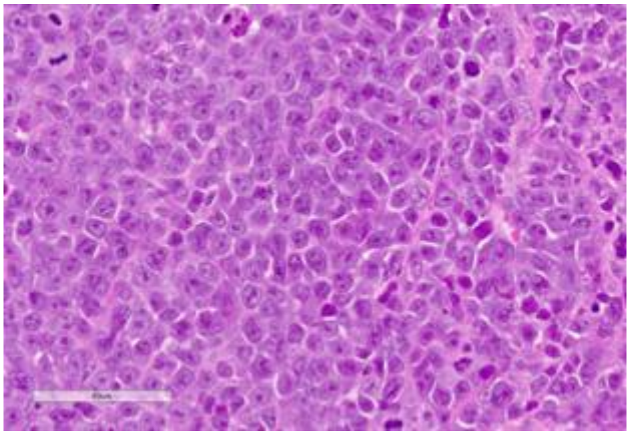

Source: MDACC

**Clinical Information**

<u>Originating Biopsy Diagnosis</u>: HGBCL

<u>Prior Indolent Lymphoma</u>: Follicular Lymphoma

<u>Sex</u>: Male

<u>Race</u>: White or Caucasian

<u>Ethnicity</u>: Not Hispanic or Latino

<u>Age at diagnosis</u>: 60

<u>Age at PDX biopsy</u>: 67

<u># LoTs Pre Biopsy</u> (best response): 1 (PD)

<u># LoTs Post Biopsy</u> (best response): 1 (PD)

<u>Therapies</u>: Chemotherapy; Targeted

**Immunohistochemical Staining:**

Hans COO call: GCB

BCL6 (-)

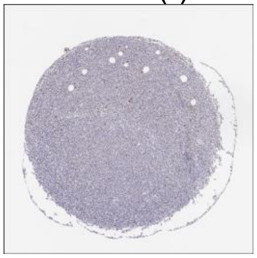

CD10 (+)

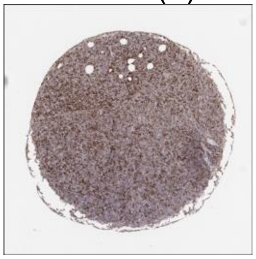

MUM1 (-)

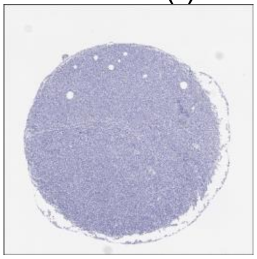

MYC (+)

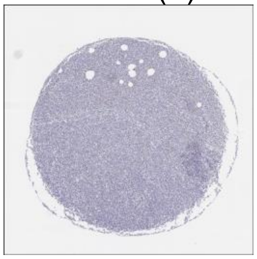

CD20 (+)

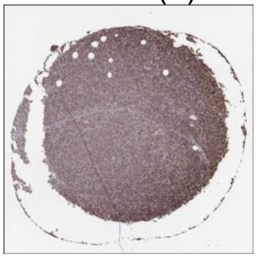

CD19 (+)

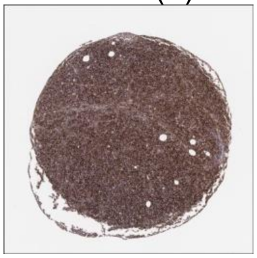

**DLBCL90 call:** GCB, DZsig+

**Genetic Characteristics:**

BCL2 FISH: Negative

MYC FISH: Positive

BCL6 FISH: Positive

Copy Number Alterations:

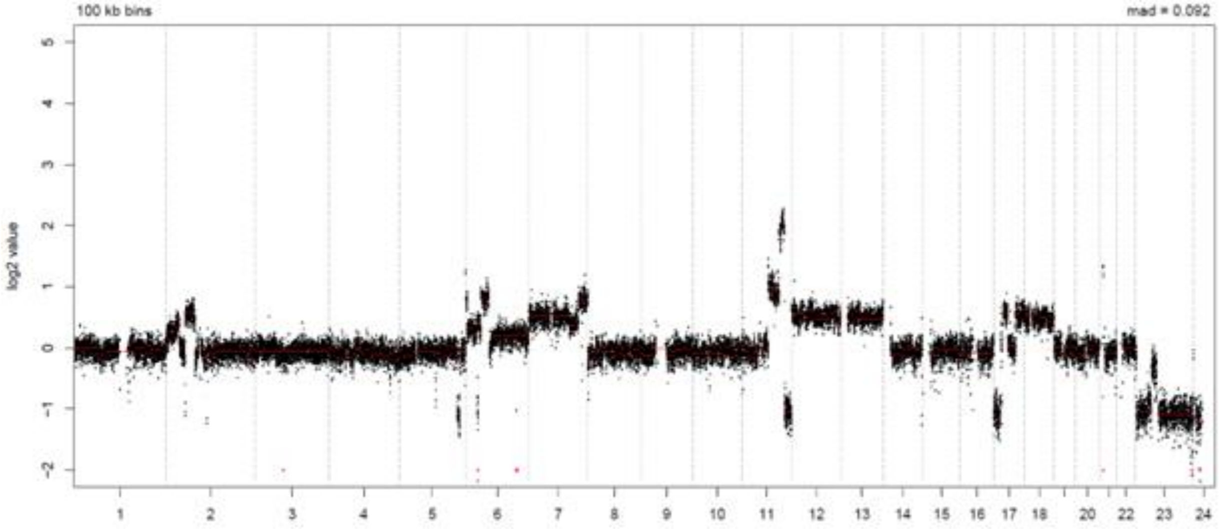


LymphGen Class (Confidence): BN2

BN2: 0.86

EZB: 0.48

N1: 0.07

MCD: 0.00

ST2: 0.00

A53: 0.00

### LPMI612

Source: MDACC

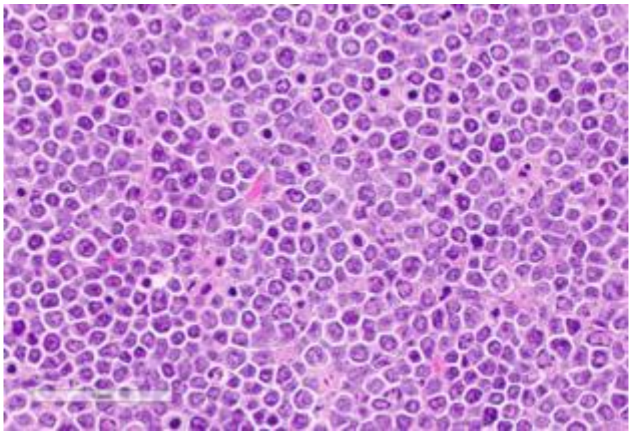

**Clinical Information**

Originating Biopsy Diagnosis: DLBCL, NOS

Prior Indolent Lymphoma: Follicular Lymphoma

Sex: Male

Race: White or Caucasian

Ethnicity: Not Hispanic or Latino

Age at diagnosis: 53

Age at PDX biopsy: 61

### LoTs Pre Biopsy (best response): 4 (PD)

### LoTs Post Biopsy (best response): 1 (CR)

Therapies: Chemotherapy; Chemotherapy; Chemotherapy; Chemotherapy; CART

**Immunohistochemical Staining:**

Hans COO call: GCB

BCL6 (-)

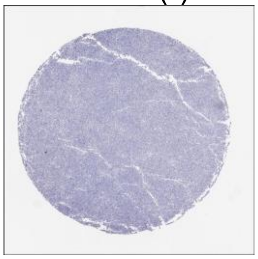

CD10 (+)

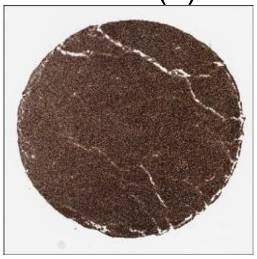

MUM1 (+)

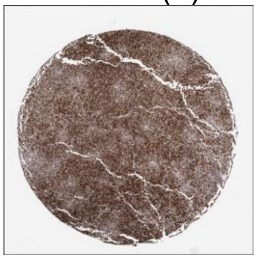

MYC (+)

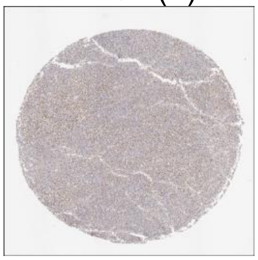

CD20 (+)

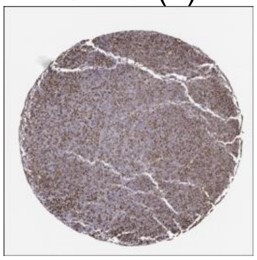

CD19 (+)

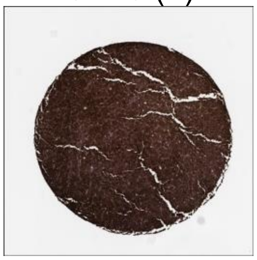

**DLBCL90 call:** GCB, DZsig+

**Genetic Characteristics:**

BCL2 FISH: Positive | MYC FISH: Positive | BCL6 FISH: Negative

Copy Number Alterations:

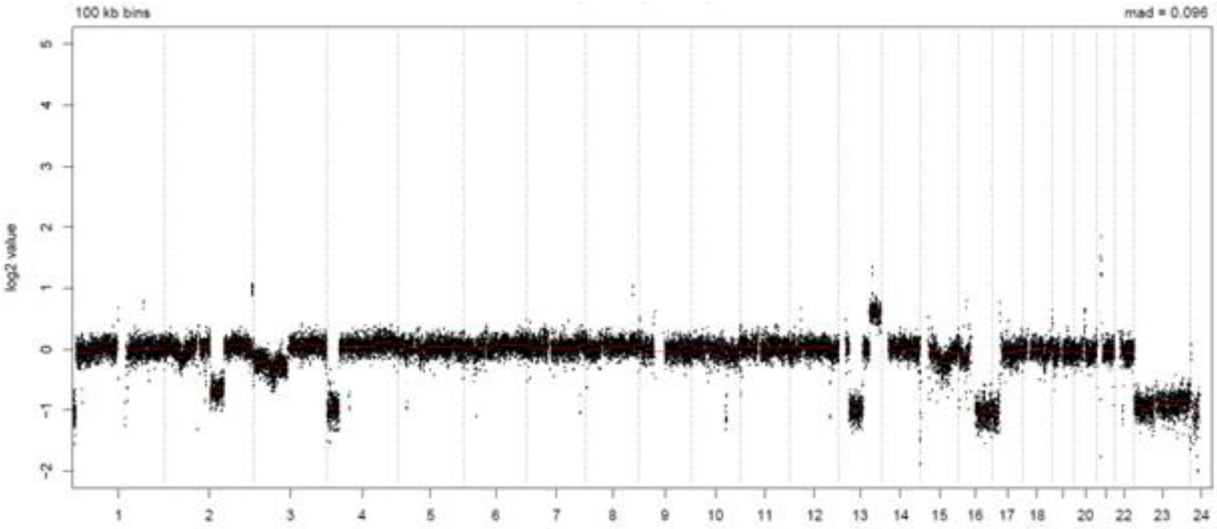


LymphGen Class (Confidence): EZB

| | | |
|---|---|---|
| BN2: 0.01 | EZB: 0.99 | N1: 0.07 |
| MCD: 0.00 | ST2: 0.06 | A53: 0.00 |

### LPMI702

Source: MDACC

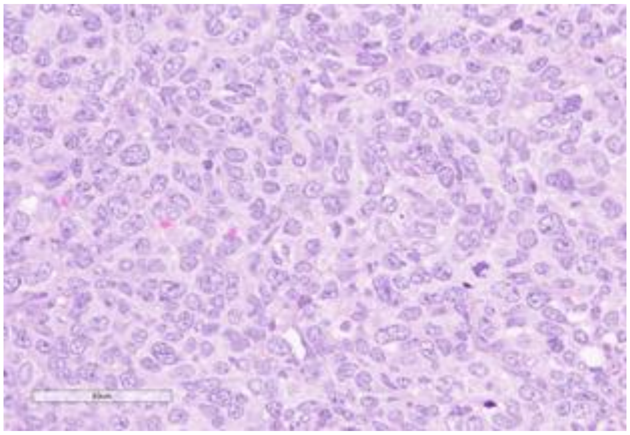

**Clinical Information**

Originating Biopsy Diagnosis: DLBCL, GCB

Prior Indolent Lymphoma: Follicular Lymphoma

Sex: Male

Race: Asian

Ethnicity: Not Hispanic or Latino

Age at diagnosis: 38

Age at PDX biopsy: 43

### LoTs Pre Biopsy (best response): 3 (PR)

### LoTs Post Biopsy (best response): 2 (PR)

Therapies: Chemotherapy; Chemotherapy; Chemotherapy; Chemotherapy; CART

**Immunohistochemical Staining:**

Hans COO call: GCB

BCL6 (+)

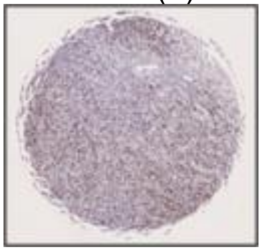

CD10 (+)

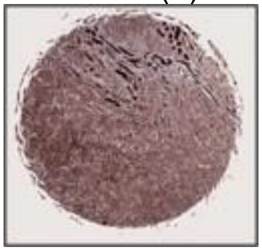

MUM1 (+)

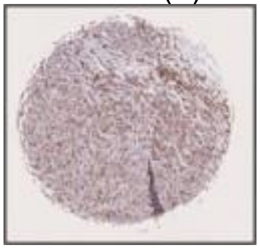

MYC (-)

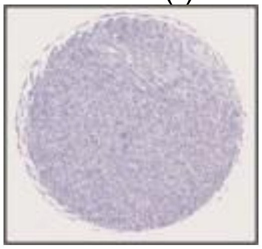

CD20 (+)

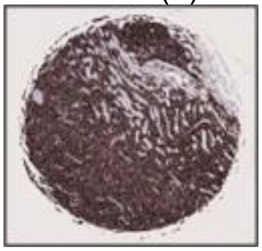

CD19 (+)

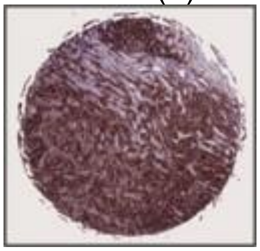

**DLBCL90 call:** GCB

**Genetic Characteristics:**

BCL2 FISH: Positive    MYC FISH: Negative    BCL6 FISH: Negative

Copy Number Alterations:

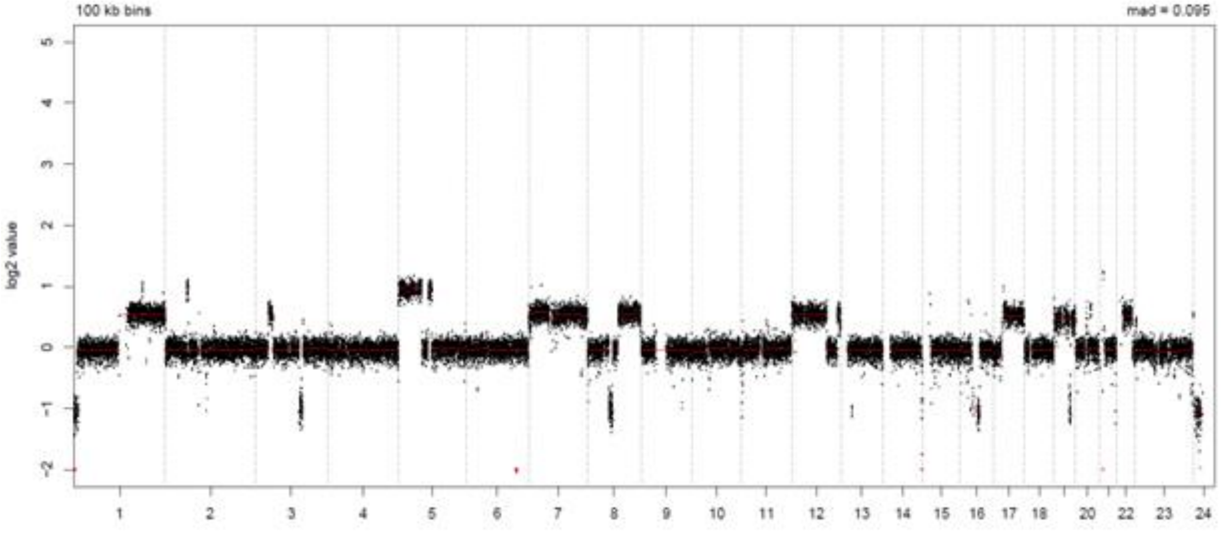


LymphGen Class (Confidence): EZB

BN2: 0.01    EZB: 0.98    N1: 0.07
MCD: 0.00    ST2: 0.27    A53: 0.00

### LPMI725

Source: MDACC

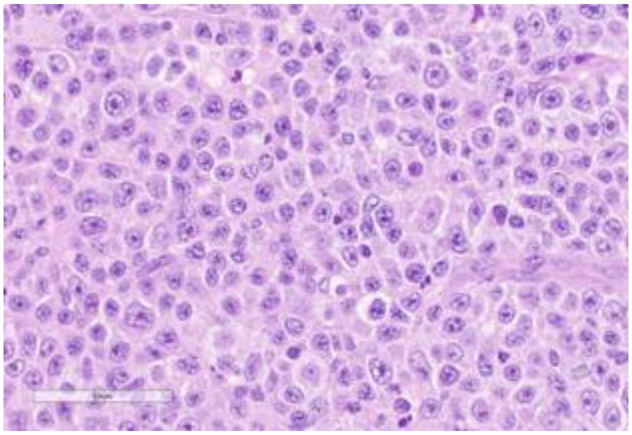

**Clinical Information**

Originating Biopsy Diagnosis: DLBCL

Prior Indolent Lymphoma: None

Sex: Male

Race: White or Caucasian

Ethnicity: Not Hispanic or Latino

Age at diagnosis: 38

Age at PDX biopsy: 40

### LoTs Pre Biopsy (best response): 6 (PD)

### LoTs Post Biopsy (best response): 5 (PD)

Therapies: Chemotherapy; Chemotherapy; Chemotherapy; Targeted; Chemotherapy; CART; Bispecific; RT; Targeted; ADC; Chemotherapy

**Immunohistochemical Staining:**

Hans COO call: GCB

BCL6 (+)

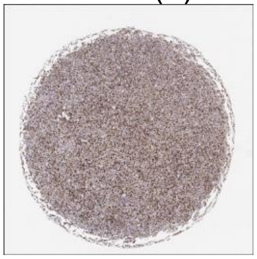

CD10 (+)

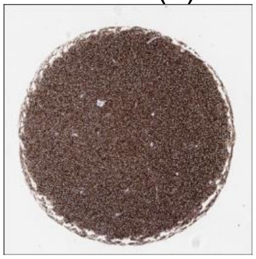

MUM1 (-)

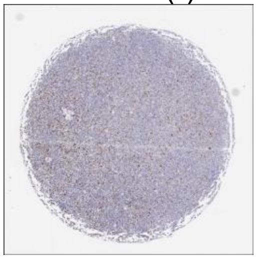

MYC (-)

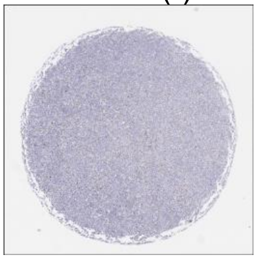

CD20 (+)

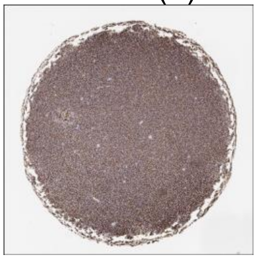

CD19 (+)

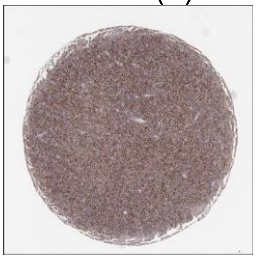

**DLBCL90 call:** GCB

**Genetic Characteristics:**

BCL2 FISH: Negative | MYC FISH: Negative | BCL6 FISH: Negative

Copy Number Alterations:

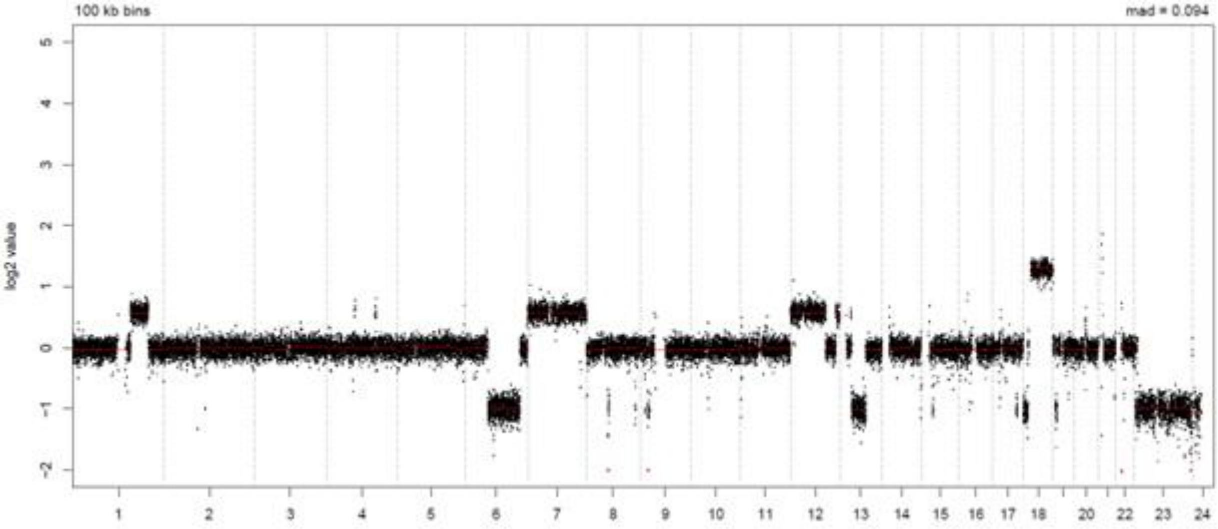


LymphGen Class (Confidence): Other

BN2: 0.01 | EZB: 0.00 | N1: 0.07
MCD: 0.00 | ST2: 0.06 | A53: 0.00

### LPMI734

Source: MDACC

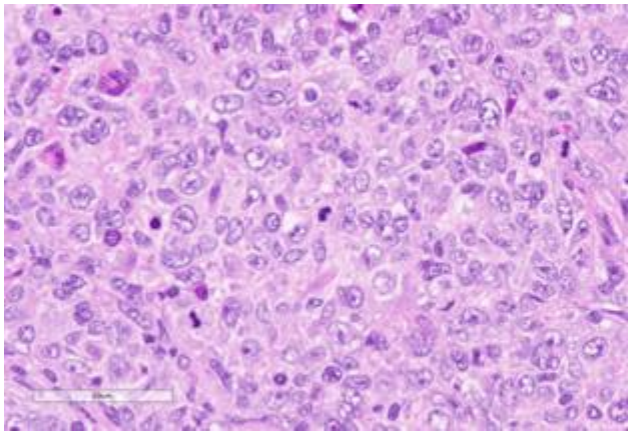

**Clinical Information**

Originating Biopsy Diagnosis: DLBCL, ABC

Prior Indolent Lymphoma: Marginal Zone Lymphoma

Sex: Female

Race: White or Caucasian

Ethnicity: Not Hispanic or Latino

Age at diagnosis: 54

Age at PDX biopsy: 55

### LoTs Pre Biopsy (best response): 2 (PR)

### LoTs Post Biopsy (best response): 7 (SD)

Therapies: Chemotherapy; Chemotherapy; Targeted; Targeted; Chemotherapy; Bispecific; Chemotherapy; Chemotherapy; CART

**Immunohistochemical Staining:**

Hans COO call: N/A

BCL6 (NA)

CD10 (NA)

MUM1 (NA)

MYC (NA)

CD20 (NA)

CD19 (NA)

The linked image cannot be displayed. The file may have been moved, renamed, or deleted. Verify that the link points to the correct file and location.

The linked image cannot be displayed. The file may have been moved, renamed, or deleted. Verify that the link points to the correct file and location.

The linked image cannot be displayed. The file may have been moved, renamed, or deleted. Verify that the link points to the correct file and location.

**DLBCL90 call:** ABC

**Genetic Characteristics:**

BCL2 FISH: N/A

MYC FISH: N/A

BCL6 FISH: N/A

Copy Number Alterations:

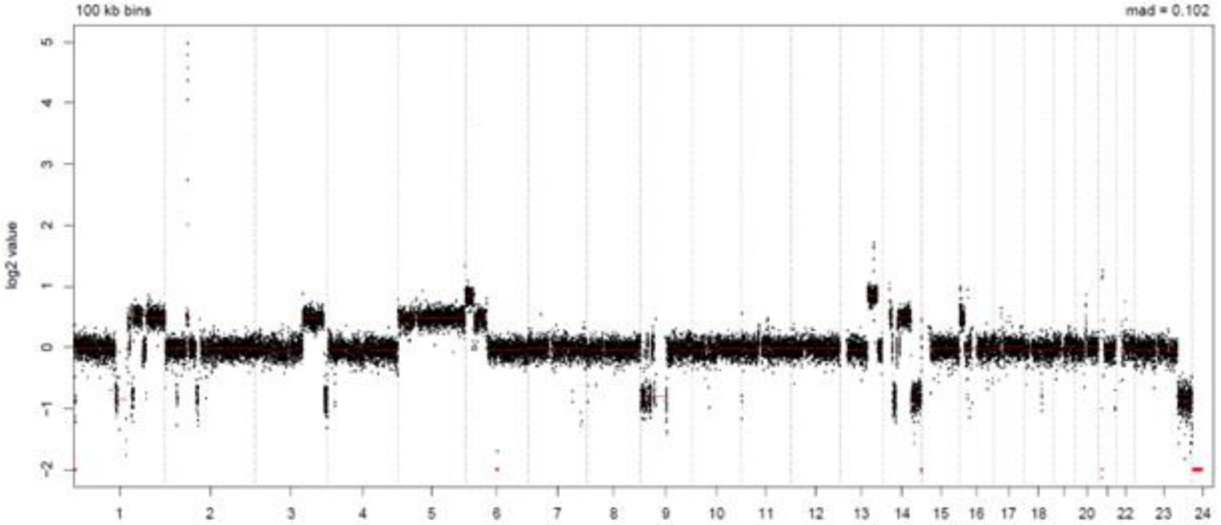


LymphGen Class (Confidence): BN2

BN2: 0.95

EZB: 0.00

N1: 0.07

MCD: 0.00

ST2: 0.57

A53: 0.00

### LPMI797

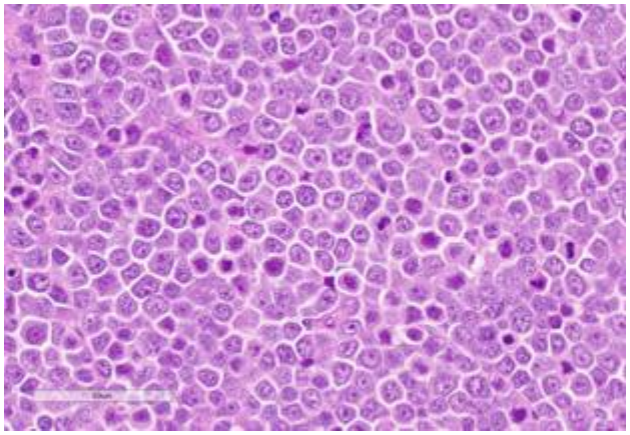

Source: MDACC

**Clinical Information**

<u>Originating Biopsy Diagnosis</u>: DLBCL, NOS

<u>Prior Indolent Lymphoma</u>: None

<u>Sex</u>: Male

<u>Race</u>: White or Caucasian

<u>Ethnicity</u>: Not Hispanic or Latino

<u>Age at diagnosis</u>: 75

<u>Age at PDX biopsy</u>: 76

<u># LoTs Pre Biopsy</u> (best response): 6 (PD)

<u># LoTs Post Biopsy</u> (best response): 2 (PD)

<u>Therapies</u>: Chemotherapy; Chemotherapy; CART; Chemotherapy; Immunotherapy; RT; Targeted; Targeted

**Immunohistochemical Staining:**

Hans COO call: GCB

BCL6 (-)

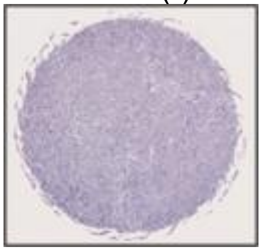

CD10 (+)

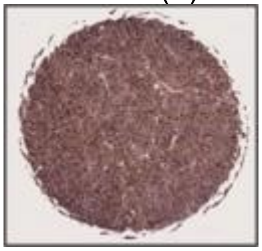

MUM1 (+)

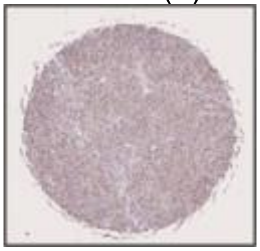

MYC (+)

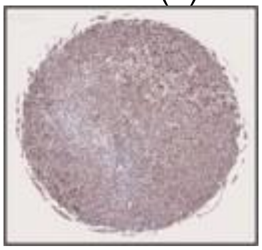

CD20 (+)

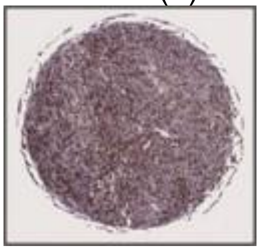

CD19 (+)

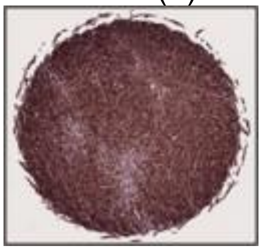

**DLBCL90 call:** ABC

**Genetic Characteristics:**

BCL2 FISH: Positive | MYC FISH: Positive | BCL6 FISH: Negative

Copy Number Alterations:

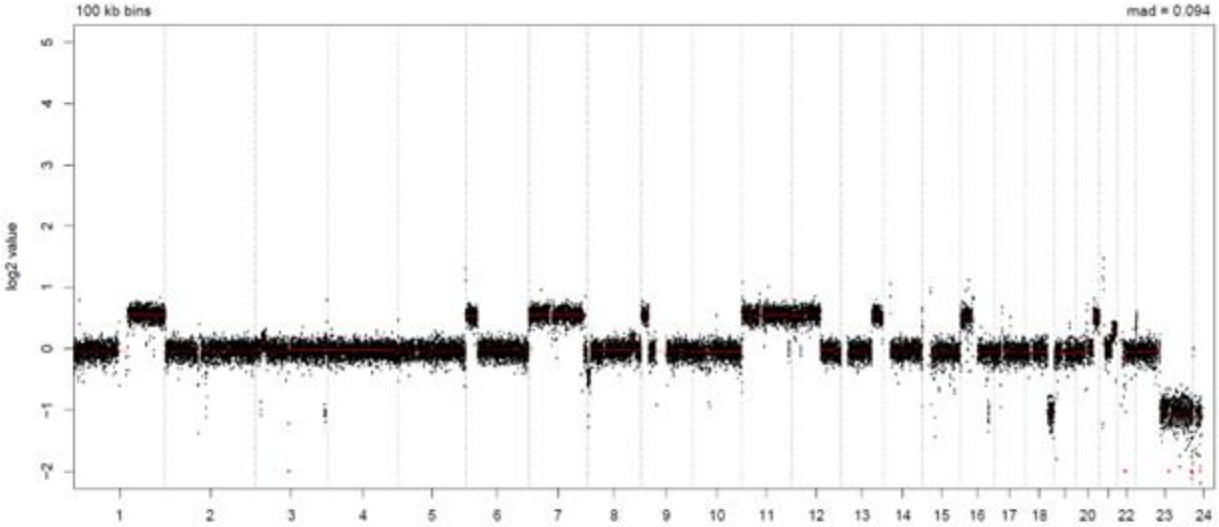


LymphGen Class (Confidence): EZB

BN2: 0.07 | EZB: 0.92 | N1: 0.07
MCD: 0.00 | ST2: 0.15 | A53: 0.00

### LPMI805

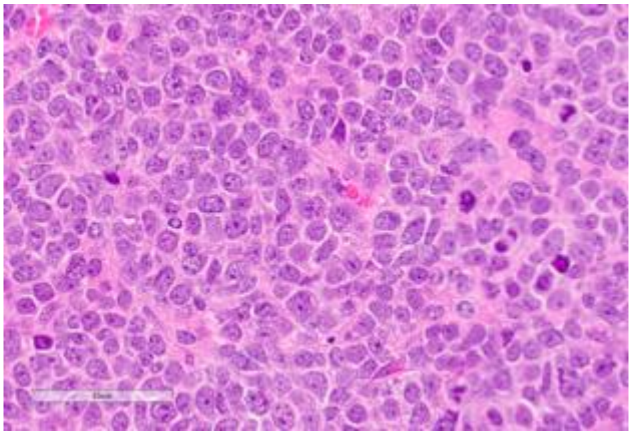

Source: MDACC

**Clinical Information**

Originating Biopsy Diagnosis: DLBCL, NOS

Prior Indolent Lymphoma: None

Sex: Male

Race: White or Caucasian

Ethnicity: Not Hispanic or Latino

Age at diagnosis: 65

Age at PDX biopsy: 65

### LoTs Pre Biopsy (best response): 0 (NA)

### LoTs Post Biopsy (best response): 1 (CR)

Therapies: Chemotherapy

**Immunohistochemical Staining:**

Hans COO call: GCB

BCL6 (+)

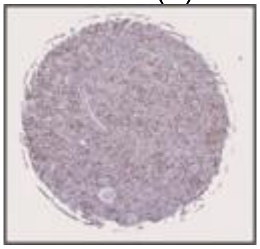

CD10 (+)

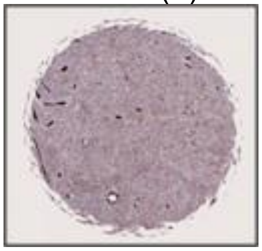

MUM1 (+)

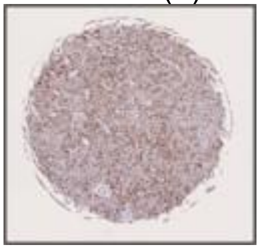

MYC (-)

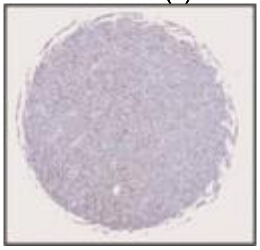

CD20 (+)

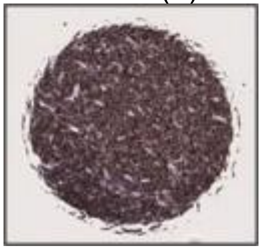

CD19 (+)

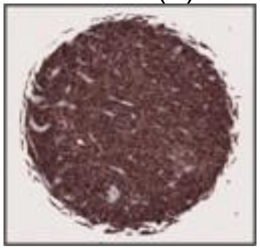

**DLBCL90 call:** UNCLASS

**Genetic Characteristics:**

BCL2 FISH: Negative | MYC FISH: Negative | BCL6 FISH: Negative

Copy Number Alterations:

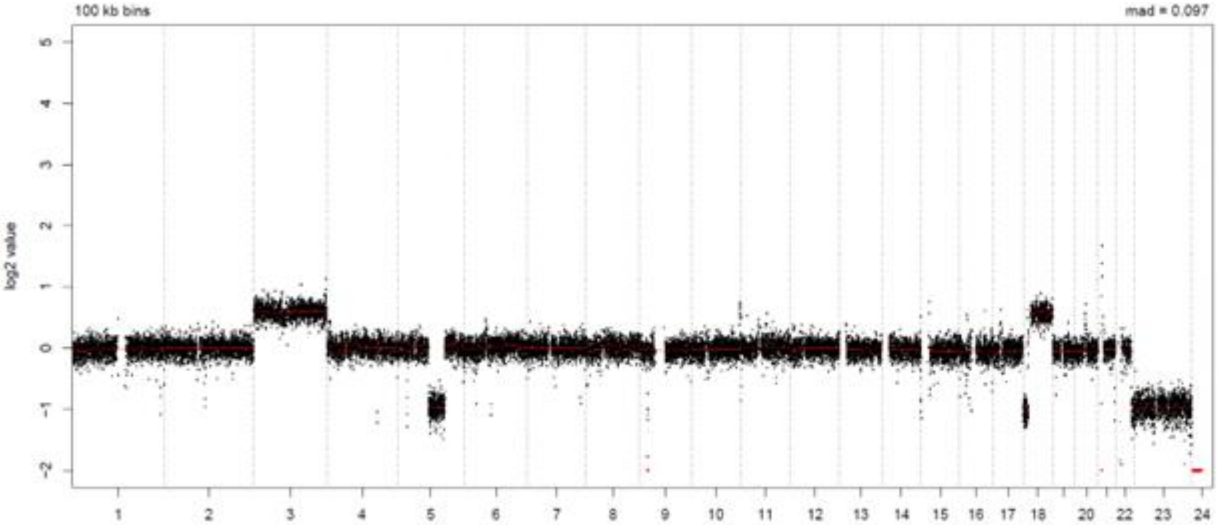


LymphGen Class (Confidence): Other

| | | |
|---|---|---|
| BN2: 0.01 | EZB: 0.12 | N1: 0.07 |
| MCD: 0.00 | ST2: 0.01 | A53: 0.00 |

### LPMI809

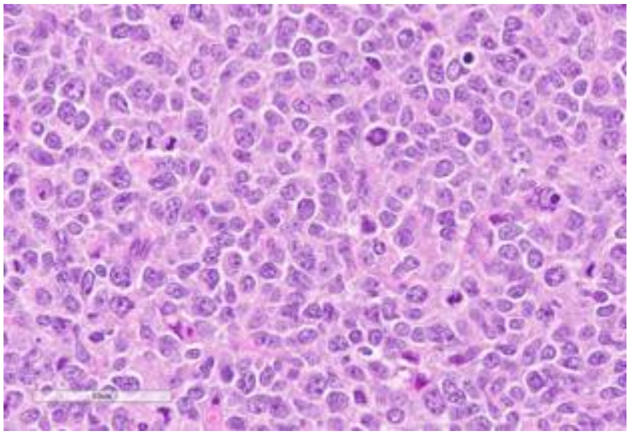

Source: MDACC

**Clinical Information**

Originating Biopsy Diagnosis: DLBCL, GCB

Prior Indolent Lymphoma: None

Sex: Female

Race: White or Caucasian

Ethnicity: Not Hispanic or Latino

Age at diagnosis: 70

Age at PDX biopsy: 70

### LoTs Pre Biopsy (best response): 0 (NA)

### LoTs Post Biopsy (best response): 1 (CR)

Therapies: Chemotherapy

**Immunohistochemical Staining:**

Hans COO call: GCB

BCL6 (+)

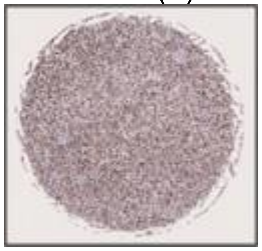

CD10 (+)

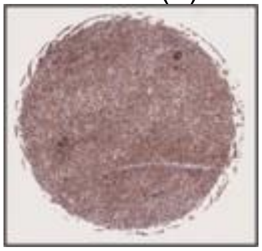

MUM1 (-)

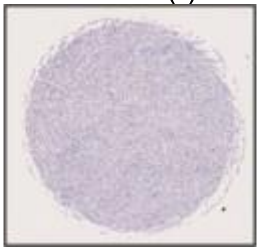

MYC (+)

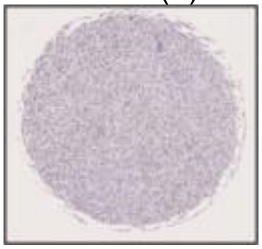

CD20 (+)

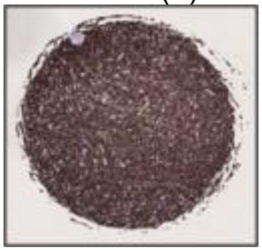

CD19 (+)

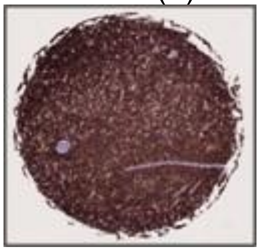

**DLBCL90 call:** GCB

**Genetic Characteristics:**

| | | |
|---|---|---|
| BCL2 FISH: Negative | MYC FISH: Negative | BCL6 FISH: Negative |

Copy Number Alterations:

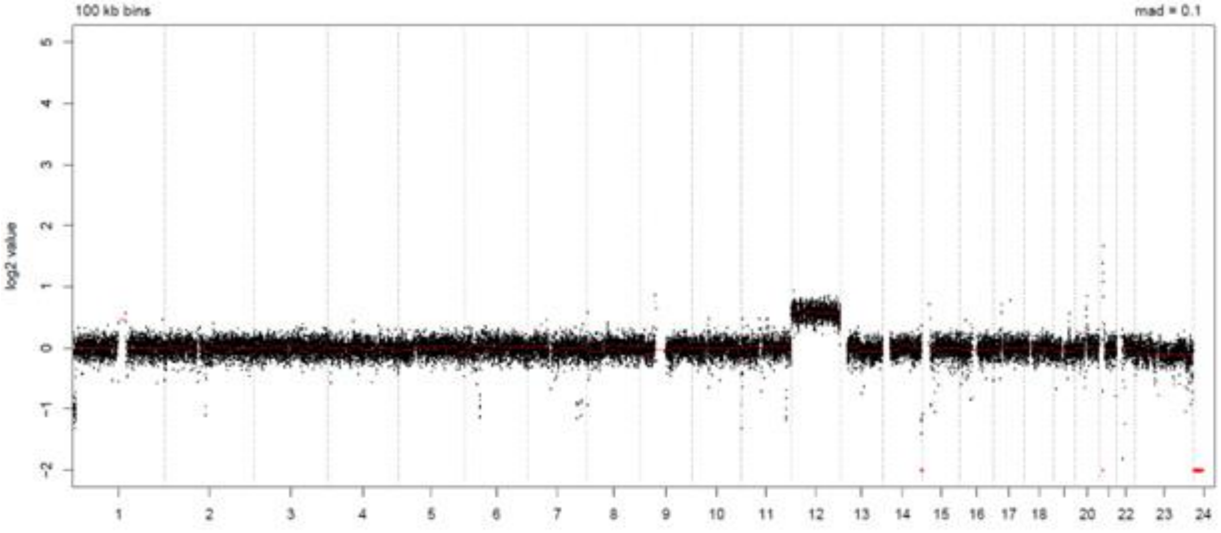


LymphGen Class (Confidence): ST2

| | | |
|---|---|---|
| BN2: 0.25 | EZB: 0.04 | N1: 0.07 |
| MCD: 0.00 | ST2: 0.95 | A53: 0.00 |

### LPMI815

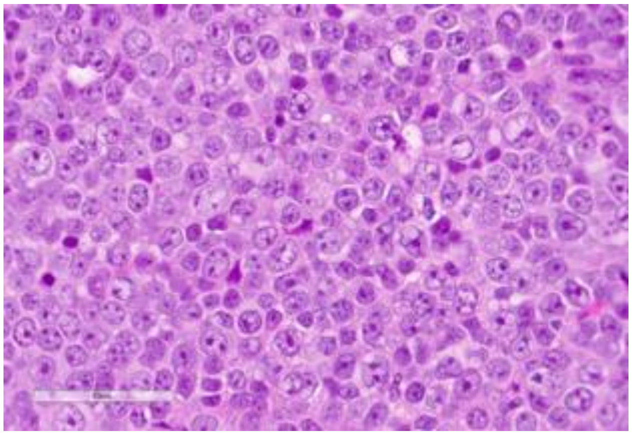

Source: MDACC

**Clinical Information**

Originating Biopsy Diagnosis: DLBCL, NOS

Prior Indolent Lymphoma: None

Sex: Male

Race: White or Caucasian

Ethnicity: Not Hispanic or Latino

Age at diagnosis: 76

Age at PDX biopsy: 76

### LoTs Pre Biopsy (best response): 1 (CR)

### LoTs Post Biopsy (best response): 2 (PD)

Therapies: Chemotherapy; CART; Chemotherapy

**Immunohistochemical Staining:**

Hans COO call: GCB

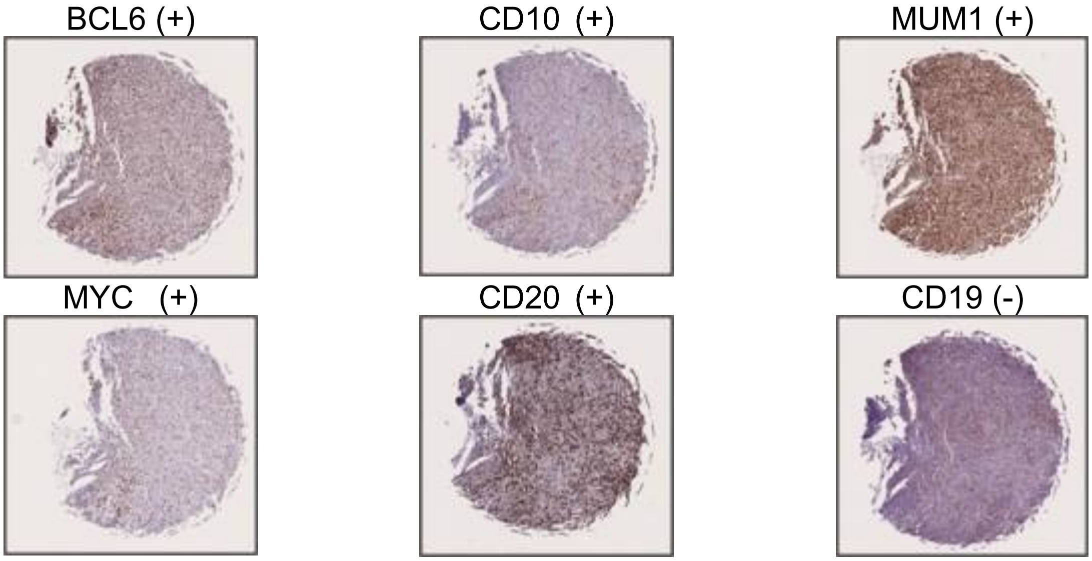

**DLBCL90 call:** ABC

**Genetic Characteristics:**

BCL2 FISH: Negative
MYC FISH: Negative
BCL6 FISH: Negative

Copy Number Alterations:

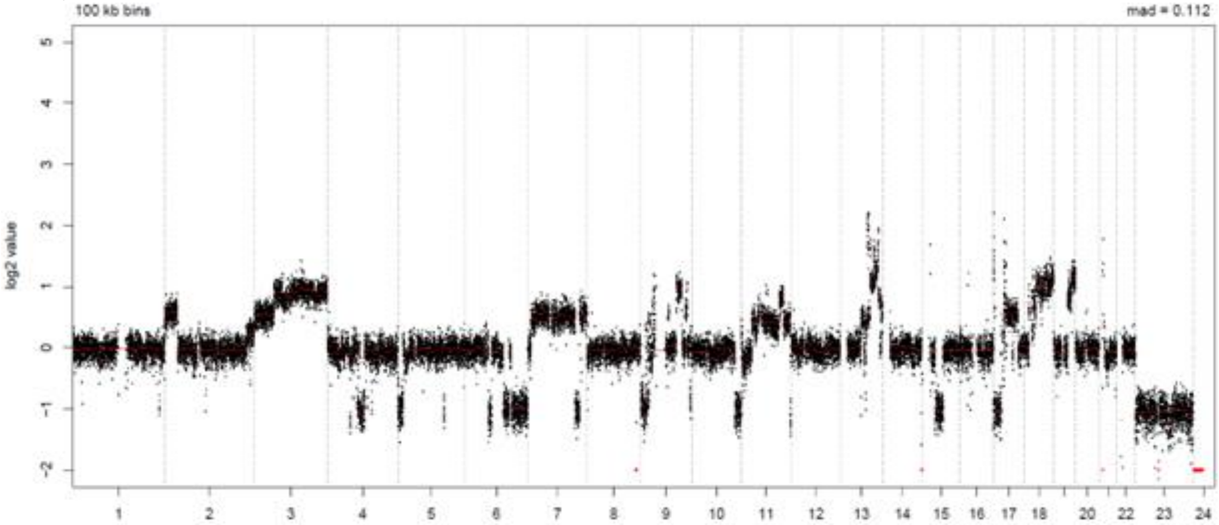


LymphGen Class (Confidence): Other

BN2: 0.01
EZB: 0.00
N1: 0.07
MCD: 0.00
ST2: 0.00
A53: 0.00
